# Can you trust your reconstructed lineage tree? A homoplasy-based approach for irreversible evolution

**DOI:** 10.1101/2025.07.27.667007

**Authors:** Pini Zilber, Sebastian Prillo, Yaara Neumeier, Nir Yosef, Boaz Nadler

**Affiliations:** Department of Computer Science and Applied Mathematics, Weizmann Institute of Science, Israel; Department of Electrical Engineering and Computer Sciences, University of California, Berkeley, USA; Department of Systems Immunology, Weizmann Institute of Science, Israel

**Author notes:** Correspondence to be sent to: Boaz Nadler,; Nir Yosef,.

**Keywords:** CRISPR-Cas9 lineage tracing, homoplasy, phylogeny, irreversible evolution, accuracy evaluation, lineage tree reconstruction

## Abstract

Phylogeny inference is a fundamental problem in computational biology, with many proposed algorithms. Emerging techniques that couple single-cell genomics with Cas9-based genome editing open the way for in-depth analysis of cell phylogenies that underlie processes of clonal expansion, selection and diversification, from embryogenesis to cancer. A key distinguishing feature of cell lineage analysis with these techniques is the non-modifiability of Cas9-induced mutations, which motivates revisiting questions in phylogenetics. In this work, we ask one such fundamental question: is it possible to assess the reliability of an inferred lineage tree, even though we do not know its underlying ground truth? We present a homoplasy-based approach for this question that leverages the non-modifiability property. We show via simulations that under a broad range of settings, our method can effectively distinguish accurate reconstructions out of a pool of candidate solutions. Importantly, our homoplasy-based score is substantially more powerful than the commonly used parsimony score - a result that we back by both empirical and theoretical analysis. The computation of the homoplasy score is simple and scalable, thus opening the way for more rigorous analysis of cell lineages.

## 1 Introduction

Reconstruction of phylogenies has long stood as a cornerstone of evolutionary biology, with numerous methods developed over the years (Hennig, 1999; Nei and Kumar, 2000; Hall, 2004; Wiley and Lieberman, 2011). In the classical setting, the objective is to reconstruct the tree of life. The leaves of the tree represent extant species, and the internal nodes provide a way to reason about their latent ancestors far back in evolutionary history. Phylogenetics has also been used to formalize the analysis of sequences of nucleic or amino acids and shed light on the evolutionary forces that shaped them. The process of natural evolution of sequences, captured by classical models such as Jukes-Cantor, reflects the chances of mutations to occur and is assumed to be *modifiable*. Namely, the occurrence of a mutation in a site does not necessarily nullify the chances of that site being mutated again.

The absence of this property is one of the major distinguishing features of an emerging branch of phylogenetics, where trees depict cell lineages, with leaves corresponding to individual sampled cells, and internal nodes reflecting their ancestral cell division relationships (Gong et al., 2021). While cell lineages can be inferred based on somatic mutations or other natural and modifiable forms of heritable variation, the most powerful techniques (in terms of numbers of cells and the resolution of lineage trees) are based on inducible mutations at synthetic DNA elements (Kester and Van Oudenaarden, 2018; McKenna and Gagnon, 2019; Yang et al., 2022). In these applications, the heritable information that guides the retrospective inference of lineages is accrued through genome editing (e.g. with CRISPR/Cas9) of synthetic “recorder sites” that are integrated into the genome (usually at the 3’ end of synthetic genes). Since CRISPR/Cas9 often has a much lower affinity to the edited sites, it is highly unlikely for additional mutations to occur in already-mutated sites, making this process *non-modifiable* (Sashittal et al., 2023).

Several studies investigated this new setting and proposed suitable algorithms to reconstruct cell lineages based on the mutation profiles accumulated in each cell (Jones et al., 2020; Feng et al., 2021; Seidel and Stadler, 2022; Wang et al., 2023; Sashittal et al., 2023), see also Gong et al. (2021) for a broad performance comparison. One strategy relies on the long-studied Camin-Sokal model (Camin and Sokal, 1965). This model assumes the less restrictive scenario of *irreversibility*, which permits mutated sites to undergo further mutations, but not to revert to an ancestral state. This is a notable difference from non-modifiability, except for the special case of a two-state evolution, where the two become identical. We note that in practical applications, other algorithms, including those designed for the classical problem such as neighbor-joining (Saitou and Nei, 1987), are often effective (Sugino and Lee, 2019; Jones et al., 2020; Prillo et al., 2025).

This multitude of strategies and algorithms for the inference problem raises the following question: given several reconstructions of the cell lineage tree that are all based on the same (non-modifiable) data, which reconstruction is more accurate? (i.e. more similar to the true one). More fundamentally, can we assess whether any individual reconstructed topology is accurate or not? This latter objective extends beyond simply ranking candidate topologies since it also aims to determine if any of the candidate topologies, e.g. the top-ranked ones, are close enough to the true topology and can be used for downstream analysis.

A common strategy to evaluate candidate solutions assumes that the true topology is known and compares directly to it (for example, using the Robinson-Foulds distance or the proportion of correct triplets (Jones et al., 2020; Gong et al., 2021)). While this strategy is naturally restricted, other approaches have been proposed that do not assume knowledge of the ground truth, including comparison of parsimony (the minimal number of ancestral mutation events that explain the data), or likelihood (given an evolutionary model of mutation accrual). These approaches, however, are often used only for ranking solutions and not for making “absolute” decisions of whether or not a given tree is similar to the latent ground truth. Moreover, due to the limited sequence length in cell lineage tracing experiments, parsimony and likelihood scores may be poor indicators of topological accuracy. Indeed, given typical lineage tracing data, the reconstructed tree after contracting mutationless branches typically contain polytomies. Refining them may produce a collection of binary trees all with the same parsimony score. The phenomenon of multiple distinct trees with identical quality is known as phylogenetic terraces (Sanderson, McMahon, and Steel, 2011; Sanderson, McMahon, Stamatakis, et al., 2015). From a broader perspective, there has been significant efforts on assessing the robustness and reliability of reconstructed trees and their parameters. Particular emphasis has been made on branch support, deriving bootstrap or Bayesian measures for the reliability of specific clades in the reconstructed tree, see Simon (2022) for a review. For non-modifiable cell lineage tree reconstruction, Seidel and Stadler (2022) and Zwaans et al. (2025) developed TiDeTree, a Bayesian framework to estimate both the tree as well as various global parameters, such as mutation rates and transition probabilities. Given a collection of multiple reconstructed trees, a different approach to possibly construct a more accurate tree and to assess the branch support is to consider a consensus tree and compare the various trees to it. For non-modifiable cell lineage tracing, both Gong et al. (2021) and Dai and Molloy (2026) have shown that consensus trees may be more accurate. Finally, another approach to assess the relative reliability of different tree reconstructions is to use external additional knowledge such as gene expression or migration data, see for example Sashittal et al. (2023). Yet, despite this extensive body of work, we are not aware of methods that can effectively distinguish accurate from inaccurate tree reconstructions.

In this work, we present the *pairwise homoplasy score (PHS)* - a simple and theoretically grounded approach to address these problems. Leveraging the non-modifiable nature of the evolution process, our algorithm calculates a sequence of tail probabilities - one for each pair of leaves in the reconstructed tree - and combines them into a single score. We demonstrate that compared to parsimony and likelihood, the PHS approach is substantially more powerful in distinguishing accurate tree reconstructions out of a pool of candidate solutions. Furthermore, we present a theoretical analysis that provides insight into the advantage of PHS over parsimony. The procedure for calculating the PHS has polynomial-time complexity, and can be easily run on trees with thousands of leaves.^1^ We expect it to become an important part of future pipelines for phylogenetic analysis in non-modifiable systems, and particularly CRISPR/Cas9-based tracing of cell lineages.

## 2 Problem Setup

We first briefly describe a typical cell lineage tracing experiment, and then the mathematical model and assumptions made in our manuscript. A typical experiment starts with a single progenitor cell that has *k* target sites along its genome, which are all unedited (i.e., yet unmutated). As the experiment ensues, the resulting progeny grows through cell divisions, with possible divergence from neutral growth due to advantageous properties of some clades. This evolutionary process can be described by a binary tree, where nodes correspond to cells. Each node *u* has a birth time *τ*_*u*_ and a sequence *s*^(*u*)^ of *k* characters, which are the Cas9-induced mutations at its *k* sites. The root node of the tree, denoted by *r*, corresponds to the progenitor cell. It has a birth time *τ*_*r*_ = 0 and an unmutated sequence *s*^*r*^ = (0, 0, …, 0). At the end of the experiment, the sequences at a subset of *n* cells are observed. Without loss of generality, we rescale time so the end of the experiment is at *τ* = 1. The *n* observed cells induce an underlying ground truth binary tree, denoted *T*_GT_, which describes their clonal history. For future use, we denote the set of sequences at the *n* cells. By definition, *S*_*n*_ is the subset of *S*_GT_ that corresponds to the terminal leaves of *T*_GT_. Finally, in the tree *T*_GT_, the path length *τ*_*u,v*_ between a node *u* and a descendant node *v* is defined as the difference between their birth times, *τ*_*u,v*_ = *τ*_*v*_ −*τ*_*u*_.

In the main text, we assume for simplicity that the sequences at the *n* sampled cells are fully observed, namely, all entries of the matrix *S*_*n*_ are observed. Settings with missing data, due to either stochastic or heritable missingness, are addressed in Appendix B. Given the *n* observed sequences *S*_*n*_, standard problems are to reconstruct either the underlying tree topology *T*_GT_ or the full tree *T*_GT_*S*_GT_. In contrast, we focus on assessing whether a given reconstructed tree is accurate and close to *T*_GT_. To this end, in the next subsection we describe a commonly assumed model for CRISPR-Cas9 cell lineage data, and in Section 2.2 we mathematically formulate our problem.

### 2.1 CRISPR-Cas9 non-modifiable mutation model

We consider the following probabilistic model for a CRISPR-Cas9 cell lineage tracing experiment, similar to Wang et al. (2023), Sashittal et al. (2023), and Dai and Molloy (2026), among others. The first assumption is that mutations at different target sites occur independently along the tree. Specifically, each unmutated character *i* ∈ [*k*] (namely with *s*_*i*_ = 0) evolves according to a Poisson process with a mutation rate *λ*. For simplicity, this rate *λ* is assumed fixed for all character locations in the tree; some extensions are discussed in Section 3. Once a mutation occurs, the mutated state is drawn from a probability distribution 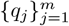 over a set of *m* possible integer states 1, …, *m*, where *m* = 1 corresponds to the binary case. The evolution is *non-modifiable*: once a character is mutated, i.e. it has a nonzero state *s*_*i*_ > 0, it cannot mutate again and its state remains fixed. Furthermore, all mutations of a cell are passed on to all its descendants.

In practice, this property is a design feature of CRISPR-Cas9 lineage recorders. An indel at a target site disrupts the sequence recognized by the guide RNA, so the site is not re-cut and the edit is not reverted, and is inherited unchanged by all descendant cells (Yang et al., 2022; Chan et al., 2019).

There are two key quantities that characterize the non-modifiable process described above. One is the probability *ρ* that at the end of the experiment an observed character is mutated. It is given by

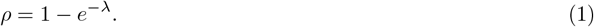

The second quantity is the collision probability, denoted by *q*. This is the probability that two independent mutation events result in the same mutated state, and it is given by

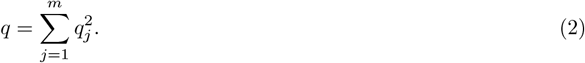

In general, the collision probability satisfies *q* ≤ 1, with *q* = 1 if and only if states are binary (*m* = 1).

The parameters characterizing the tree topology and the generative process for the sequences are summarized in Table 1. We conclude this subsection with the following definition that will be used extensively in the manuscript.

**Table 1:** Model parameters.

|  |  |
| --- | --- |
| number of leaves | $n$ |
| sequence length | $k$ |
| number of mutated states | $m$ |
| mutation probability to state $j$ | $q_j$ |
| collision probability | $q \equiv \sum_{j=1}^m q_j^2$ |
| mutation rate | $\lambda$ |
| character mutation probability at a leaf | $\rho \equiv 1 - e^{-\lambda}$ |

#### Definition 1

(full tree). A full tree *T S* is a structure that consists of both a tree topology *T*, all of its branch lengths (elapsed time between divisions), and a collection of the character sequences *S* at all its tree nodes.

### 2.2 Distinguishing between accurate and inaccurate reconstructed trees

As mentioned in the introduction, several reconstruction algorithms were developed for non-modifiable CRISPR-Cas9 evolution models, for example Jones et al. (2020), Feng et al. (2021), Seidel and Stadler (2022), Wang et al. (2023), and Sashittal et al. (2023). The celebrated Neighbor-Joining method (Saitou and Nei, 1987), originally developed for reversible evolution models, was shown to perform well also in non-modifiable settings (Sugino and Lee, 2019; Jones et al., 2020; Prillo et al., 2025).

Given the input sequences *S*_*n*_, with a limited sequence length *k*, it may be impossible to exactly reconstruct the ground-truth tree. Different reconstruction algorithms often output different tree topologies. Some of these reconstructed trees may be accurate, whereas others may be quite distant from the ground-truth tree. The problem at the focus of our work is to develop a method to decide whether a given reconstructed tree is accurate, or far from the ground-truth tree. Crucially, this task needs to be accomplished with the ground-truth tree being unknown. Our method exploits the non-modifiability of CRISPR-Cas9 mutations to calculate the probability of shared mutations across a lineage (homoplasies), thereby providing a robust statistical test for the accuracy of reconstructed trees.

In the phylogenetics literature, several works considered a different, though related problem of proposing meaningful distances between a reconstructed tree *T* and a known ground truth *T*_GT_, such as the Robinson-Foulds (RF) distance (Robinson and Foulds, 1981) and the triplets score (Jones et al., 2020). These measures are useful in simulation studies, but not for assessing tree reconstructions in practical settings, where *T*_GT_ is unknown.

Two common measures that do not require knowledge of *T*_GT_ are the parsimony and the likelihood of a tree. In the parsimony approach, a simple polynomial-time procedure is applied to a reconstructed topology *T* to find the minimal number of ancestral mutations that give rise to the observed leaf states (Gusfield, 1997); see Appendix C. In contrast, the likelihood approach considers all possible combinations of ancestral states. Under a Markov assumption, this score can be calculated efficiently, while further requiring a model of mutation accrual and knowledge of edge lengths. According to the maximum parsimony principle, trees with a smaller number of mutations *M* (*TS*) are considered closer to the ground-truth one (Fitch, 1971; Albert, 2005). Similarly, trees with higher likelihood values are considered more accurate. Likelihood is more statistically grounded than parsimony: Under the assumption of a generative model of evolution, trees with maximum likelihood *L*(*TS*) are statistically consistent (Felsenstein, 1981; Warnow, 2018).

In principle, a set of reconstructed trees can be ordered by their parsimony or likelihood scores. This ordering is useful if the goal is to evaluate how different methods successfully optimized a given parsimony- or likelihood-based objective function. However, success in optimization does not always translate to meaningful differences in topological accuracy (e.g., as measured by RF distance). Furthermore, even if we evaluate trees solely by parsimony or likelihood rather than by topological accuracy, relying on their induced ordering can be misleading. As these scores do not provide an absolute measure of quality, ordering the reconstructed trees may be meaningless if their parsimony or likelihood scores are all very far from that of the ground truth. We empirically illustrate these issues in Section 4.

A fundamental challenge is thus to devise a score that is able to detect if a given tree is accurate or not, for example indicate if its normalized RF distance from the unknown ground-truth tree is smaller than some prescribed value *ϵ*. We mathematically formulate this challenge as follows: Given a tree *T S*, a distance measure *d* and a threshold *ϵ* ∈ (0, 1), we consider the following hypothesis testing problem,

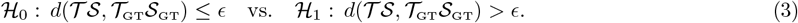

We emphasize that the main obstacle is that *T*_GT_ *S*_GT_, and thus *d*(*TS, T*_GT_ *S*_GT_), are unknown. Note that in (3), the null hypothesis is the desirable one, in which the candidate tree is close to the ground-truth one. In this paper, we consider the following four distance functions:

i. the normalized RF distance

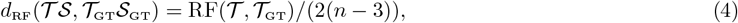
ii. the triplets distance

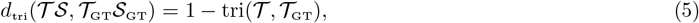
iii. a parsimony-based distance

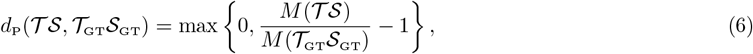

and (iv) a likelihood-based distance,

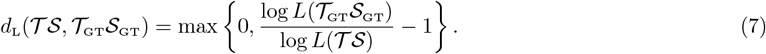

The max operator in Eqs. (6) and (7) ensures that these distances are non-negative. This is required as, given ob-served data _*n*_, the ground-truth tree *T*_GT_ *S*_GT_ might not be the one with maximum parsimony or highest likelihood.

Our main contribution is an approach to resolve (3), which is applicable and powerful for a wide choice of distance functions between trees and of cutoff values *ϵ*. Specifically, we propose a homoplasy-based test statistic that quantifies the *consistency* of the candidate tree with respect to the observed sequences *S*_*n*_ and the non-modifiability of the CRISPR-Cas9 model. As we show, our approach can distinguish accurate from inaccurate trees, under all four distance functions *d*_RF_, *d*_tri_, *d*_P_ and *d*_L_. In Section 4 we present simulation results for *d*_RF_ and *d*_P_; simulations for *d*_tri_ and *d*_L_ appear in the appendix. In addition, in Section 5 we provide theoretical support for our approach.

*Remark* 1. There is a fundamental difference between the problem defined in Eq. (3) and classical statistical hypothesis testing. The latter is often formulated as a decision problem, whether observed data was generated from an assumed statistical model (the null) or an alternative one. In this work, in contrast, we assume that the CRISPR-Cas9 statistical model is correct. Instead, we test the consistency of the “data” *TS* with the CRISPR-Cas9 model. Here, the “data” includes both the observed sequences and the reconstructed tree and its inner sequences.

## 3 A Pairwise Homoplasy Approach

To describe our approach for the hypothesis testing in Eq. (3), we first recall the definition of two classical concepts in tree phylogeny: latest (i.e. lowest or most recent) common ancestor (LCA) and homoplasy.

### Definition 2

(LCA). The latest common ancestor of a pair of leaves *u, v* with respect to a tree topology *T* is the most recent node (with latest birth time *τ*) whose descendants include both *u* and *v*, denoted LCA(*u, v*).

### Definition 3

(homoplasy and PHS). For a pair of leaves *u* and *v*, there is a homoplasy in their *i*-th character with respect to a full tree *T S* if both leaves have the same mutation, 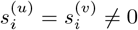, but their latest common ancestor was unmutated, 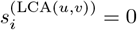. We denote the indicator of such a homoplasy in the *i*-th character of *u* and *v* by

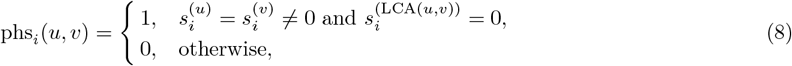

and by phs(*u, v*) the *pairwise homoplasy score* for the pair of leaves *u, v* with respect to a full tree *TS*, defined as their mean PHS over the *k* characters,

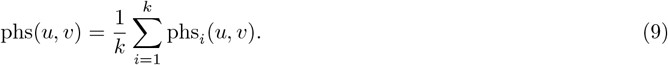

For example, consider a full tree *T S* with *s*^(*u*)^ = 10221, *s*^(*v*)^ = 13321, and *s*^(LCA(u,v))^ = 00020. Then phs_1_(*u, v*) = phs_5_(*u, v*) = 1, while phs_2_(*u, v*) = phs_3_(*u, v*) = phs_4_(*u, v*) = 0, and thus phs(*u, v*) = 2*/*5.

Note that to compute the PHS, the unobserved sequence *s*^(*w*)^ at the inner node *w* = LCA(*u, v*) needs to be known. In cases where a reconstruction algorithm provides us only with a tree topology *T* but not with its estimated inner sequences *S*, we first impute the ancestral states using a simple procedure of post-order traversal (Appendix A.2). Given two sister nodes, the state of their parent can in most cases be inferred without ambiguity due to non-modifiability. The one exception is when the two sister nodes are mutated and have the same state. In this case, we make the parsimonious choice and set the parent cell to the same state as its child nodes.

### 3.1 PHS probability distribution

Let *T*_GT_ be a fixed ground-truth topology, and suppose its sequences *S*_GT_ are generated according to the non-modifiable mutation model of Section 2.1. As the sequences are random, so are the PHS values of all pairs of tree leaves. Our test statistic is based on the tail probabilities of these values. To construct our test statistic, we shall make use of the following lemma, which describes the PHS probability distribution. Its proof is in Appendix E.

#### Algorithm 1

Computation of the PHS test statistic

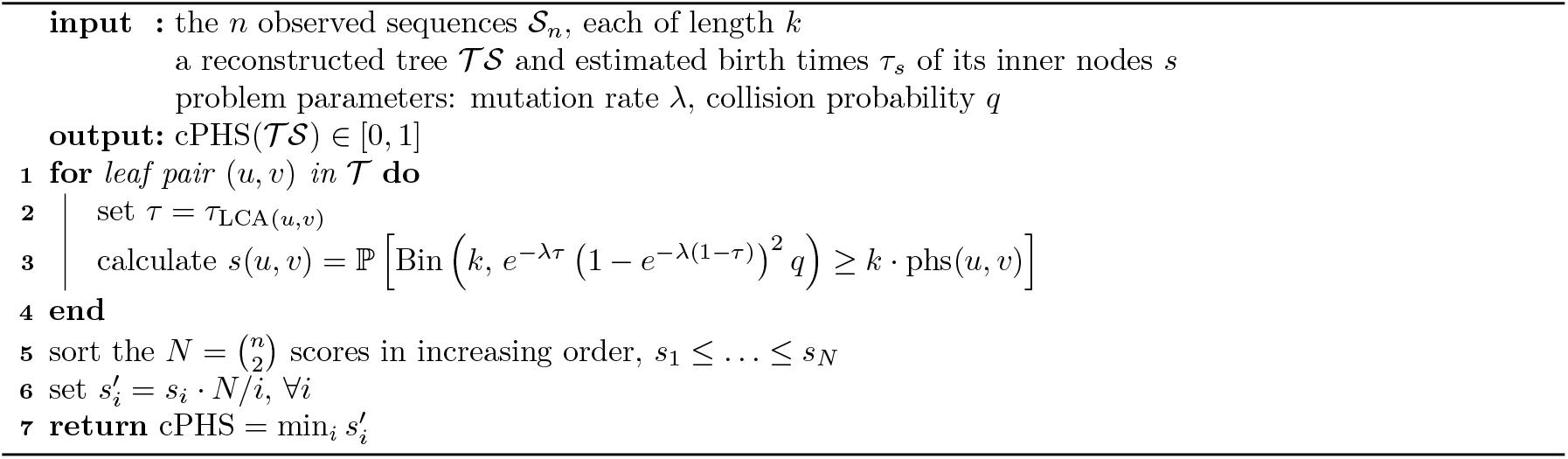

#### Lemma 1.

*Let u and v be a pair of leaves in T*_*GT*_. *Denote their LCA by w* = *LCA*(*u, v*) *and its birth time by τ*_*w*_ ∈ [0, 1). *Let* 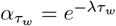 *and* 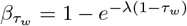, *where λ is the mutation rate. Then, for sequences of length k generated as in Section 2.1 with collision probability q (see Eq. (2))*,

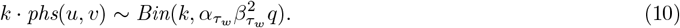

Lemma 1 reveals two appealing properties of the PHS distribution. First, in terms of the model parameters, it depends on the mutation rate *λ*, sequence length *k* and the collision probability *q*, but not on the individual mutation probabilities *q*_*j*_. Second, for any pair of leaves *u, v*, the PHS distribution does not depend on the full structure of the tree, but only on a single sufficient statistic: the birth time of their LCA. This simplifies the computation of our proposed test statistic (described in the next section), and facilitates its theoretical analysis (see Section 5).

### 3.2 A PHS-based test statistic

We are now ready to present our proposed PHS test statistic for the hypothesis problem in (3). For simplicity, in this section we assume the model parameters *λ* and *q*, as well as the birth times of all inner nodes of *T*, are known. As described in Appendix A.1, in our simulations we estimate *λ* and the birth times from the data. The collision probability *q* is assumed to be known also in simulations since, as detailed in Appendix A.1, *q* can be accurately estimated in various experimental settings. Moreover, as illustrated in Figure S5 (Appendix K), our test statistic is not sensitive to the exact value of *q*, and taking *q* = 1*/m*, which corresponds to a uniform mutation distribution *q*_*j*_ = 1*/m*, works well.

The calculation of our test statistic is outlined in Alg. 1. Given a candidate tree *T S*, it consists of two steps.

#### Step 1 (calculating tail probabilities)

Based on Lemma 1, for each pair of leaves (*u, v*) with latest common ancestor *w* = LCA(*u, v*) in the reconstructed tree *T*, we compute the tail probability of its PHS value as if this tree were the ground-truth one,

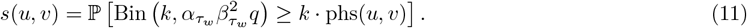

Each score *s*(*u, v*) can be viewed as a p-value (i.e., right-tail probability under ℋ_0_) for the consistency of the observed sequences at the nodes *u* and *v* with the reconstructed tree. Since there are *n* leaves, Eq. (11) provides us with 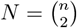 scores. The next step is to fuse these multiple p-values into a single score.

#### Step 2 (fusing the multiple scores)

Motivated by multiple hypothesis testing procedures, we sort the *N* scores in increasing order, *s*_1_ ≤ *s*_2_ ≤ … ≤ *s*_*N*_, and focus on the first few smallest ones. Similar to the Benjamini-Hochberg (BH) procedure (Benjamini and Hochberg, 1995) (see Remark 3 below for more details), for each *i ≤ N* we compute an adjusted score

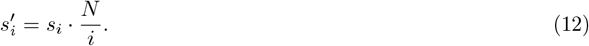

Finally, our test statistic is the minimal adjusted score,

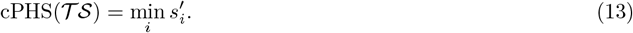

Note that cPHS(*T S*) ≤ 1, since 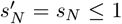. Given a reconstructed tree *T S*, we reject ℋ_0_ if cPHS(*T S*) is below some threshold *t* < 1, and accept it otherwise.

*Remark* 2. Let us provide some motivation and context for the above procedure. Often, some parts of the re-constructed tree are close to the ground truth. By construction, the corresponding scores *s*(*u, v*) in those parts are distributed approximately uniformly over [0, 1]. In contrast, other parts of the reconstructed tree may be less accurate. In those cases, some of the scores *s*(*u, v*) are expected to be closer to zero. Typically, the number of *s*(*u, v*) values that are close to zero is relatively small w.r.t. *N* ≫ 1. Hence, our setting is similar to high-dimensional multiple hypothesis testing problems, whereby only a few out of many hypotheses deviate from the null.

*Remark* 3 (Relation to the BH procedure). As described in Benjamini, Heller, and Yekutieli (2009, Section 2(b)), the BH procedure can be presented in terms of adjusted p-values, defined as

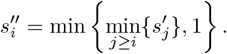

Under this formulation, all hypotheses *i* for which 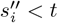 are rejected. In our case, we do not aim to reject or accept individual hypotheses, corresponding to individual p-values 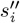. Instead, we adopt a more stringent criterion: if even a single adjusted score 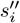 falls below the threshold, the entire tree is rejected. Accordingly, our test statistic coincides with the minimum BH-adjusted p-value:

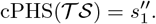

As illustrated in Section 4 and in Appendix K below (see e.g. Figure 1(d)), cPHS is highly powerful in distinguishing between accurate and inaccurate trees. Furthermore, as cPHS separates accurate and inaccurate reconstructed trees by a large margin (see Figure 1), it does not require a delicate tuning of the threshold *t*. In practice, as illustrated in Section 4, a threshold value that depends only on the cutoff value *ϵ* of Eq. (3) works well for a wide range of model parameters (such as the sequence length *k*, the number of mutation states *m*, etc.).

**Figure 1:**
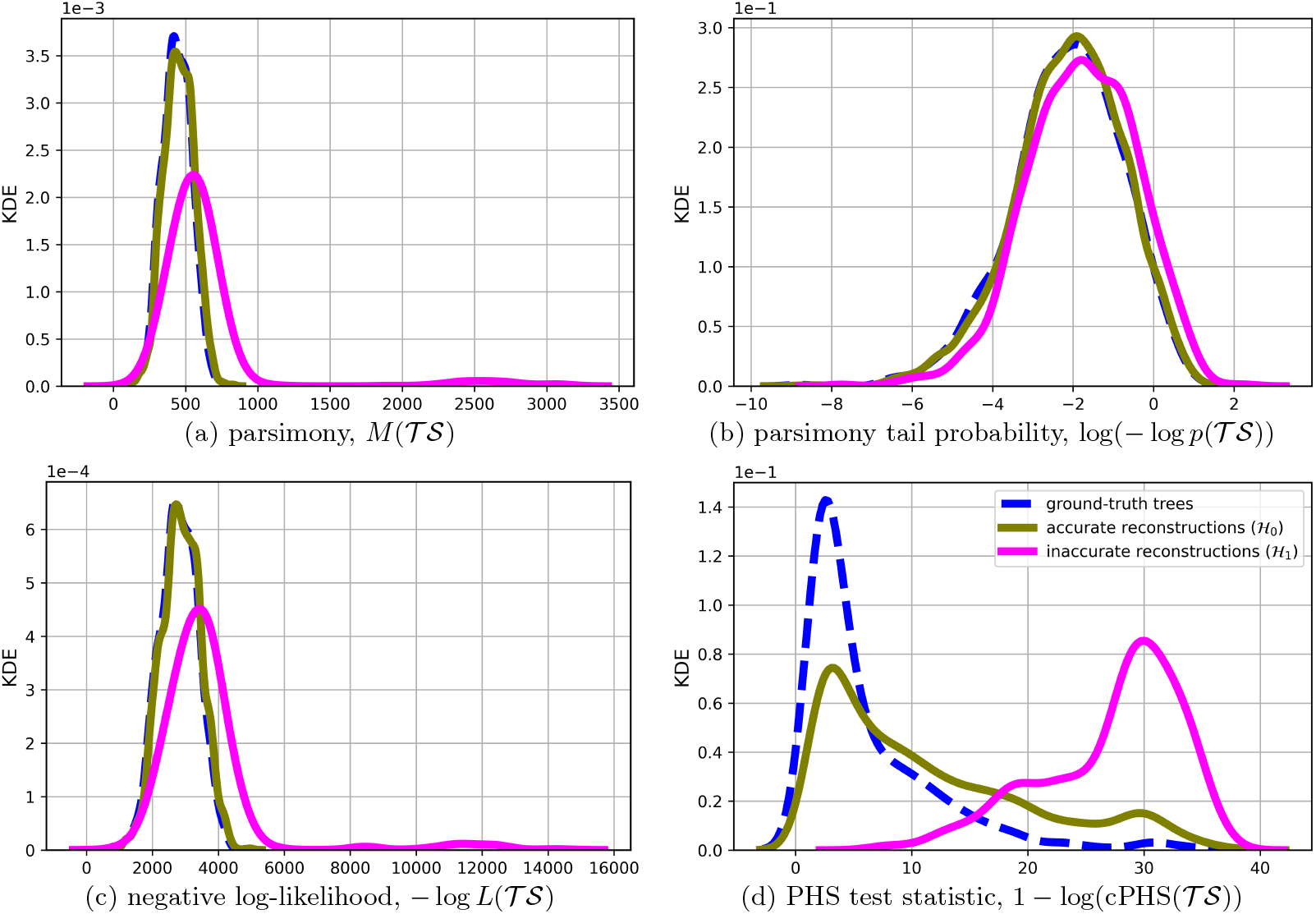
Kernel density estimates for four accuracy measures: parsimony (upper left); parsimony tail-probability (upper right); likelihood (bottom left); and our proposed cPHS (bottom right). The parsimony tail probability and cPHS test statistics are log-transformed for clarity of view. The hypotheses *H*_0_ and *H*_1_ are defined by (3) with *d* = *d*_RF_ (4) and *ϵ* = 1*/*4. The simulations were performed using Cassiopeia (Jones et al., 2020) as follows: binary tres with 10^6^ terminal cells were generated, from which *n* = 10^3^ observed cells were subsampled (subsampling ratio of 10^−3^). The sequence length is *k* = 30, each site has *m* = 32 possible mutations with probabilities *q*_*j*_ drawn from an exponential distribution, and *λ* = − log 2, so that there is *ρ* = 50% probability to observe a mutation at a leaf. These parameter values are similar to those of experimental settings, see Jones et al. (2020).

*Remark* 4 (Non-uniform mutation rate). Lemma 1, and thus Eq. (11), assume that the mutation rate *λ* is the same for all *k* characters. However, as long as characters are assumed to evolve independently, our PHS approach can be easily extended to the case of non-uniform mutation rates; see Claim 1 in Appendix E.

#### Complexity and runtime

Given a reconstructed tree *T S*, computing its cPHS value can be done in a polynomial number of operations. Specifically, as the algorithm passes over all leaf pairs for each character, its time complexity is *O*(*kn*^2^). With our Python implementation, calculating cPHS for a tree with *n* = 1000 leaves and sequences of length *k* = 50 takes approximately two minutes on a standard PC.

#### Missing data

In practical settings, some entries of *S*_*n*_ may be missing, due to either limited sensitivity (the socalled stochastic drop out) or heritable mechanisms (e.g. concomitant resection of two adjacent target sites (Jones et al., 2020)). As described in Appendix B, our PHS approach can be easily extended to handle missing data, and empirically, it continues to perform well also in such cases.

## 4 Simulations

We demonstrate through a comprehensive set of simulations the capability of the cPHS test statistic (13) to resolve the hypothesis testing problem outlined in (3). We compare cPHS with three test statistics: (i) the parsimony of the reconstructed tree, *M* (*T S*); (ii) t he negative log-likelihood of the reconstructed tree, *L*(*T S*); and (iii) the parsimony tail-probability, *p*(*T S*) =_*M*≥*M*(*T S*)_ ℙ [*M* | *T*]. The last test statistic is based on the parsimony probability distribution *conditional* on the specific reconstructed tree topology *T*, denoted as ℙ [*M* | *T*]. As described in Appendix D, this distribution can be computed analytically.

For each simulation in this section, we used Cassiopeia (Jones et al., 2020) to generate 1000 ground-truth trees and their inner sequences, *T*_GT_ *S*_GT_. Similar to other studies (Jones et al., 2020; Feng et al., 2021; Seidel and Stadler, 2022; Sashittal et al., 2023), each ground-truth tree topology was randomly generated according to a birth-death process. The rates of birth and death are not homogeneous throughout the tree but instead depend on a heritable, sub-clonal fitness level; see Appendix J. Given a tree topology, its inner sequences were generated according to the non-modifiable CRISPR-Cas9 mutation model described in Section 2.1. The considered settings are similar to those in Sashittal et al. (2023): sequence length of *k* = 30, *m* = 32 possible mutations, and *n* = 200 cells. In this section, the matrix *S*_*n*_ is fully observed; simulations with missing data appear in Appendix B. The observed data *S*_*n*_ from each ground-truth tree was given as input to seven reconstruction algorithms: Neighbor-Joining (Saitou and Nei, 1987), Cassiopeia-Greedy (Jones et al., 2020), Shared-Mutation-Joining (Wang et al., 2023), MaxCut and MaxCut-Greedy (Snir and Rao, 2006; Jones et al., 2020), Spectral and Spectral-Greedy solvers (Jones et al., 2020). The seven reconstructed trees were then classified as either *accurate* or *inaccurate* according to (3), using normalized RF (*d* = *d*_RF_) and parsimony (*d* = *d*_P_) as distance functions. Results for the other two distance functions, triplets (*d* = *d*_tri_) and likelihood (*d* = *d*_L_), are provided in Appendix K. We remark that in general, the reconstructed binary tree, as well as the ground-truth one, may contain edges that are not supported by any mutations. Similar to Sashittal et al. (2023) and Dai and Molloy (2026), we first contract mutationless branches, and compute the distance functions on the contracted trees. In particular, if two original binary trees disagreed only on mutationless branches, after this procedure, they are viewed as topologically equivalent, and their RF distance is zero.

### 4.1 Separability Capabilities of cPHS

To motivate our approach, we first demonstrate that given a reconstructed tree *T S* corresponding to an unknown ground truth *T*_GT_*S*_GT_, existing measures are unable to accurately distinguish between the null and the alternative hypotheses in (3). In this simulation, we considered a tree as inaccurate if *d*_RF_(*T S, T*_GT_*S*_GT_) > 1*/*4. This corresponds to the hypothesis testing problem in Eq. (3) with distance function *d* = *d*_RF_ and *ϵ* = 1*/*4; results for the other three distances *d* appear in Appendix K.

As shown in panels (a-c) of Fig. 1, for each of the three baseline test statistics there is substantial overlap between their values on accurate and inaccurate reconstructed trees. A similar overlap exists even between ground-truth and inaccurate reconstructed trees. This substantial overlap implies that *regardless of the chosen threshold t*, these measures cannot accurately separate ground-truth trees, or accurate trees, from inaccurate ones. In Figures S7 and S8 of Appendix K, we show that even when accuracy is defined by the *parsimony* -based distance *d*_P_ (rather than the normalized RF), both the parsimony score and its tail-probability measure are ineffective in distinguishing between accurate and inaccurate reconstructed trees.

In contrast, Fig. 1(d) shows that our proposed PHS-based test statistic, described in Section 3, achieves a high separation between candidate trees with low (good) and high (bad) RF distances. Furthermore, it achieves a *nearly perfect* separation between ground-truth trees and candidate trees with high RF distance. In the next subsection and Appendix K we show that a similar conclusion holds for the other distances *d*_RF_, *d*_tri_, and *d*_L_. Specifically, for each of these other distance functions, there is significant overlap in the distribution based on the parsimony or likelihood measures. In contrast, our PHS-based test statistic achieves a high separability. Notably, as demonstrated in Appendix K, cPHS is a better accuracy measure than parsimony even if the distance measure is parsimony itself, and similarly for likelihood. That is, if our goal is to tell whether the parsimony/likelihood of the candidate tree is close to the ground-truth parsimony/likelihood, it is more useful to look at its cPHS than at its parsimony/likelihood.

In Section 5 we provide theoretical support for the empirical advantage of our homoplasy-based approach over parsimony. Specifically, we prove that to detect certain incorrect topologies, parsimony requires exponentially longer sequences (in tree depth) compared to our PHS approach.

### 4.2 Classification Accuracy

The previous section demonstrated the separability capabilities of cPHS in comparison to other test statistics. In practice, however, we aim to classify reconstructed trees as either accurate or inaccurate by applying a threshold to these test statistics. In this section, we evaluate the classification performance of each baseline test statistic by assessing its ability to classify correctly if a reconstructed tree is accurate or not. Given a distance function and a cutoff value *ϵ*, for any tree denote by *y* ∈ {accurate, inaccurate} its true label according to (3). Similarly, denote by *ŷ*_*t*_ the label estimated by some test statistic with a threshold *t*. For example, for cPHS,

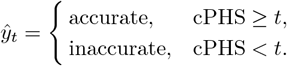

We first examine the performance of the four test statistics across all possible thresholds, then focus on specific choices of *t*. In the first simulation, we compute the true positive rate (TPR) and false positive rate (FPR) of each test statistic as a function of the threshold *t*. These rates are defined as follows:

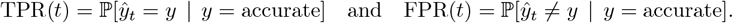

Plotting the TPR and FPR for all threshold values *t* gives ROC curves, presented in Fig. 2, with 95% confidence intervals computed via bootstrap. Panel (a) displays the results for *d* = *d*_RF_ with *ϵ* = 1*/*7, and panel (b) displays the results for *d* = *d*_P_ with *ϵ* = 0.01. It is apparent that, *regardless* of the selected threshold *t*, the performance of cPHS is superior to that of the other test statistics. Notably, the ROC curve of cPHS exhibits a steep ascent on the left side, indicating that cPHS achieves a satisfactory TPR even at low FPR levels.

**Figure 2:**
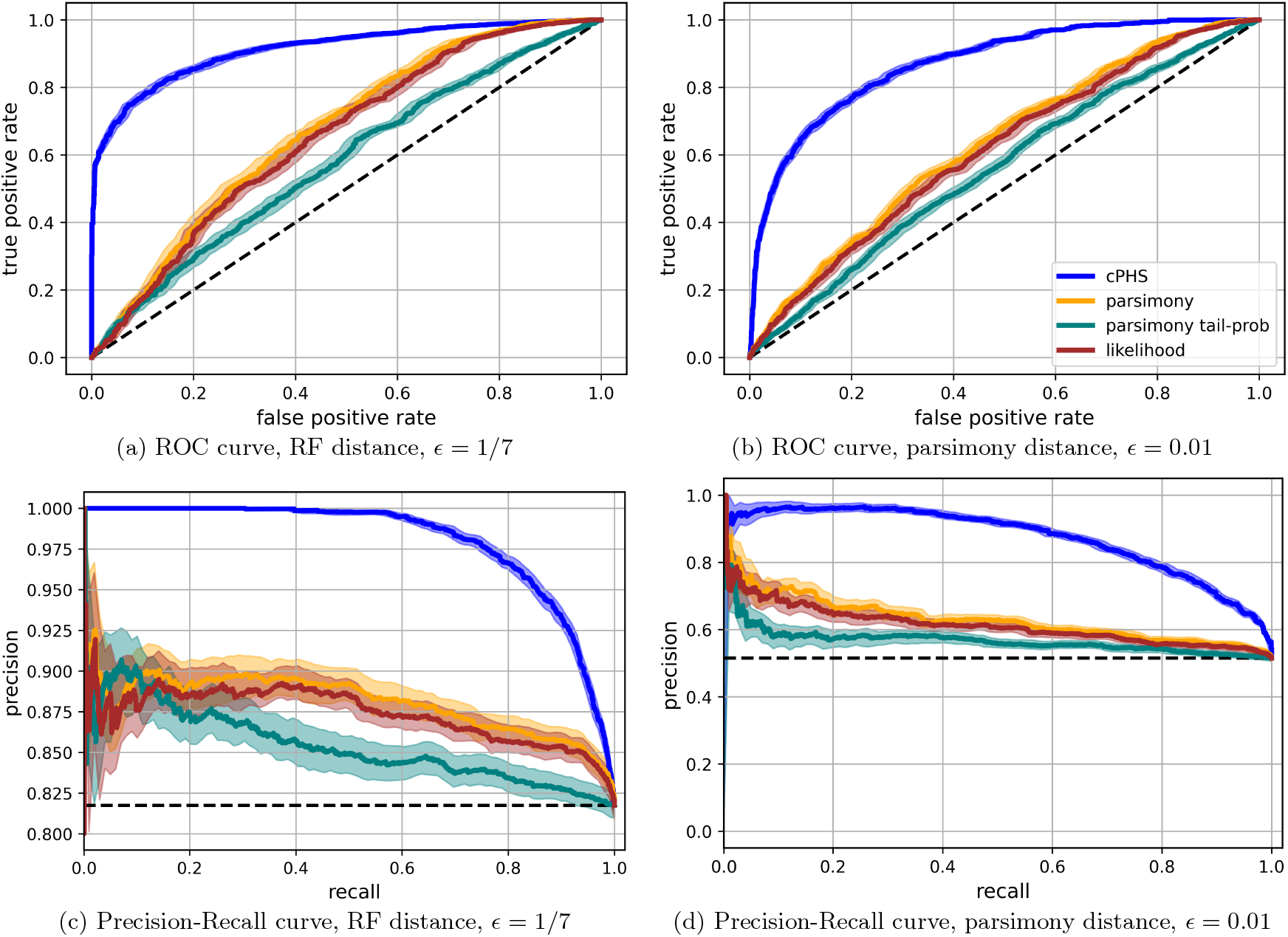
ROC (top) and Precision-Recall (bottom) curves of several test statistics for the hypothesis testing in (3). Each point on the curve corresponds to a different threshold *t*. The difference between the left and right panels is the distance measure in use: normalized RF with *ϵ* = 1*/*7 (left) and parsimony distance with *ϵ* = 0.01 (right). Dashed lines represent the expected performance of random guessing. The rates are calculated over 1000 realizations.

Panels (c) and (d) of Fig. 2 present Precision-Recall curves. These metrics are defined as follows:

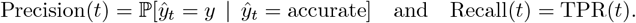

A clear distinction in performance between cPHS and the other test statistics is evident under these metrics as well. Specifically, when the hypotheses in (3) involve the RF distance (*d* = *d*_RF_), as seen in panel (c), cPHS achieves a high Precision of 0.93 at an equally high Recall.

The simulations above were performed with sequence length *k* = 30 and number of mutated states *m* = 32. Next, we examined the performance as these parameters are varied. Figure 3 shows the AUC (area under the ROC curve) as a function of the sequence length *k* (left panel) and of the number of mutated states *m* (right panel). Additional results for varying values of *m, n* and *ρ* (bijectively related to *λ*, see (1)) can be found in Appendix K, as well as results for the AUPRC (area under the Precision-Recall curve). As shown, under a wide range of settings, cPHS consistently outperforms the other test statistics by a significant margin.

**Figure 3:**
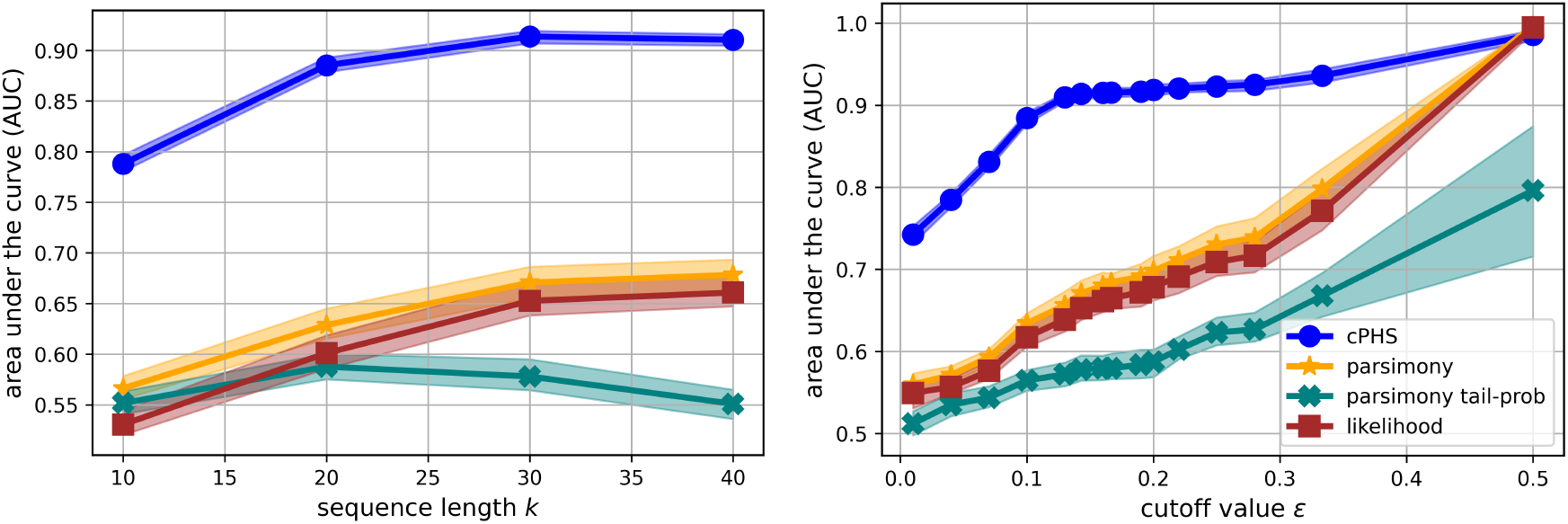
AUC values for the ROC curves of several test statistics for the hypothesis testing in (3) with *d* = *d*_RF_. Left: *ϵ* = 1*/*7 is fixed and the sequence length *k* is varying. Right: *k* = 30 is fixed and *ϵ* is varying. Each point is based on 1000 realizations of the simulation.

We now turn to results obtained for specific choices of the threshold *t*. Our goal is to evaluate the robustness of cPHS when using a fixed threshold, rather than optimizing *t* for each parameter setting. To this end, we focus on the balanced accuracy metric, which is commonly used to evaluate classification performance. Because our dataset exhibits approximately equal class support, we prefer this metric over the F1-score to ensure a symmetric evaluation that fully incorporates true negatives. Balanced Accuracy is defined as the average of sensitivity and specificity:

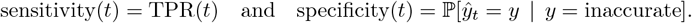

For each parameter configuration, we compute the balanced accuracy across several thresholds and record the maximum value. We then compare this optimal performance to that of cPHS using a fixed threshold *t* chosen based only on *ϵ*, independent of model parameters like *k* or *m*. Figure 4 summarizes this comparison. Specifically, the left panel shows results for different values of *k* with a cutoff value fixed at *ϵ* = 1*/*7, and the right panel for different values of *ϵ* with *k* fixed at 30. In the left panel, the cPHS threshold is fixed at *t* = 10^−3^. Two conclusions can be drawn from these results: First, the performance of cPHS is similar for a fixed threshold (based solely on *ϵ*) and an optimally tuned one. Second, whether using fixed or optimal thresholds, cPHS consistently outperforms all other test statistics, evaluated with their own optimal thresholds. Additional results in Appendix K confirm that cPHS maintains strong performance with a fixed threshold across a wide range of values of *m, n* and *ρ*, as well as for a different value of *ϵ*.

**Figure 4:**
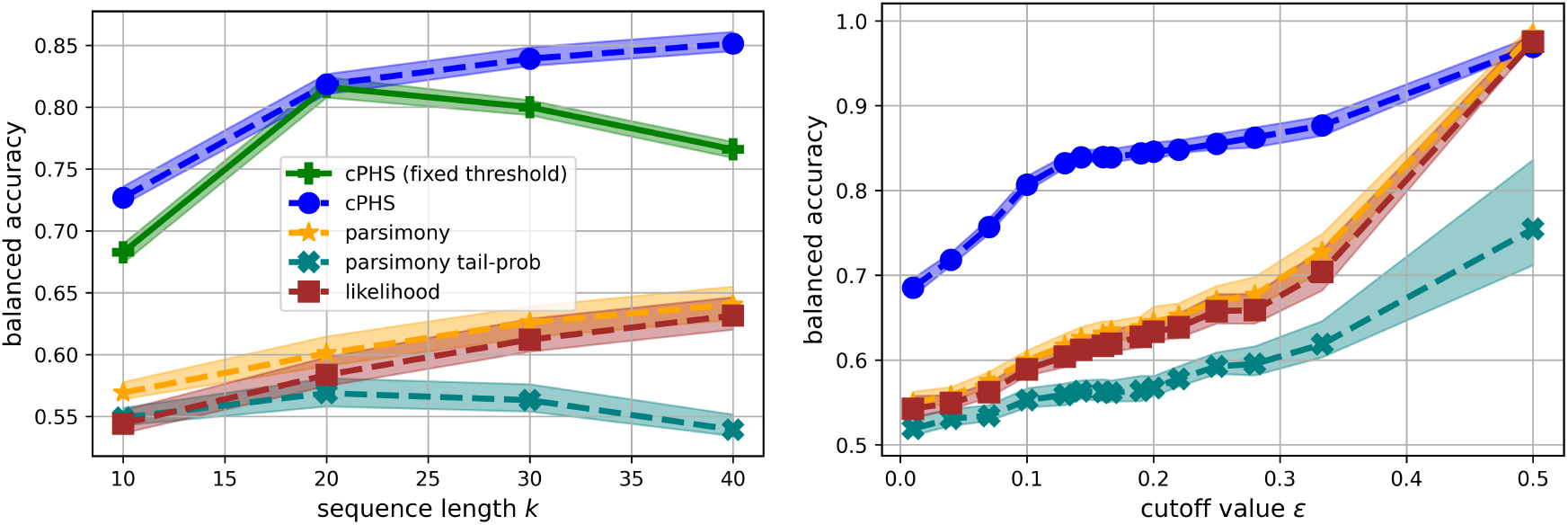
Balanced accuracy of several test statistics for the hypothesis testing in (3), with the same setting as in Figure 3. Notably, cPHS with a fixed threshold achieves a significantly higher balanced accuracy than the other approaches, even when those are applied with optimally tuned thresholds.

### 4.3 Identification of the Ground-truth tree

Finally, we illustrate the statistical power of cPHS over parsimony and likelihood by a different type of analysis. Specifically, we consider the following question: Suppose we are given a set of candidate trees, which includes trees reconstructed by various algorithms as well as the ground-truth tree. Is it possible to detect which tree is the ground-truth one? In other words, what is the probability that the test statistic of the ground-truth tree is better than its values for all reconstructed ones? Figure 5 illustrates results for various test statistics as a function of the sequence length *k*. The left panel illustrates the advantage of cPHS over parsimony and likelihood in the setting considered in this section thus far, namely with *n* = 200 cells and *m* = 32 possible mutations. The right panel of Figure 5 shows results for larger trees, with *n* = 10^3^ observed cells subsampled from total of 10^6^ in the original tree and *m* = 50. Notably, in this setting the ground-truth tree is very rarely the most parsimonious or the maximum likelihood tree. In contrast, in approximately 40% of the cases, cPHS(*T*_GT_*S*_GT_) > cPHS(*T S*) for all reconstructed trees *T S*. These results are in agreement with those described in previous sections.

**Figure 5:**
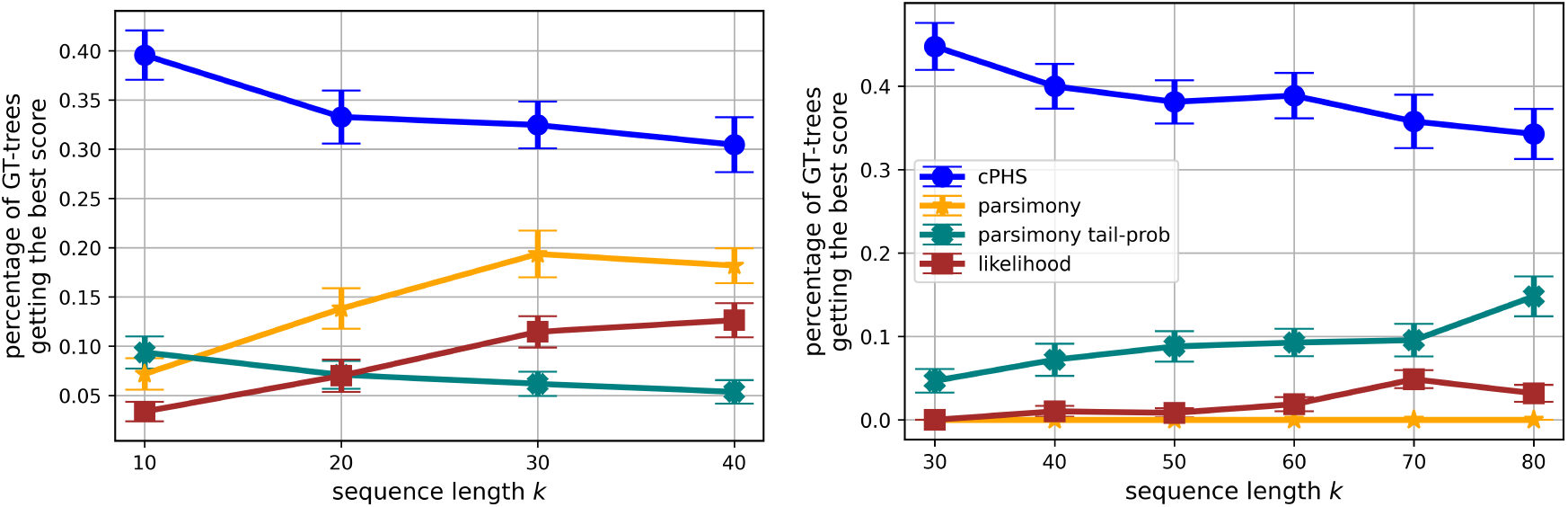
Ability to detect the ground-truth tree as a function of the sequence length *k*. Left: same setting as in Figure 3; in particular, *n* = 200, *k* = 30 and *m* = 32. Right: larger trees, with *n* = 10^3^ cells, *k* = 50 and *m* = 50. Each point is calculated over 1000 realizations.

## 5 Theoretical Support for PHS

In this section, we present theoretical support for our PHS-based approach. In Section 3.2 we proposed to assess the accuracy of a full tree by the cPHS test statistic (13). This quantity, however, is difficult to analyze theoretically, as it depends in a non-trivial way on the tail probabilities of the PHS values at all leaf pairs in the tree. Instead, we analyze a simpler PHS-based test statistic, tPHS(*TS*), defined below. Our main result is that even this simpler measure is significantly more powerful than parsimony in detecting inaccurate tree topologies.

The PHS-based test statistic we consider in this section is defined as follows.

### Definition 4

(Total PHS). The total PHS of a full tree *T S* is the sum of the PHS values over all its 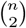 pairs of terminal leaves,

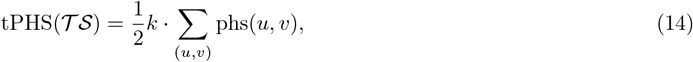

where phs(*u, v*) is defined in Eq. (9).

To simplify the analysis, we assume that the ground-truth tree has a homogeneous (binary) tree topology.

### Definition 5

(Homogeneous tree topology). A tree *T* is called homogeneous if all its edges have the same length (elapsed time). The depth *d* of the tree is the number of edges from the root to any of its leaves.

We consider the following scenario. Given the sequences at the terminal leaves, we assume that an algorithm reconstructed a tree topology *T* which is nearly identical to the ground-truth *T*_GT_, and differs from it by a single *swap* of two leaves *u, v* from different sides of the original tree. Our key result is that tPHS is *qualitatively* more powerful than parsimony in detecting such incorrect topologies. We show that the difference in tPHS between the incorrect tree and the ground-truth one, normalized by the tPHS standard deviation, is much larger than the parsimony difference, normalized by its standard deviation. In simple words, the tPHS measure can provably distinguish between such correct and incorrect reconstructed trees, whereas parsimony cannot.

Next, we formally define a leaf swap. An illustration appears in Fig. S2 of Appendix F.

### Definition 6

(Leaf swap). Let *u, v* be a pair of leaves in a full tree *T*_GT_*S*_GT_. Denote by (*T S*)_*u* ↔*v*_ the tree obtained from *T*_GT_ *S*_GT_ by swapping the leaves *u* and *v*. This swap operation is followed by a correction of the sequences in the ancestors of *u* and *v* to satisfy the non-modifiability constraint: at each character *i* ∈ [*k*], if a mutated ancestor *w* of *u* or *v* has an unmutated descendant, or if two of its descendants are mutated to different states, then the ancestor is set to be unmutated, 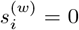. Further, denote the difference in tPHS and parsimony values following the swap by

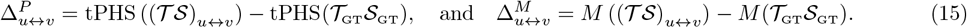

The following theorem shows that the difference 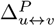 is much larger than 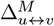, both properly normalized.

The proof appears in Appendix H.

### Theorem 1.

*Let T*_*GT*_ *S*_*GT*_ *be a homogeneous full tree of depth d whose sequences, of length k, were generated according to the model in Section 2.1, with mutation rate λ, collision probability q, and mutation probability at a leaf ρ given in* (1). *Let u and v be a pair of leaves whose LCA*(*u, v*) *is the tree root. Let* 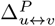 *and* 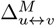 *be the respective change in tPHS and in parsimony, as defined in* (15), *following the swap u* ↔ *v*. *Then*, *for a sufficiently large d, their normalized expectations* 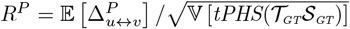 *and* 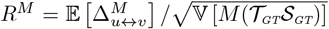 *satisfy the following lower bound and upper bound, respectively*,

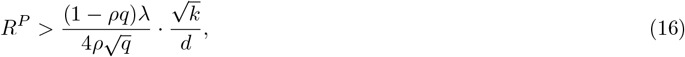

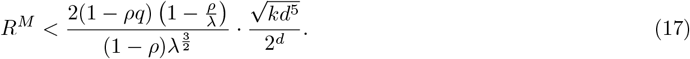

The theorem holds provided that *d* is sufficiently large, with the required bound depending on *λ*. For instance, when *λ* ≤ 3, it suffices to have *d* ≥ 10.

Let us discuss the consequences of this theorem. Consider a large tree with *n* ≫ 1 leaves, and depth *d* = log_2_ *n*. Viewing the model parameters *λ, q* as fixed, according to Eq. (16), for the change in tPHS to be significant it suffices that 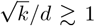. That is, the required sequence length scales only *polylogarithmically* with the number of leaves, *k* ≳ (log_2_ *n*)^2^. We note that, as proven in Erdős et al. (1999) and Wang et al. (2023), a similar relation between *k* and *n* suffices for accurate tree reconstruction under reversible and non-modifiable mutation models, respectively. In contrast, for the change in parsimony to be significant, a necessary condition is that the right-hand side of (17) is large. This translates into a significantly longer sequence of length *k* ≳ *n*^2^*/*(log_2_ *n*)^5^, which grows *polynomially* with *n*. In simple words, compared to tPHS, parsimony requires exponentially longer (in tree depth) sequences to detect an incorrect topology of the form of a leaf swap.

*Remark* 5 (leaf swap on the same side of tree). Theorem 1 addresses the swap of leaves whose LCA is the tree root. The theorem can be easily generalized to the swap of leaves *u, v* whose LCA(*u, v*) is an inner node. This would result in two changes to the RHS of Equations (16) and (17). First, instead of *d* we would have *d* − *l*, where *l* is the depth of LCA(*u, v*). Second, instead of *k* we would have *k* · *e*^−*λl/d*^, the expected number of unmutated characters at the node LCA(*u, v*). As long as *l* is much smaller than *d*, namely LCA(*u, v*) is close enough to the root, the consequences of the theorem continue to hold. Specifically, tPHS would be able to detect such a leaf swap with a sequence length polylogarithmic in *n*, whereas parsimony would require a sequence length polynomial in *n*.

*Remark* 6 (several leaf swaps). Theorem 1 considers a single leaf swap. The theorem can be readily extended to the case of several leaf swaps, provided that the different LCAs of the leaf pairs are located at distinct branches of the tree. In particular, the LCAs must be inner nodes; see Remark 5. In this case, the effect of the leaf swaps on both the tPHS and parsimony is additive.

## 6 cPHS evaluation on a dataset of lineage tracing in lung adenocarcinoma

To evaluate the cPHS score with real-world data, we applied it to a dataset of lineage tracing in a mouse model of lung adenocarcinoma from Yang et al. (2022). This dataset features an inducible lineage recording system consisting of Cas9, synthetic DNA elements that can be targeted by Cas9 (acting as lineage recording sites), and guide RNAs that help direct Cas9 to those sites. The lineage recording system was integrated into the genome of mice that already harbored inducible oncogenic mutations of the genes Kras and Trp53. By direct administration of Cre-carrying virus to the lungs of the engineered animals (referred to as KPTracers), the authors have induced two processes that operate on the lung epithelium in parallel: malignant transformation (switching on oncogenic mutations in Kras and Trp53) and lineage recording (switching on Cas9). The resulting tumors, including both primary and metastatic lesions, were obtained between five to six months after induction and analyzed using single cell RNA-sequencing. This analysis provided both lineage traces (i.e., the sequence of each target site - either mutated or untouched) and transcriptome information (i.e., estimation of expression for thousands of genes) for each tumor cell.

Our dataset consisted of 63 samples (tumors), spanning a wide range of parameters, including the number of cells per tumor (106 ≤ *n* ≤ 4798, average = 689), collision probabilities (0.06 ≤ *q* ≤ 0.77, average = 0.28), mutation rates (1.2 ≤ *λ* ≤ 7.3, average = 3.1), and number of successfully captured recording sites (6 ≤ *k* ≤ 67, average = 25). See Supplementary Table (`KPTracer_summary.xlsx`) for the complete description of these samples and their analysis. To evaluate cPHS we first generated a set of negative controls for each tumor, consisting of five random binary trees built over the same set of leaves (i.e., tumor cells and their lineage traces). To compute cPHS scores of these trees we followed the rationale presented in Section 4 and used dedicated simulation analysis to select threshold values *t* based on the observed parameters (see Appendix M). We find that the ability to reject random trees tracks closely with the quality of the respective sample (Figures S17, S18). Specifically, randomized trees built for the 42 samples that had a sufficient number of recording sites (*k* > 15) and did not exhibit high collision probabilities (*q* < 0.35) were all rejected (0 of 210 random trees accepted as accurate by cPHS). These correspond to good quality samples with sufficient rates of integration of lineage recording sites and to a sufficient capture rate of RNA molecules (resulting in sufficiently large *k*). They also correspond to cases with well-calibrated affinity of the guide RNAs, avoiding early saturation in which many cells in the eventual tumor have identical lineage traces (thus high *q*). These results are further supported by a more extensive simulation analysis, finding that also in fully simulated data where the ground truth is known, in the regime of insufficient-quality (*k* ≤ 15 or *q* ≥ 0.35) no value of *t* reliably separates accurate from inaccurate reconstructions for any reasonable choice of *ϵ* (Tables S1, S2, S3, S4).

We next reconstructed lineage trees for each tumor sample using six algorithms that are implemented in the Cassiopeia package (Jones et al., 2020), including Neighbor Joining, Shared Mutation Joining (SMJ), Max-Cut, a compatibility-based heuristic (Cassiopeia greedy), and two algorithms based on spectral decomposition. Of the 42 tumors with sufficient quality, cPHS confirmed at least one of the six reconstructions as valid in 28 of the cases. Importantly, all the 14 samples for which the trees were rejected fall within a regime of low *k* to *n* ratio (*k/n* < 0.05). All other samples had a higher ratio and had at least one inferred tree (out of the six inferred per sample) that was accepted by cPHS. This result mirrors the theoretically-established relationship between *n* and the minimal value of *k* that is required to achieve accurate reconstruction (Wang et al., 2023). Among the algorithms used for inference, SMJ, Max-Cut, and compatibility-based greedy received the highest acceptance rate by cPHS (50% − 55% of trees; see Figure 6). The two spectral methods had the lowest acceptance rate, and consistently had the lowest ranks in terms of parsimony and likelihood (Figure 6).

**Figure 6:**
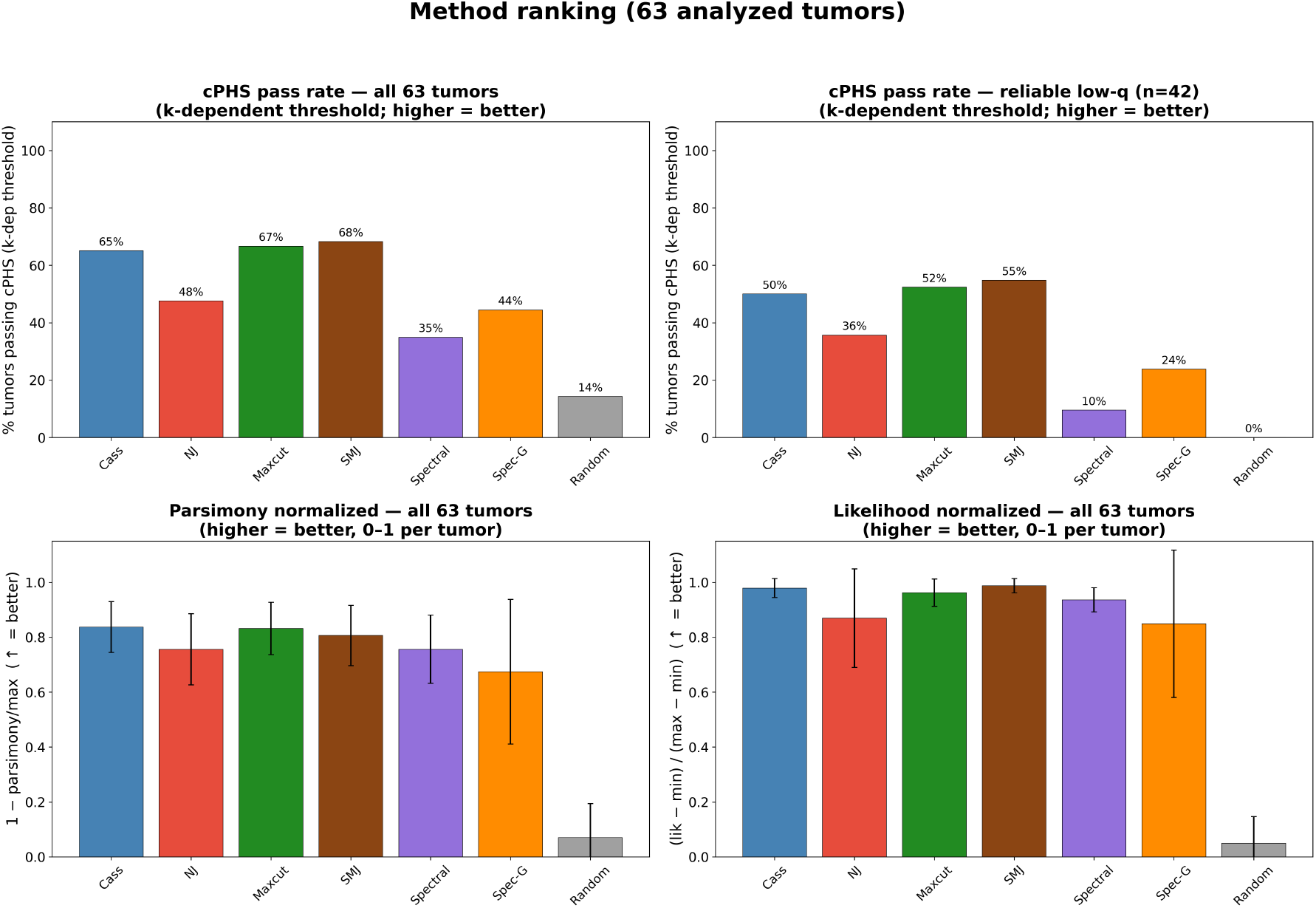
Method ranking. Top row: cPHS pass rate across all 63 tumors (left) and across the 42 reliable low-*q* tumors (right), both using the calibrated *k*-dependent thresholds (*t* = 10^−3^ for *k* ≤ 24, *t* = 10^−4^ for *k* ≥ 25). Bottom row: normalized parsimony (left) and normalized likelihood (right) across all 63 tumors, where higher values indicate better performance. Error bars show standard deviations across tumors.

We next aimed to verify that cPHS reflects topological errors rather than merely separating reconstructions from random trees. Focusing on two of the inference algorithms (SMJ and Cassiopeia greedy), we degraded the reconstructed trees by a controlled number of Nearest Neighbor Interchange (NNI) moves, which swap subtrees across internal edges and progressively scramble the topology while leaving the character data unchanged. As the fraction of NNI moves increases, the topology drifts further from the original reconstruction. The various metrics reflect this worsening: parsimony and Robinson-Foulds distance rise, while likelihood and triplets decrease (Figure S21). The cPHS acceptance rate acts accordingly with the level of perturbation, with a monotonic decline as the NNI rate increases (Figure S20; Appendix P).

Together, these results demonstrate the applicability of cPHS to real-world data and particularly the merit of enabling an accept/reject decision, beyond ranking. First, it flags parameter regimes in which the inferred trees are a-priori less likely to be accurate, corresponding to tumor samples with insufficient quality or to large samples with too few recording sites. Second, it is able not only to distinguish between random and inferred trees by its raw score, but to also reject all random trees using its classification step. Finally, it is able to sense increasingly-dominant perturbations to the inferred trees, with a reduction in the acceptance rates.

## 7 Discussion

As demonstrated via simulations in Section 4 and supported by theoretical analysis in Section 5, our method offers significantly greater discriminative power than traditional parsimony. Intuitively, this advantage may be explained as follows: parsimony aggregates a local measure — the number of mutations per edge — without accounting for the broader tree structure. In contrast, PHS integrates information across all pairs of leaves, both close and distant, thereby capturing the consistency of the tree’s topology as a whole.

In our work, we considered the minimal adjusted p-value as our test statistic (13). The key parameter of our procedure is the threshold *t* against which the cPHS statistic is compared. As illustrated in Section 6, in practical settings, a suitable threshold *t* can be estimated via simulations designed to reflect the characteristics of the experimental setting. Specifically, one may generate multiple ground-truth trees, for example via Cassiopeia (Jones et al., 2020). Given the observed sequences for each ground-truth tree, various reconstruction algorithms can be run, followed by computing the cPHS scores for their outputs. A suitable threshold can then be determined based on the cPHS scores obtained for the most accurate reconstructions. We remark that there are other possible schemes to fuse the scores in (11) into a single test statistic, e.g. the harmonic mean (Wilson, 2019). Our simulations indicate that the statistical power of the harmonic mean is comparable to that of cPHS. However, it is less practical, as in contrast to cPHS, the harmonic mean is very sensitive to the specific value of the threshold *t*.

In this manuscript, we focused on a non-modifiable model of evolution, whereby Lemma 1, and consequently our cPHS test statistic (13), are based on this assumption. Our proposed homoplasy approach may also be beneficial for assessing the accuracy of reconstructed trees under reversible models. This requires deriving an analogue of Lemma 1 for a given reversible model of evolution. We leave this extension for future work.

Another interesting direction for future research is to use a homoplasy approach not merely to assess the accuracy of trees, but in fact to develop a PHS-based tree reconstruction algorithm. As we illustrated both empirically and theoretically, PHS provides a better tree accuracy measure than parsimony. Hence, it would be interesting to explore if a PHS-based algorithm can reconstruct more accurate trees than those found by current methods that aim to find the tree with maximum parsimony for the observed data.

Finally, the core principles of the PHS framework can be leveraged to evaluate branch support. Currently, cPHS(*T S*) acts as a global test statistic by aggregating the tail probabilities of all 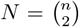 leaf pairs in the tree. However, to assess the confidence of a specific internal branch, one could restrict this evaluation to the subset of leaf pairs whose latest common ancestor corresponds to the node defining that branch. By aggregating the adjusted PHS p-values strictly within a specific clade, our approach could theoretically provide a statistical measure of branch support tailored to the irreversible constraints of CRISPR-Cas9 lineage tracing. Because this approach relies on analytical probabilities rather than sequence resampling, it could potentially offer an advantage over traditional bootstrapping, which is often sensitive to the limited sequence lengths typical of CRISPR-Cas9 target arrays.

## Supporting information

Supplemental spreadsheet KPTracer_summary

## Acknowledgments

We thank the reviewers for their valuable feedback that significantly improved the quality of this paper.

## Supplementary Material

## A Handling Model Uncertainty and Incomplete Reconstructions

In this section, we address some practical aspects in applying the cPHS statistic. Specifically, we discuss how to estimate unknown model parameters required for the computation; how to handle cases where only the reconstructed tree topology is available without internal node sequences; and how to prevent saturation of the cPHS score in reconstructed trees with very few homoplasies.

### A.1 Unknown model parameters and birth times

As discussed in the main text, our PHS test statistic depends on two model parameters: the mutation rate *λ* and the collision probability *q*. In addition, in terms of the given tree, it depends on the birth times of the inner nodes. In this section, we discuss their estimation.

Let 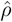 be the fraction of mutated characters in the sequences at the observed cells. Then, as in Jones et al. (2020), the mutation rate *λ* can be accurately estimated from 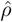 via the plug-in estimate 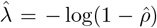 see (1). The collision probability *q*, in contrast, cannot typically be accurately estimated given the result of a single or few clones, since we do not know when up in the tree an observed mutation occurred. However, this parameter is constant for a fixed experimental system (Jones et al., 2020; Seidel and Stadler, 2022). It is thus possible to estimate it from single-cell RNA-seq data of many clones or even from bulk RNA-sequencing of the target sites, as done in Jones et al. (2020) and Seidel and Stadler (2022). In our simulations, we assume knowledge of *q*, and estimate *λ* from the observations. Finally, the birth times *τ* of the inner nodes are estimated using the maximum-likelihood-based branch length estimator proposed in Prillo et al. (2026). In particular, this estimator lower-bounds the minimal branch length for a mutationless edge.

### A.2 Reconstructed topology *T* without internal node sequences *S*

In the main text, it was assumed that the output of a given algorithm is a reconstructed full tree *T S*, which includes both the tree topology *T* and the internal node sequences *S*. However, some algorithms output only a reconstructed tree topology *T* without a corresponding *S*. Given *T* and *S*_obs_, there is more than one possible set of internal node sequences *S* consistent with the non-modifiability model assumption. Since to compute the cPHS score requires also the set of internal sequences, the cPHS score “induced” by only a reconstructed topology is not uniquely defined.

Similarly, the hypothesis testing (3) with either the parsimony or the likelihood distance is also not well-defined. In this section, we propose a procedure to reject candidate topologies w.r.t. *the parsimony distance*, even without internal sequences.

We first present the procedure, and then its justification. Assuming we are given only a candidate topology *T* without reconstructed internal sequences, we first compute the solution to the small MP problem *S*_MP_, under the non-modifiability constraint. This is described in Appendix C and Alg. 2. Next, we compute the test statistic cPHS (*T S*_MP_). If our test rejects *T S*_MP_, then we reject the topology *T* itself.

The justification of this procedure is based on the following lemma, whose proof appears in Appendix E. It states that the solution to the small MP problem maximizes the cPHS score among all the possible sets of sequences that comply with the non-modifiability constraint and coincide with the observed sequences at the leaves.

#### Lemma 2.

*Let T be a fixed tree topology, and let S*_*n*_ *be a given set of sequences at its leaves with no missing data. Denote by S*_*MP*_ *a solution of the small MP problem. Then for any set of sequences S at all tree nodes that complies with the non-modifiability constraint and whose leaves coincide with S*_*n*_, *it holds that*

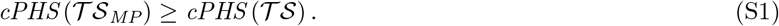

**Figure S1:**
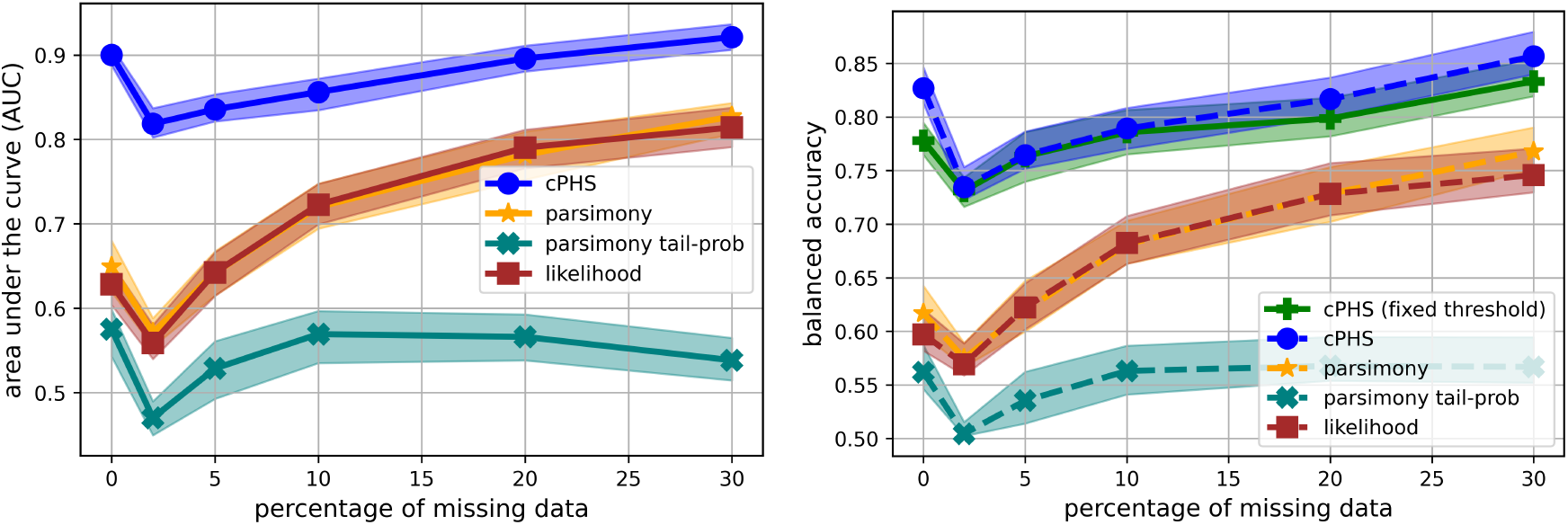
AUC values (left) and balanced accuracy (right) of several test statistics for the hypothesis testing in (3), with the same setting as in Figures 3 and 4 at *k* = 30 and *m* = 32. The x-axis is the total proportion of missing data *p*_miss_.

Suppose that our procedure rejected *T S*_MP_. Assuming our test result is correct, *T S*_MP_ satisfies hypothesis ℋ_1_ of (3), namely *T S*_MP_ > (1 + *ϵ*) · *M* (*T*_GT_*S*_GT_). Hence, by definition of the small MP problem, *T S*_MP_ satisfies ℋ_1_ of (3) for any *S*. The topology *T* should thus be rejected, as required.

### A.3 Very few homoplasies

In an extreme case where the reconstructed full tree has very few homoplasies, it may occur that all sorted scores satisfy

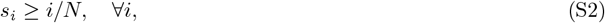

In this case, using (13) leads to a saturation in cPHS, outputting the maximal possible value of 1. This precludes a finer comparison of different trees that both obtain cPHS = 1. To overcome this issue, we exclude leaf pairs with no homoplasies, {*i* : *s*_*i*_ = 1}, before taking the minimum. Formally, Eq. (13) is modified to

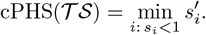

## B Missing Data

In the main text, we assumed a complete (i.e., fully observed) data matrix *S*_*n*_. In practice, however, some entries in *S*_*n*_, corresponding to characters at the tree leaves, may be unknown. The CRISPR-Cas9 evolution model has two distinct mechanisms of missing data. The first is stochastic missingness, which arises from imperfections in the sequencing process, leading to randomly missing entries in *S*_*n*_. The second is heritable missingness, which occurs when characters at internal nodes are absent due to transcriptional silencing or double Cas9 resection. These missing characters behave analogously to mutations: their unknown state is non-modifiable and is inherited by all descendant nodes, including the leaves. Let *p*_sto_ and *p*_her_ denote the rates of stochastic and heritable missingness in *S*_*n*_, respectively. Assuming independence, the overall missing data rate is then given by *p*_miss_ = *p*_sto_ +*p*_her_ − *p*_sto_ ·*p*_her_.

To deal with the presence of missing data, two minor modifications are required in the definition of the cPHS measure. Let *u* and *v* be a pair of leaf nodes. First, we revise the definition of phs_*i*_(*u, v*) in Eq. (8) as follows: if the *i*-th character of either *u* or *v* is missing, then phs_*i*_(*u, v*) = 0. Second, we modify Eq. (11) in Step 1 of the cPHS computation procedure by replacing *k* with *k*_eff_(*u, v*), defined as the number of characters that are non-missing in both *u* and *v*.

As illustrated in Figure S1, the adapted cPHS performs well in the presence of missing data, and in particular, it retains its advantage over the other test statistics. Interestingly, the performance of cPHS, as well as that of the other test statistics, improves as *p*_miss_ increases. This trend, however, may be attributed to a technical factor: the balance between accurately and inaccurately reconstructed trees shifts with increasing *p*_miss_. Specifically, as *p*_miss_ increases, the reconstruction task becomes more challenging, leading to a higher proportion of inaccurate trees produced by the algorithms. It is reasonable to assume that this shift is the primary driver of the observed improvement in the test statistics’ performance.

## C Small and Large Parsimony Problems

Let us recall the definitions of the small and large maximum parsimony (MP) problems Felsenstein (2004, Chapters 2 and 4).

### Definition 7

(Small MP). In the small MP problem, given a tree topology *T* and observed sequences at the leaves *S*_*n*_, the goal is to reconstruct the sequences at the ancestral (internal) nodes such that the resulting tree has the smallest number of mutations:

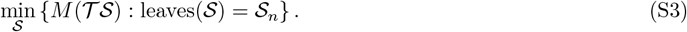

Whereas small MP optimizes only over the internal sequences for a fixed topology, large MP additionally optimizes over the tree topologies:

### Definition 8

(Large MP). In the large MP problem, given the observed sequences at the leaves *S*_*n*_, the goal is to reconstruct a full tree with the smallest number of mutations:

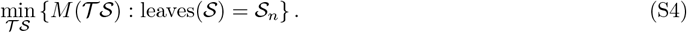

The small MP problem can be solved efficiently by the Sankoff algorithm (Sankoff, 1975). The non-modifiability constraint can be forced by setting the substitution matrix accordingly. The resulting algorithm, presented in Alg. 2, is simple, and bears a resemblance to Fitch algorithm (Fitch, 1977). In the absence of missing data, the solution to the small MP problem under the non-modifiability constraint is unique (Sashittal et al., 2023). The large MP problem, in contrast, is NP-hard in general (Felsenstein, 2004), and remains so under the non-modifiable model (Day, Johnson, and Sankoff, 1986; Sashittal et al., 2023).

## D Parsimony Tail Probability

In this section, we derive the parsimony distribution conditional on a given candidate tree topology, where the inner sequences are random. The distribution is calculated recursively along the tree topology Let *w* be an unmutated node in the tree, and denote the number of mutations at the *i*-th character in the subtree whose root is *w* by 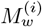. If *w* is a leaf, then no mutation can occur from *w* downwards, namely 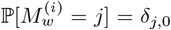 . Next, consider the case where *w* is an inner node. Denote its children by *u* and *v*, and the edge length from *w* to *u* and to *v* by *τ*_*w,u*_ and *τ*_*w,v*_, respectively. Recall that the edge length is the time elapsed between the birth time of a node and its child.

Let *U* and *V* be the events of mutation occurrence at the *i*-th character from *w* to *u* and to *v*, respectively. Then 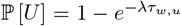 and 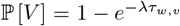. In addition, by the Markovian property of the generative process, *U* and *V* are independent given that *w* is unmutated. By the law of total probability,

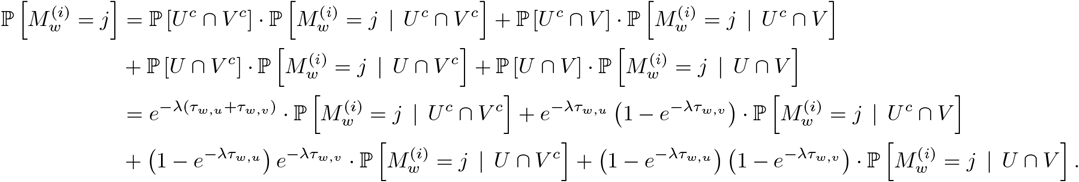

If both *u* and *v* are un mutated (namely under 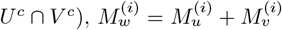. As 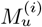 and 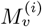 are independent, 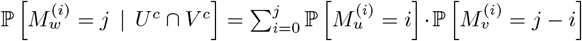. If *v* is mutated but not 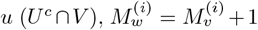 due to the non-modifiability of the evolution process. Hence, 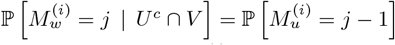. Similarly, 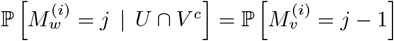. Finally, if both *u* and *v* are mutated, 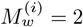 due to non-modifiability,

### Algorithm 2

Solution to the small MP problem under non-modifiable model

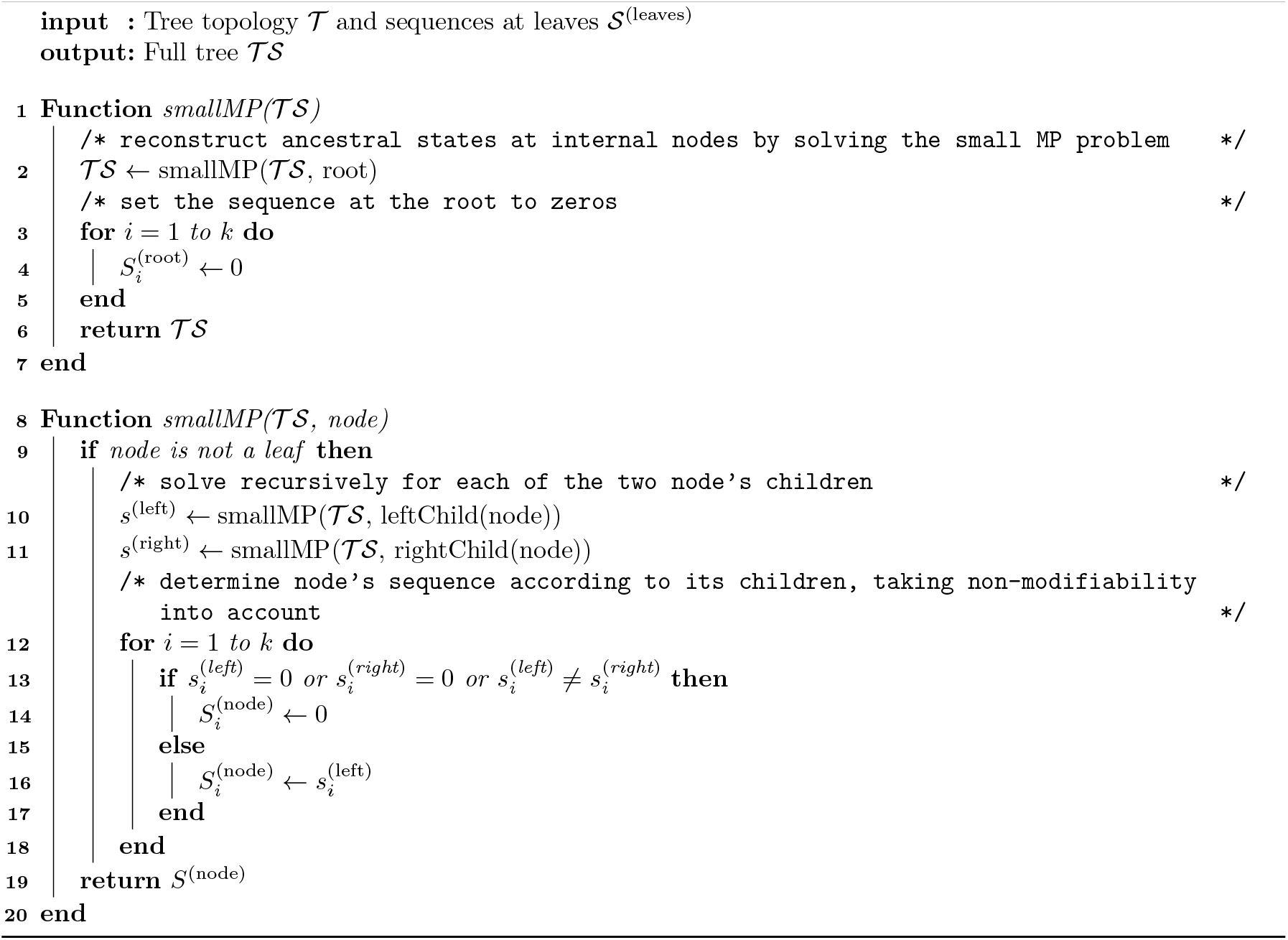

and thus 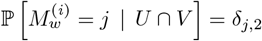 . Putting everything together, we conclude

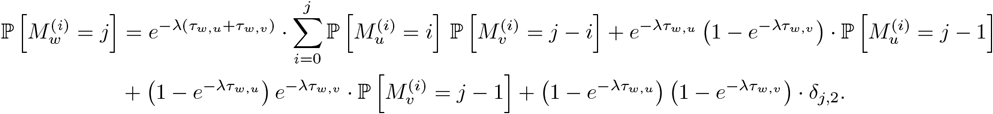

This recursive equation allows us to compute the parsimony distribution at a node *w* at the *i*-th character, 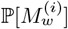, as a function of its children distributions, 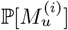 and 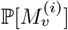. The distribution of the total parsimony (sum of mutations at all characters) of a tree topology *T* with random sequences *S* is given by convolving the individual distributions 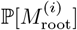,

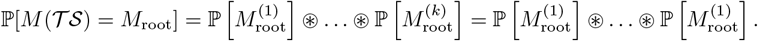

The second equality follows from the assumption of uniform mutation rate; see Remark 4. Finally, the parsimony tail probability for a given tree topology *T* is ℙ[*M* (*T S*) ≤ *M*_root_].

## E Proofs of Lemmas 1 and 2

*Proof of Lemma 1*. Let *p*_*u,v*_ = ℙ[phs_1_(*u, v*) = 1], where the probability is over the random sequences *S*_GT_. Then, by the definition of homoplasy, with *w* = LCA(*u, v*),

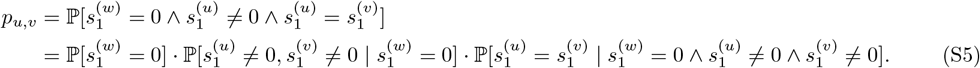

Let us analyze each of the three terms in the RHS of (S5). The first expression is simply the probability that the first character in *s*^(*w*)^ is unmutated, which is given by

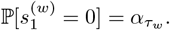

Regarding the second term, since characters at node *u* and at node *v* evolve independently, conditional on their LCA *w* being unmutated,

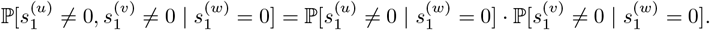

The probability that the character at *u* is unmutated given that *w* is unmutated is 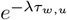 where *τ*_*w,u*_ = *τ*_*u*_ − *τ*_*w*_ = 1− *τ*_*w*_, where the second equality follows from the fact that *u* is a leaf. Hence, 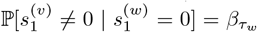. As a similar argument holds for *v*,

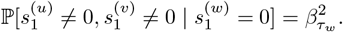

As for the third term, since a mutated character is *j* with probability *q*_*j*_, the probability of a collision (same character in both nodes) is given by

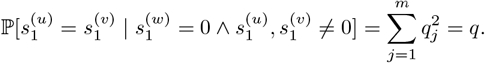

Inserting all of these into (S5) yields 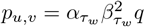. The Binomial distribution in (10) of the lemma follows from the fact that characters at different locations evolve independently with the same parameters *λ* and *q*.

*Proof of Lemma 2*. Let *S* be a set of sequences that coincides with *S*_GT_ at the leaves. Assume that for any pair of leaves *u, v*,

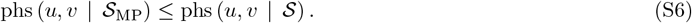

Then, by (11), *s*(*u, v* | *S*_MP_) ≥ *s*(*u, v* | *S*). Therefore, upon computing (10), (11), sorting the *s*-values, adjusting them by (12) and taking the minimum as in (13), (S1) of the lemma follows. Hence, we now prove that (S6) indeed holds.

Let *u, v* be a pair of leaves, and let *w* = LCA(*u, v*). For simplicity, we focus on the first character location. Since characters evolve identically and independently, the same argument holds for all the characters.

Let *z*_1_, *z*_2_ be the immediate children of *w*. There are four generic possible scenarios for 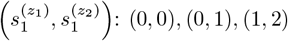 and (1, 1). All the other possibilities are equivalent to one of these four. Under the non-modifiable model, 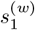 must be 0 in the first three cases. The only freedom of choice left to the inner sequences reconstruction algorithm is in the case of 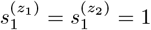, which we now analyze in detail. In this case, all the descendants of *w* have 1 in their first character. Hence, assigning 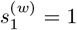 minimizes the PHS of all leaf pairs that are descendants of *w*, including *u* and *v*. It is left to show that in *S*_MP_, *w* is indeed assigned with 1 in this case.

Let us analyze the effect of 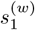 on the total number of mutations, *M* (*T S*_MP_). Assigning 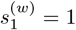 increases the total number of mutations by either 0 (if the sibling of *w* has 1) or 1 (if it has 0). Assigning *w* with 0, on the other hand, increases the total number of mutations by at least 2, as two changes are required from 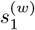 to 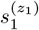 and to 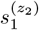. Hence, the small MP solution assigns 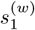 with 1, as required.

In the main text, we assume a uniform mutation rate *λ* across all *k* characters. However, as claimed in Remark 4, our results can be easily generalized. This is formalized in the following claim.

**Figure S2:**
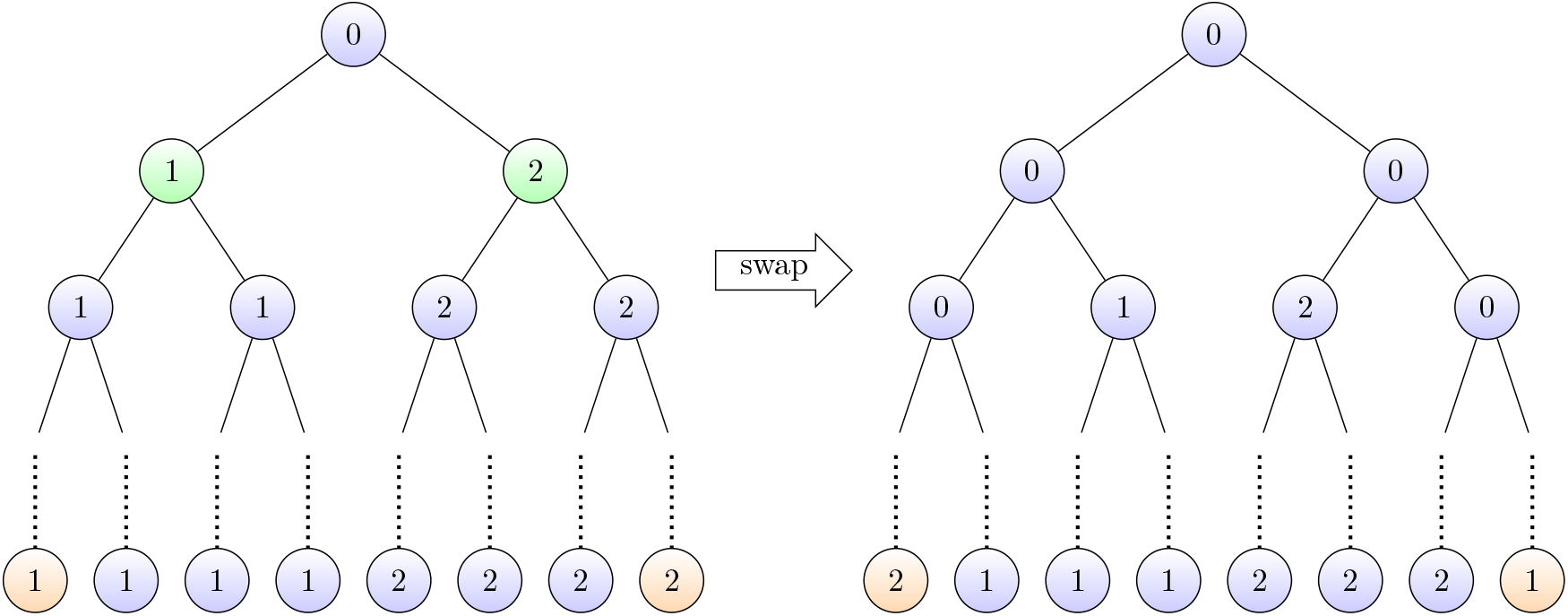
Effect of a leaf swap. Left: a ground-truth tree where mutations occurred early in the tree (at the green nodes). As a result, each of its two subtrees has a different mutated state. Right: a reconstructed tree which differs from the GT tree by a single swap (the swapped leaves are marked in orange). The inner node sequences are reconstructed to satisfy the non-modifiability assumption.

### Claim 1

(non-uniform mutation rates). *Under the assumptions of Lemma 1 but with non-uniform mutation rates λ*_*i*_ *for each i* ∈ [*k*], *the probability distribution of k* · *phs*(*u, v*) *is Poisson Binomial with*

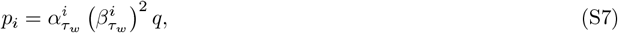

*where* 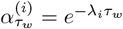.*and* 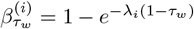.

*Proof*. By following the steps of the proof of Lemma 1, we get that 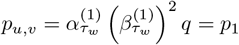. More generally,

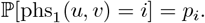

The claim follows from the independence of different characters and the definition of the Poisson Binomial distribution.

## F Illustration of a Leaf Swap, and its Effects on PHS and Parsimony

Let us illustrate the difference in tPHS and parsimony values following a leaf swap. For simplicity, the sequence length in our example consists of a single character, *k* = 1. Suppose that in the ground-truth tree, the immediate offspring of the root mutated to two different states: the left branch mutated to 1 and the right branch to 2. As a result, the state of all the leaves in the left subtree is 1, and it is 2 in the right subtree; see Figure S2(left) for an illustration. Now, suppose that a tree reconstruction algorithm made a single error of swapping the leftmost and rightmost leaves; see Figure S2(right). In this example, there are no homoplasies in the ground-truth tree, and all its leaf pairs have phs(*u, v*) = 0. In the reconstructed tree, in contrast, a non-negligible fraction of leaf pairs have a PHS of 1. As stated in Lemma 10 below, in this case the increase in tPHS scales as the number of leaf pairs, which is roughly 4^*d*^. This is comparable to the tPHS standard deviation over random inner sequences; see Lemma 4. Hence, PHS is sufficiently powerful to identify the reconstructed tree as incorrect. In contrast, as presented in Lemma 5, the parsimony increase is only linear in *d*. This is negligible compared to the parsimony standard deviation over random inner sequences, which scales as the number of leaves, *n* = 2^*d*^; see Lemma 3. As a result, the parsimony measure is unable to distinguish between the correct and the incorrect tree.

## G Auxiliary Lemmas for Theorem 1

The proof of Theorem 1 makes use of four auxiliary lemmas, outlined below. In the following, ℒ (*T*) denotes the set of leaves of a tree *T*.

### G.1 Mean and variance of parsimony and tPHS

We start with two lemmas, regarding the mean and variance of parsimony and tPHS. The first lemma provides exact expressions for the parsimony mean and variance, assuming a homogeneous tree. The second lemma provides upper bounds for the mean and variance of tPHS in general trees, not necessarily homogeneous. Their proofs appear in Sections I.1 and I.2, respectively.

#### Lemma 3.

*Let T*_*GT*_ *be a homogeneous tree of depth d, whose root node is unmutated. Let D be the distribution of its set of sequences S*_*GT*_ *of length k, generated according to the non-modifiable model of Section 2.1 with mutation rate λ. Then, taking expectation over D, for α*_*d*_ = *e*^−*λ/d*^ ≠ 1*/*2,

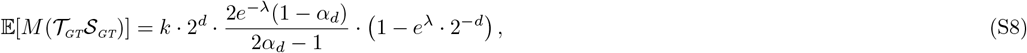

*and*

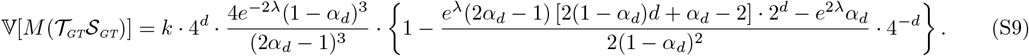

#### Lemma 4.

*Con*_(_*sid*_)_*er the same setting as in Lemma 3, but with an arbitrary tree T*_*GT*_ *(not necessarily homogeneous). Denote by* 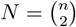 *its number of leaf pairs. Then, taking expectation over D*,

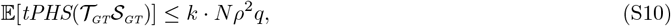

*and*

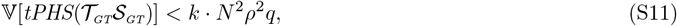

*where q is the collision probability and ρ is the probability of observing a mutation at a leaf given in* (1).

Comparing Lemmas 3 and 4 reveals an important difference between parsimony and tPHS: the parsimony does not depend on the collision probability *q*. In contrast, the mean and variance of tPHS are linear in *q*. As a result, in general, a small value of *q* - which is directly related to the number of homoplasies - increases the statistical power of cPHS; see (16).

*Remark* 7. Lemma 3 assumes that *α*_*d*_ 1*/*2. In the case *α*_*d*_ = 1*/*2, the mean and variance read

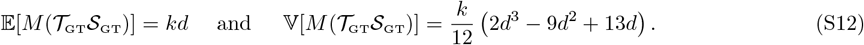

This can be easily verified by plugging *α*_*d*_ = 1*/*2 in the proof of the lemma, or by taking the limit *α*_*d*_ → 1*/*2 in Eqs. (S8) and (S9).

### G.2 The effect of a leaf swap on parsimony and on tPHS

We present two lemmas regarding the mean change in parsimony and tPHS, following a swap of two leaves *u, v* from different sides of the tree (see Section 5). For convenience, the lemmas consider the change in parsimony and tPHS at a single character. Since both parsimony and tPHS are additive in the sequence length *k*, the mean change in these quantities is simply *k* times that of a single character.

#### Definition 9.

[per-character parsimony] Given a full tree *T S*, for any *i* ∈ [*k*] denote by *M*_*i*_(*T S*) the number of mutations in the *i*-th character across all edges of the tree.

#### Definition 10.

[per-character tPHS] Given a full tree *T S*, denote by tPHS_*i*_(*T S*) the number of leaf pairs with a PHS of 1 at their *i*-th character, *i* ∈ [*k*]:

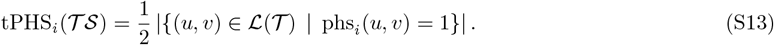

The factor of 1/2 compensates for double summation of pairs. Note that by definition, the parsimony and tPHS of a tree are the sum of its per-character parsimony and tPHS scores: 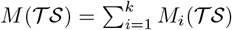 and 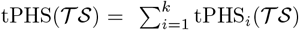, respectively. In addition, tPHS_*i*_(*T S*) = *k* ·∑ _(*u,v*)_ phs_*i*_(*u, v*).

Let us now present the lemmas. Their proofs appear in Appendices I.3 and I.4, respectively.

#### Lemma 5.

*Consider the same setting as in Lemma 3. Further suppose λ* ≤ *d/*2. *Let u, v* ∈ ℒ (*T*_*GT*_) *be two leaves whose LCA is the tree root. Then, taking expectation over D, for any i* ∈ [*k*], *the expected parsimony change in the i-th character following the leaf swap u* ↔ *v is given by*

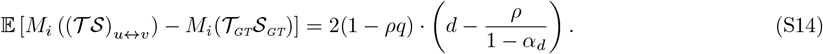

#### Lemma 6.

*Assume the conditions of Lemma 5. Then, taking expectation over D, the expected tPHS change in the i-th character following the leaf swap u* ↔ *v satisfies*

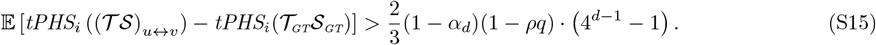

Lemmas 5 and 6 demonstrate a key difference between parsimony and tPHS: Following the leaf swap described above, the mean tPHS value increases *exponentially* with respect to the tree depth *d*, whereas on average, parsimony increases only linearly.

## H Proof of Theorem 1

*Proof*. First, we prove the lower bound (16) on the normalized change in tPHS due to a leaf swap. Let *α*_*d*_ = *e*^−*λ/d*^. For any *d* ≥ 2*λ*,

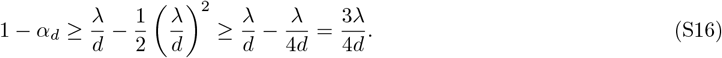

Inserting this inequality into (S14) of Lemma 6 yields

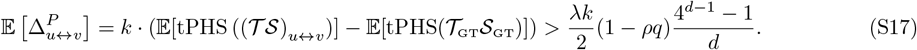

Next, we apply Lemma 4. Since the tree is homogeneous, its number of leaves is *n* = 2^*d*^. Now, for *d* ≥ 2, 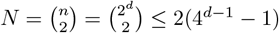. Hence, Eq. (S11) of the lemma gives

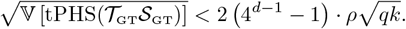

Equation (16) follows by combining this with (S17).

Next, we prove the upper bound (17) regarding the normalized change in parsimony. Combining the additivity of the parsimony in the sequence length *k* with (S14) of Lemma 5 regarding the change in parsimony at a single character, gives that

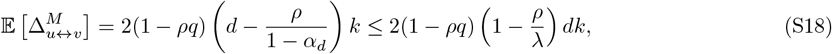

where the inequality above follows from 1 − *α*_*d*_ ≤ *λ/d*. Next, we apply Lemma 3. For a sufficiently large *d, α*_*d*_ > 1*/*2, and thus satisfies the assumption of the lemma. Further, for a sufficiently large *d*, the expression in the curly brackets in the RHS of (S9) is larger than 2*/*3. Hence,

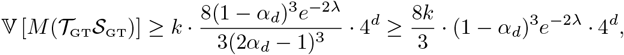

where the second inequality above follows from *α*_*d*_ ≤ 1. Combining this with Eqs. (1) and (S16) yields

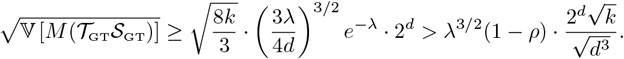

Equation (17) follows by combining this with (S18).

**Figure S3:**
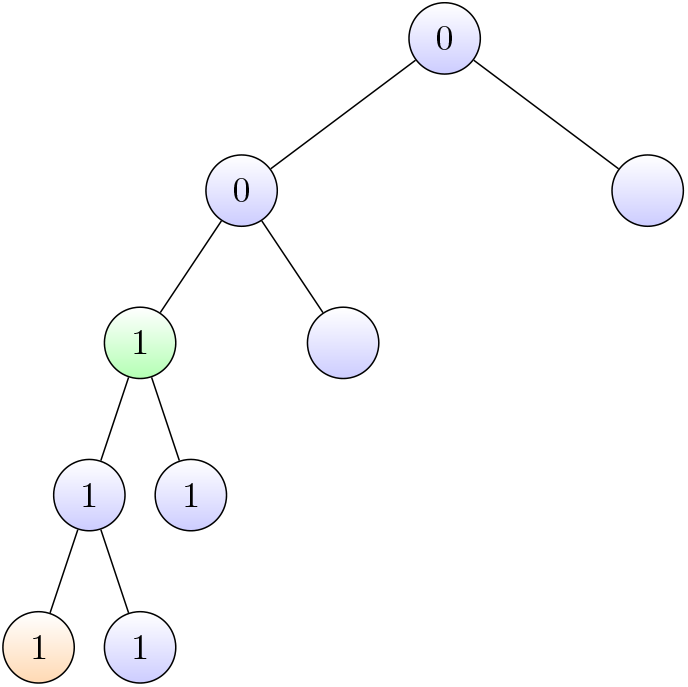
Illustration of the origin of an observed mutation (Def. 11) at a node *u* (colored in orange) with *k* = 1. The most distant node with the same mutation, *A*_1_(*u*), is colored in green node. Their edge distance is *E*_1_(*u*) = 2.

## I Proofs of Lemmas 3, 4, 5 and 6

To prove the lemmas, we first introduce some definitions. Recall that ℒ (*T*) denotes the set of leaves of a tree *T*. For convenience, for any leaf *z* define phs(*z, z*) = 0. In addition, we define the following three quantities regarding the origin of observed mutations, that will be used in our proof. See Fig. S3 for their illustration.

### Definition 11.

Let *T S* be a full tree whose sequences were generated by the non-modifiable mutation process of Section 2.1. Let *u* ∈ ℒ (*T*) be a leaf in the tree. For each character location *i*, we denote by *A*_*i*_(*u*) the most distant ancestor of *u* that had a mutation in the *i*-th character. If either 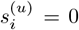 or 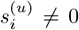 but its father node was unmutated, then we define *A*_*i*_(*u*) = *u*. In addition, we denote by *E*_*i*_(*u*) the number of edges from *A*_*i*_(*u*) to *u*, and by 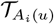 the subtree whose root is *A*_*i*_(*u*).

Note that by non-modifiability of the mutation model, 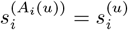.

### I.1 Proof of Lemma 3 (parsimony mean and variance)

The proof of Lemma 3 makes use of the following auxiliary lemma, to be proved shortly.

#### Lemma 7.

*Assume the same setting as in Lemma 3. Let T*_*l*_ *be a subtree of T*_*GT*_ *of depth l* ≤ *d. Let D*_*l*_ *be the distribution of its set of sequences S*_*l*_ ⊆ *S*_*GT*_, *conditioned on the event that the sequence at the root node of T*_*l*_ *is unmutated. For any i* ∈ [*k*], *denote the parsimony in the i-th character of the subtree T*_*l*_*S*_*l*_ *by* 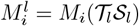, *Then, taking expectation over D*_*l*_,

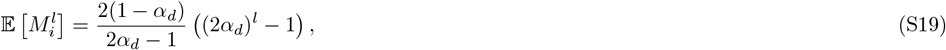

*and*

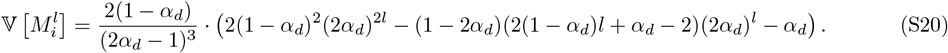

*Proof of Lemma 3*. By the linearity of expectation, 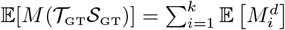, where 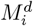 is defined in Lemma 7. As characters at different locations *i* ∈ [*k*] evolve independently from each other, 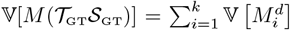. (The lemma follows by taking *l* = *d* in Eqs. (S19) and (S20) of Lemma 7.

*Proof of Lemma 7*. Without loss of generality, we may assume that *i* = 1. To simplify notation, we omit the subscript 1 from 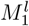. We prove the claim by induction on *l*. For *l* = 0, the subtree consists of only the root, and thus *M*_0_ = 0 deterministically. As the RHS of Eqs. (S19) and (S20) vanish at *l* = 0, the induction base is proved.

**Figure S4:**
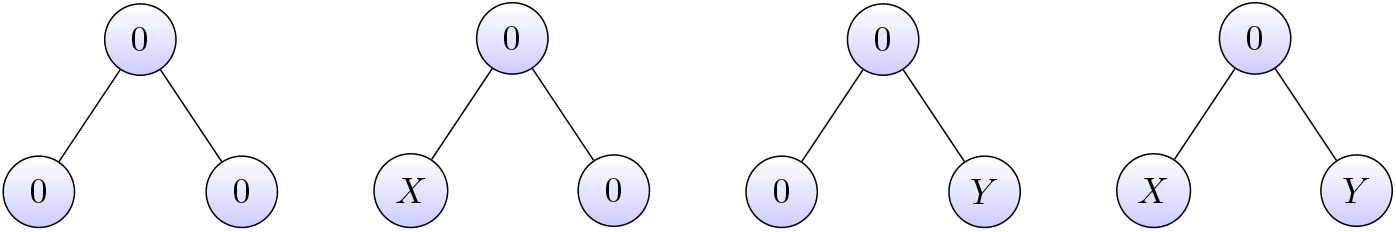
Given an unmutated root, there are four possible scenarios for the states of its two children: (i) no mutation in both branches, (ii) a mutation only in the left branch; (iii) a mutation only in the right branch, and mutations in both branches. Here *X, Y* ∈ [*m*].

Next, assume that the induction hypothesis holds, namely (S19) and (S20) hold for some *l* − 1 ≥ 0. We now prove it holds for *l*. Let *E*_*l*_ = E[*M* ^*l*^] and *V*_*l*_ = V[*M* ^*l*^] be the mean and variance, respectively, of the number of mutations in a subtree of depth *l* whose root has an unmutated sequence. The mean and variance depend only on *l* and not on the specific subtree, because all subtrees of a homogeneous tree are homogeneous as well. Let 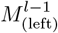 and 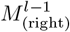 be the parsimony scores of the subtrees at the left and right branches of *T*_*l*_*S*_*l*_, respectively, each conditioned on the event that the corresponding subtree roots are unmutated. For future use, note that by the Markovian property of the generative process, 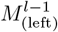 and 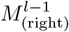 are independent given that the root is unmutated.

To analyze *E*_*l*_ and *V*_*l*_, we split into cases. Let *A* ∈ {0, 1, 2 }be the number of mutations from the root of *T*_*l*_ to its immediate children; see Figure S4 for an illustration of the different possible events. We first calculate the mean and variance conditioned on a specific value of *A*. Under the event *A* = 0 (case (i) in Figure S4),

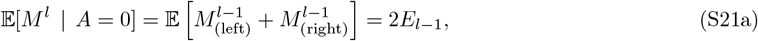

and

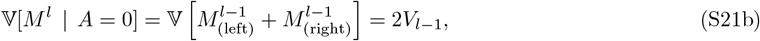

Conditioned on *A* = 1, there are two scenarios: the mutation can be either in the left or in the right branch (cases (ii) and (iii) in Fig. S4, respectively). Due to the non-modifiability of the mutation process, no further mutations can occur in the mutated branch. As a result, 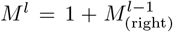 in case of a mutation in the left branch, and 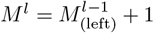 in case of a mutation in the right branch. Since the scenarios of a mutation at the left and at the right branches have equal probabilities,

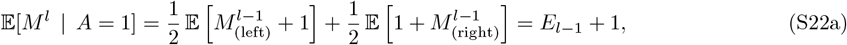

and

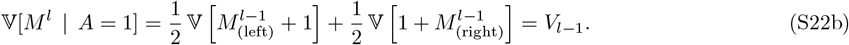

Conditioned on *A* = 2 (case (iv) in Figure S4), due to the non-modifiability of the mutation process,

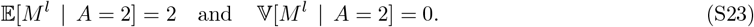

The probability of observing a mutation along a single edge, given that the parent node is unmutated, is 1 − *α*_*d*_. By independence of 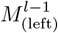 and 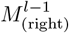, we have that ℙ[*A* = 2] = (1 − *α*_*d*_)^2^, ℙ[*A* = 1] = 2*α*_*d*_(1 − *α*_*d*_), and 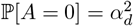. Hence, by the law of total expectation,

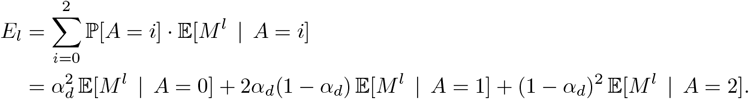

Together with Eqs. (S21a), (S22a) and (S23), we obtain

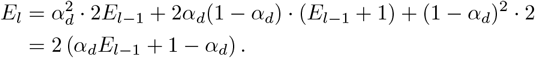

Inserting the induction hypothesis (S19) for *l* − 1 yields that (S19) hold for *l*.

Similarly, by the law of total variance,

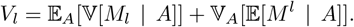

The second term above satisfies

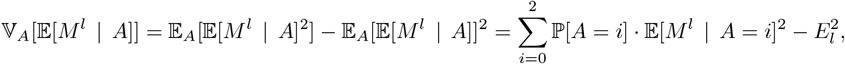

so that

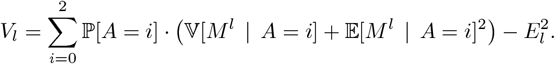

Inserting the expressions derived for the mean and variance conditioned on *A*, Eqs. (S21), (S22) and (S23) give that

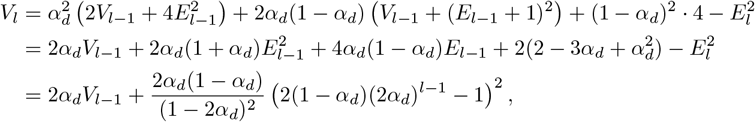

where the last equality follows from (S19). Plugging in the induction hypothesis (S20) at *l* − 1 yields (S20) at *l*.

### I.2 Proof of Lemma 4 (PHS mean and variance)

*Proof*. Let *u, v* ∈ ℒ (*T*_GT_) be a pair of leaves in the tree, and denote their LCA by *w*. Let *τ*_*w*_ ∈ [0, 1) be the birth time of *w*. By Lemma 1, *k* · phs(*u, v*) follows a Binomial distribution with *k* trials and success probability 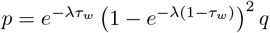. Hence, using (1),

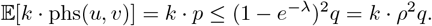

Equation (S10) of the lemma follows from the definition of tPHS (14) and the linearity of expectation.

Next, let us consider the variance. By (10), for a single pair of leaves,

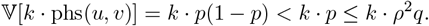

Equation (S11) follows from the fact that 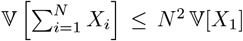 for any collection of identically distributed random variables *X*_*i*_.

### I.3 Proofs of Lemma 5 (mean parsimony change following a leaf swap)

For a fixed tree *T*_GT_, the change in parsimony following a leaf swap between *u, v* is a random variable that depends only on the random quantities *E*_*i*_(*u*) and *E*_*i*_(*v*), defined in Def. 11. Indeed, as the following auxiliary lemma shows, conditional on *E*_*i*_(*u*) and *E*_*i*_(*v*), the change is deterministic. The proof of Lemma 5 follows by combining this result with Lemma 9 below, which characterizes the distribution of *E*_*i*_(*u*). The proofs of these auxiliary lemmas appear in Appendices I.5 and I.6.

#### Lemma 8.

*Let T*_*GT*_*S*_*GT*_ *be a full tree, and let u, v* ∈ ℒ (*T*_*GT*_) *be two leaves whose LCA is the tree root. Then, for any i* ∈ [*k*], *the parsimony change in the i-th character following the leaf swap u* ↔ *v is given by*

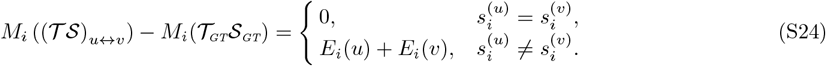

#### Lemma 9.

*Let T*_*GT*_*S*_*GT*_ *be a full tree with a homogeneous topology of depth d, generated according to the non-modifiable model of Section 2.1 with a mutation rate λ. Denote ρ* = 1 − *e*^−*λ*^ *as in* (1), *and α*_*d*_ = *e*^−*λ/d*^. *Let u be a leaf in the tree. For any i* ∈ [*k*], *if the i-th character of u is unmutated*, 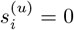, *then E*_*i*_(*u*) = 0. *Otherwise, its distribution is given by*

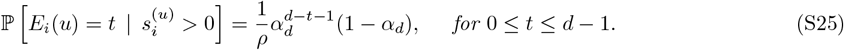

We now prove the lemma.

*Proof of Lemma 5*. Denote 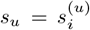 and 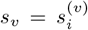. Let Δ = *M*_*i*_ ((*TS*)_*u*↔*v*_)) − *M*_*i*_(*T*_GT_*S*_GT_). The quantity of interest is E[Δ]. To this end, denote the event *A* = {*s*_*u*_ ≠ *s*_*v*_}. Under *A*^*c*^, Δ = 0. Hence,

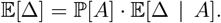

Next, we split *A* into three cases, depending on whether *s*_*u*_ or *s*_*v*_ were mutated, or both:

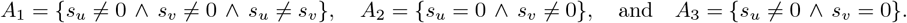

By a symmetry argument, ℙ[*A*_2_ | *A*] = ℙ[*A*_3_ | *A*] and E[Δ | *A*_2_] = E[Δ | *A*_3_]. Hence,

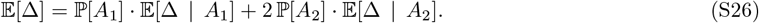

Next, we compute the probabilities in the equation above. Since the LCA of *u* and *v* is the root, which is unmutated by construction, *s*_*u*_ and *s*_*v*_ are independent random variables. We thus obtain

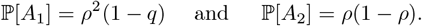

Inserting this into (S26) yields

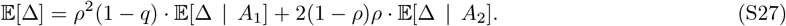

Next, we compute the conditional expectations. Denote *e*_*u*_ = *E*_*i*_(*u*) and *e*_*v*_ = *E*_*i*_(*v*). By combining this with the law of total expectation and (S24) of Lemma 8,

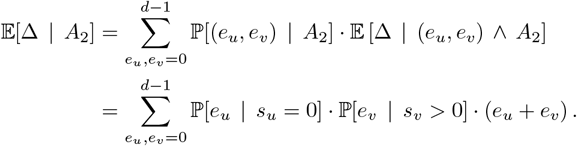

Since *s*_*u*_ = 0, *E*_*i*_(*u*) = 0 follows by definition. Hence,

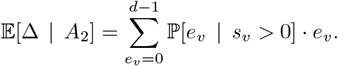

Combining this with (S25) of Lemma 9, we obtain

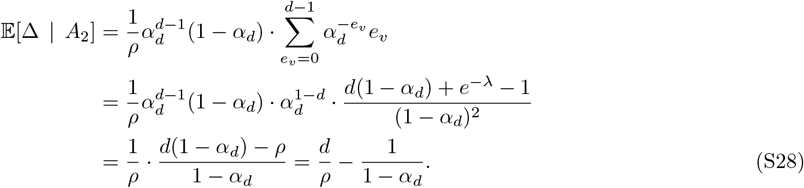

By similar arguments,

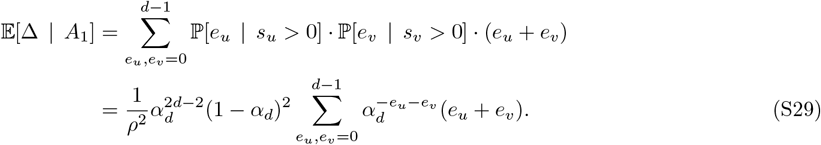

Let us calculate the sum in the RHS of (S29),

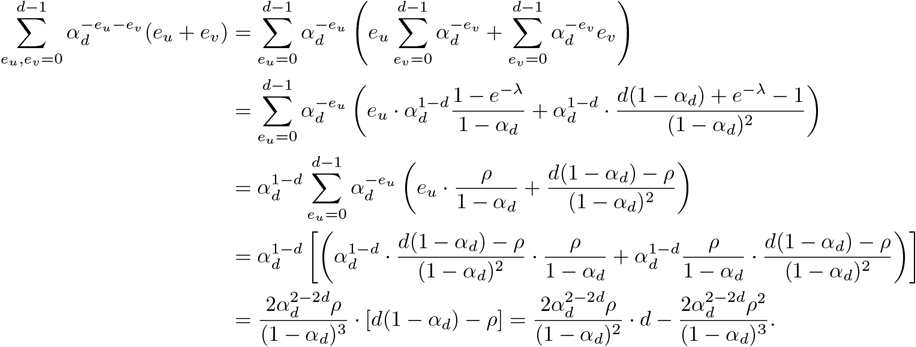

Plugging this into (S29) gives that

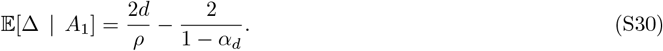

The lemma follows by combining (S27) with (S28) and (S30).

### I.4 Proof of Lemma 6 (mean tPHS change following a leaf swap)

Similar to parsimony, the change in tPHS following a leaf swap between *u, v* is a random variable that depends only on the two random quantities *E*_*i*_(*u*) and *E*_*i*_(*v*), defined in Def. 11. Indeed, as the following auxiliary lemma shows, conditional on *E*_*i*_(*u*) and *E*_*i*_(*v*), the change is deterministic. The proof of Lemma 6 follows by combining this result with Lemma 9 above, which characterizes the distribution of *E*_*i*_(*u*). The proof Lemma 10 appears in Appendix I.5.

#### Lemma 10.

*Let T*_*GT*_*S*_*GT*_ *be a homogeneous full tree, and let u, v* ∈ ℒ (*T*_*GT*_) *be two leaves whose LCA is the tree root. Suppose that either* 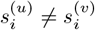 *or* 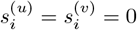. *Then, for any i* ∈ [*k*],

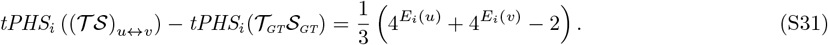

We now prove the lemma.

*Proof of Lemma 6*. Denote 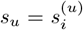 and 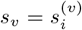. By similar arguments as in the proof of Lemma 5, (S27) holds for Δ = tPHS_*i*_ ((*T S*)_*u*↔*v*_) − tPHS_*i*_(*T*_GT_*S*_GT_) as well. That is,

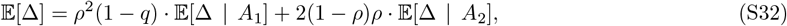

where *A*_1_ = {*s*_*u*_ ≠ 0 ∧ *s*_*v*_ ≠ 0 ∧ *s*_*u*_ ≠ *s*_*v*_} and *A*_2_ = {*s*_*u*_ = 0 ∧ *s*_*v*_≠ 0}. Since *u* and *v* are from different sides of the tree, *s*_*u*_ and *s*_*v*_ are independent, as well as *e*_*u*_ and *e*_*v*_. By combining this with the law of total expectation and (S31) of Lemma 10,

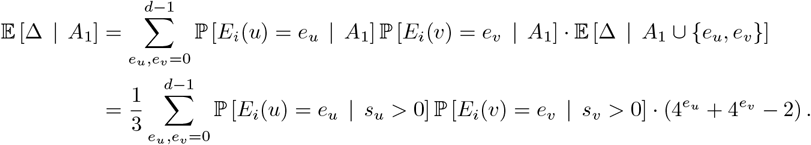

Invoking Lemma 9 gives that

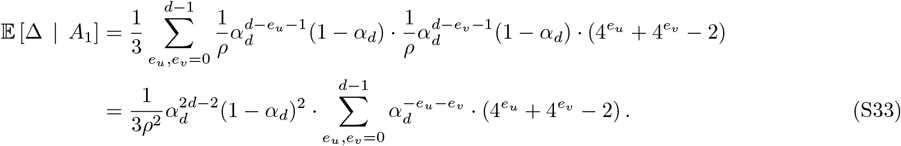

Observe that

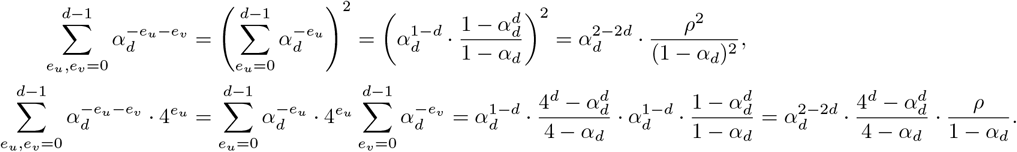

By symmetry, 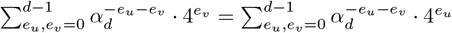. Inserting these expression into (S33) yields

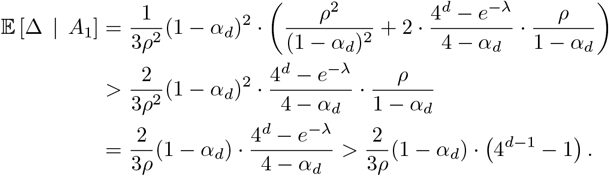

Similarly, by combining the law of total expectation, (S31) of Lemma 10 and (S25) of Lemma 9,

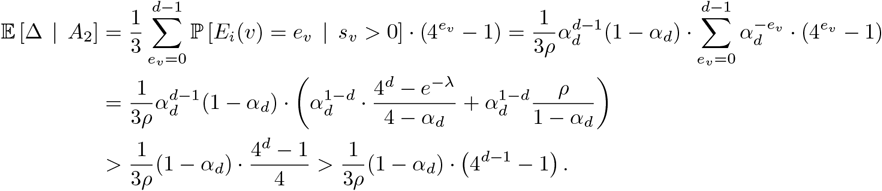

Inserting the last two results into (S32) gives (S14) and completes the proof of the lemma.

### I.5 Proofs of Lemmas 8 and 10 (deterministic parsimony and PHS change following a leaf swap)

The proofs of the lemmas use the following two propositions regarding the quantities defined in Def. 11.

#### Proposition 1.

*Let T S be a full tree, and let u be a leaf in the tree. Let i* ∈ [*k*]. *Then for any* 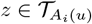,

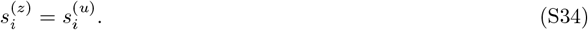

*In addition, let w be the parent of A*_*i*_(*u*). *Then*

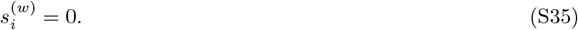

*Proof*. To simplify the notations, for any node *z*, denote 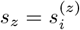. Let 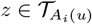. There are two cases to consider: *s*_*u*_ = 0 or *s*_*u*_≠ 0. If *s*_*u*_ = 0, then by non-modifiability all ancestors of *u* are unmutated. Hence, *A*_*i*_(*u*) = *u*, and 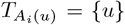. In this case, (S34) trivially holds. Next, suppose *s*_*u*_≠ 0. Since *A*_*i*_(*u*) is an ancestor of *u*, whose character is mutated, by the non-modifiability 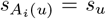. Since any 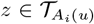 is either *A*_*i*_(*u*) or an offspring of *A*_*i*_(*u*), (S34) follows by the non-modifiability. Finally, (S35) follows from the definition of *A*_*i*_(*u*) as the most distant mutated ancestor of *u*.

#### Proposition 2.

*Let T*_*GT*_*S*_*GT*_ *be a full tree, and let u be a leaf in the tree. Let i* ∈ [*k*], *and denote* 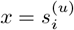. *Let T S be the tree after setting* 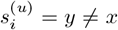 *and correcting for non-modifiability (cf. Def. 6). Then, for all ancestors z of u that belong to* 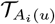 *it holds that in the original tree* 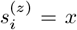, *whereas in the modified tree* 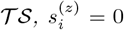. *The sequences at all the other nodes, except u, are the same in T*_*GT*_*S*_*GT*_ *and T S*.

*Proof*. Let *i* ∈ [*k*]. If 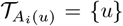, then the lemma holds trivially. Hence, we assume that there is at least one ancestor of *u* in 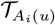, namely 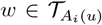 where *w* is the parent of *u*. For convenience, for any node *z* denote 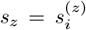. By Proposition 1, *s*_*z*_ = *x* for any 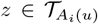 in the original tree *T*_GT_*S*_GT_. In the modified tree *T S*, by definition, *s*_*u*_ = *y*. Since 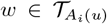, the sibling of *u*, denoted *ū*, satisfies *s*_*ū*_ = *x y*. Hence, by non-modifiability, *s*_*w*_ = 0 in the modified tree. As a result, all ancestors of *u* are corrected from *x* to 0. Since violations of non-modifiability can only occur from the modified leaf and upwards, the proof is complete.

We now prove the lemmas.

*Proof of Lemma 8*. For any node *z*, denote by 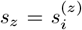 the state of the *i*-th character of *z* in *T*_GT_ *S*_GT_, and by 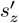 its state in (*T S*_*u*↔*v*_). Denote Δ = *M*_*i*_ ((*T S*)_*u*↔*v*_) − *M*_*i*_(*T*_GT_*S*_GT_).

If *s*_*u*_ = *s*_*v*_, then (*TS*)_*u*_↔_*v*_ = *T*_GT_ *S*_GT_, so Δ = 0 and (S24) holds. Next, we analyze the case *s*_*u*_ ≠ *s*_*v*_. Since *u* and *v* are from different sides of the tree and the root sequence is fixed with unmutated states, we can decompose the parsimony change as

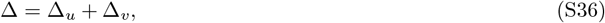

where Δ_*u*_ and Δ_*v*_ are the parsimony changes due to the modification of *s*_*u*_ (while keeping *s*_*v*_ unchanged) and of *s*_*v*_ (while keeping *s*_*u*_ unchanged), respectively.

Let us begin by calculating Δ_*u*_. Denote *a*_*u*_ = *A*_*i*_(*u*) and 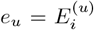. We split into cases: *e*_*u*_ > 0 and *e*_*u*_ = 0. First, suppose that *e*_*u*_ > 0. According to Proposition 2, the only inner nodes whose sequences are modified following the swap are ancestors *z* of *u* that satisfy 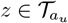. Specifically, they are modified from *s*_*u*_ to 0. Hence, for each such node *z*, there is a new mutation along the branch to the child which is not *u* or one of its ancestors. The number of these nodes *z* is *E*_*i*_(*u*). In addition, since 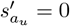, there is no longer a mutation from *a*_*u*_’s parent to *a*_*u*_. Finally, we need to ask if there is a new mutation from *w*, the parent of *u*, to *u*. Since 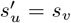, if *s*_*v*_ = 0 then there is no such mutation. Suppose now *s*_*v*_ > 0. Since 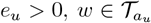, and thus 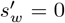. Since 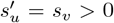, there is a new mutation from *w* to *u*. In total, we obtain that if *e*_*u*_ > 0, then

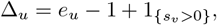

where 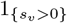 is the indicator of the event {*s*_*v*_ > 0} . Under the assumption *e*_*u*_ > 0, we have 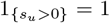. Hence, we can write

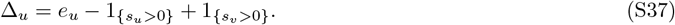

Next, suppose that *e*_*u*_ = 0. Then, by definition of *e*_*u*_, *s*_*w*_ = 0. In addition, since 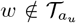. Proposition 2 implies that 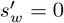. Recall that 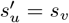. Let us split into three cases: (i) *s*_*u*_ = *s*_*v*_, (ii) *s*_*u*_ = 0 and *s*_*v*_ > 0, and (iii) *s*_*u*_ > 0 and *s*_*v*_ = 0. In case (i), 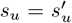 and Δ_*u*_ = 0. In case (ii), 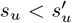 and Δ_*u*_ = 1. Finally, in case (iii), 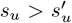 and Δ_*u*_ = 1. It can be thus verified that (S37) holds also in the case *e*_*u*_ = 0.

By similar arguments, one can show that

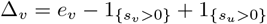

where 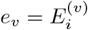 is either 0 or greater than 0. Inserting this together with (S37) into (S36) yields (S24).

In the following proof, we denote the PHS of a leaf pair 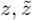 before and after the swap by 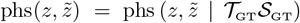 and 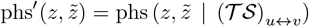, respectively.

*Proof of Lemma 10*. Let *i* ∈ [*k*]. For any node *z*, denote by 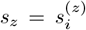 the state of the *i*-th character of *s*^(*z*)^ in *T*_GT_*S*_GT_, and by 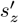 its state in (*T S*)_*u*↔*v*_ . Denote Δ = tPHS_*i*_ ((*T S*)_*u*↔*v*_) − tPHS_*i*_(*T*_GT_*S*_GT_). Further denote *x* = *s*_*u*_ and *y* = *s*_*v*_.

If *x* = *y*, then (*T S*)_*u*↔*v*_ = *T*_GT_*S*_GT_, in which case the LHS of (S31) vanishes. Since, by assumption, *x* = *y* = 0, *E*_*i*_(*u*) = *E*_*i*_(*v*) = 0, and the RHS of (S31) vanishes as well. Next, we analyze the case *x*≠ *y*. For convenience, we denote by 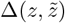 the contribution of a specific pair of leaves 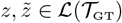 at their *i*-th character to the total PHS change Δ,

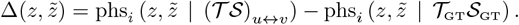

Let 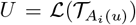 and 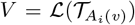 be the sets of tree leaves whose LCA is *A*_*i*_(*u*) and *A*_*i*_(*v*), respectively.

Further let *W* = *U* ∪ *V* and *R* = ℒ (*T*_GT_) \*W* . Then

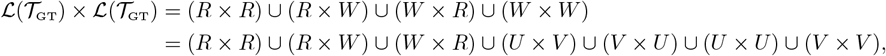

where the union is over disjoint sets. Since for 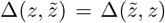 for any 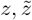, we can decompose the total PHS change as

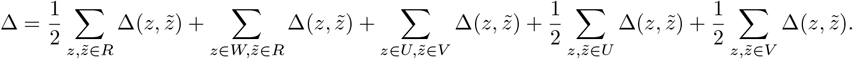

Let *U*^∗^ = *U* \ {*u*} and *V* ^∗^ = *V* \ {*v*}. Then *U* × *V* = (*U*^∗^ × *V* ^∗^) ∪ ({*u*} × *V* ^∗^) ∪ (*U*^∗^ × {*v*}) ∪ ({*u*} × {*v*}), *U* × *U* = (*U*^∗^ × *U*^∗^) ∪ ({*u*} × *U*^∗^) ∪ (*U*^∗^ × {*u*}) ∪ ({*u*} × {*u*}), and similarly for *V* × *V* . Therefore,

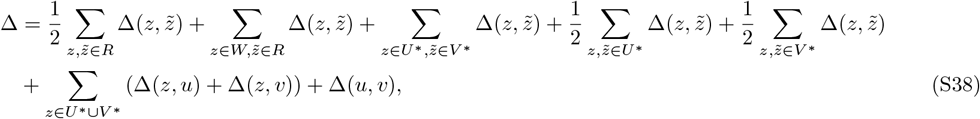

where we used the fact that by definition, Δ(*u, u*) = Δ(*v, v*) = 0.

Let us analyze each of the seven terms in the RHS of (S38). First, we show that the first two terms vanish, since

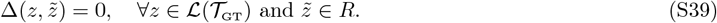

To prove (S39), let 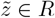, and consider three possible cases for the leaf *z*: either *z* ∈ *R, z* ∈ *U*, or *z* ∈ *V* . If *z* ∈ *R*, then also 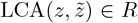. By Proposition 2, the sequences at both 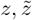, and 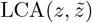 are not affected by the leaf swap, and thus 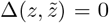. Next, suppose that *z* ∈ *U* . By Proposition 1, 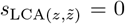. By Proposition 2, also 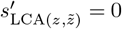, that is 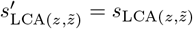. Hence 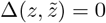 also in this case. The case *z* ∈ *V* is similar to *z* ∈ *U*.

Therefore, (S39) is proved.

Next, we show that the third term in the RHS of (S38) also vanishes, since

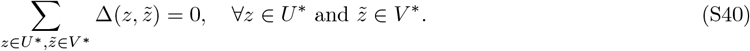

This equality follows trivially by the fact that 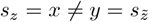 for any *z* ∈ *U*^∗^ and 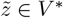.

Next, we show that the fourth term in the RHS of (S38) satisfies

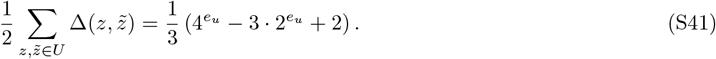

Let 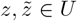. First, let us show that in the unmodified tree, 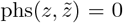. If *x* = 0, then this holds by definition of phs. Next, suppose that *x* ≠ 0. Since 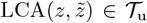, Proposition 1 implies that 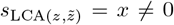. Hence 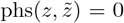 also in this case. Next, to analyze 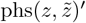 in the modified tree, let *B* be the set of ancestors of *u* which are in *T*_u_. By (S31) of Lemma 10, these ancestors *b* ∈ *B* are the only inner nodes whose states at location *i* get modified. Specifically, they satisfy 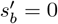. Hence, 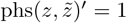 if 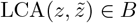, and 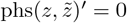 otherwise.

Let us calculate the number of pairs 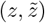 whose LCA is in *B*. Denote 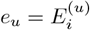, and let 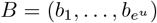, where *b*_*l*+1_ is the parent of *b*_*l*_, and *b*_1_ is the parent of *u*. Observe that *b*_*l*_ has 2^*l*−1^ leaves in each of its branches. However, since *u* is in one of the branches and 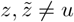, there are only 2^*l*−1^ relevant leaves in one of the branches. The total number of pairs is thus

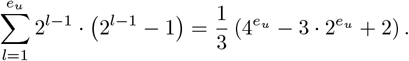

This proves (S41). Similarly, for the fifth term in the RHS of (S38),

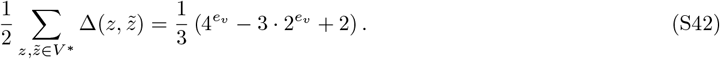

Next, we show that the last term in the RHS of (S38) vanishes,

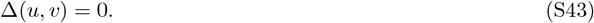

This follows from *s*_*u*_ = *x*≠ *y* = *s*_*v*_, which implies that phs(*u, v*) = phs^*′*^(*u, v*) = 0.

Finally, we show that the sixth term in the RHS of (S38) satisfies

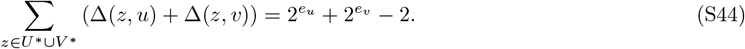

Observe that

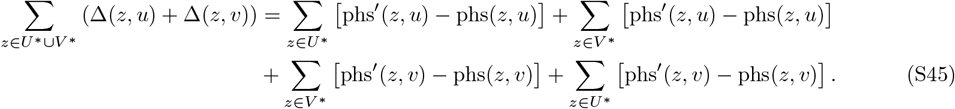

Let *z* ∈ *U*^∗^. By Proposition 1, *s*_LCA(*z,u*)_ = *x*≠ 0, and thus phs(*z, u*) = 0. In the modified tree, 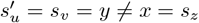 and thus also phs^*′*^(*z, u*) = 0. Next, since *s*_*v*_≠ *s*_*z*_, we have also phs(*z, v*) = 0. In the modified tree, 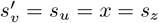. In addition, since *z* and *v* are from different sides of the tree, their LCA is the root which is unmutated. Hence, phs^*′*^(*z, v*) = 1. Now, let *z* ∈ *V* ^∗^. By a similar argument, it follows that phs(*z, v*) = phs^*′*^(*z, v*) = phs(*z, u*) = 0 and phs^*′*^(*z, u*) = 1. Hence, (S45) reads

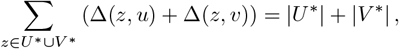

from which (S44) follows. Plugging (S39), (S40), (S41), (S42) and (S43) and (S44) into (S38) proves the lemma.

### I.6 Proof of Lemma 9 (probability distribution of *E*_*i*_(*u*))

*Proof of Lemma 9*. Let 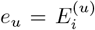. If 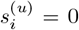, then *e*_*u*_ = 0 by definition. Next, consider the case 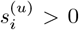. By Bayes’ law,

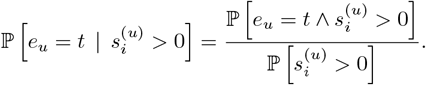

By definition, *A*_*i*_(*u*) is *e*_*u*_ edges above *u*. Let *b* be the parent of *A*_*i*_(*u*). Then *b* is unmutated and *A*_*i*_(*u*) is mutated. Since the depth of *b* is *d* − *e*_*u*_ − 1, we have

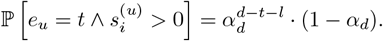

In addition,

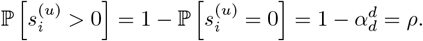

The lemma follows by combining the above three equations.

## J Generation of Random Ground-Truth Tree Topologies

We generated ground-truth trees using the birth-death process implemented in the Cassiopeia package (Jones et al., 2020), specifying parameters for birth and death rates to model realistic lineage structures observed in single-cell tracing experiments. In our simulations, the time between cell divisions follows an exponential distribution with a birth rate initialized to 2. Tree lineages die at times drawn from an exponential distribution with a rate fixed at 0.75. Upon a cell division, its fitness changes with probability 50%. If the fitness changes, then the birth rate is multiplied by 1.1^*z*^ where *z* is drawn from a Normal distribution with mean 0.5 and standard deviation 0.25. The tree grows until the cell population reaches a predefined value, and then it terminates. The times are rescaled such that the end of the experiment is set to have time *τ* = 1. Finally, a set of *n* leaves is subsampled. This subset induces an underlying ground-truth topology, denoted by *T*_GT_.

**Figure S5:**
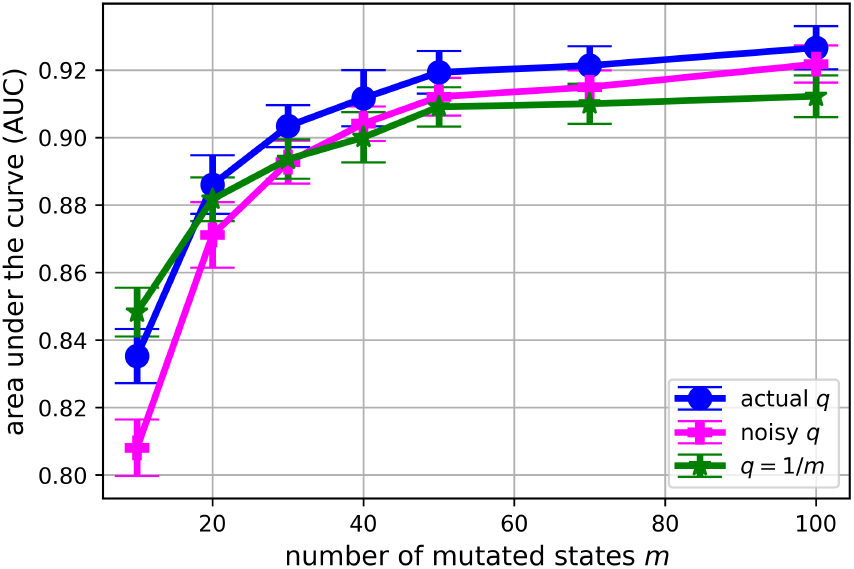
AUC values for cPHS with the true value of *q*, a noisy value of it, and with *q* = 1*/m* that corresponds to uniform mutation probabilities (*q*_*j*_ = 1*/m*). The setting is the same as in Figure 3(right).

## K Additional Simulation Results

As described in Section 3, our PHS-based test statistic requires the knowledge of the model parameter *q*. In our simulations, we assumed that this parameter is known for reasons explained in Appendix A.1. Moreover, empirically, our test statistic is not sensitive to the exact value of *q*. To illustrate this point, we calculated cPHS with three alternatives for the value of *q*: (i) the actual *q*; (ii) a noisy version of it, *q* +*N* (0, 0.3*q*), trimmed to the range [0.1*q*, 1]; and (iii) *q* = 1*/m*, which corresponds to uniform probability distribution of mutation states, namely *q*_*j*_ = 1*/m* for *j* = 1, …, *m*. The result, under the same setting of Figure 3, is depicted in Figure S5. As shown, the performance of cPHS is only slightly affected by an inaccurate estimation of *q*.

In the main text, Figures 2, 3, and 4 summarize the outcomes obtained from 7000 reconstructed trees evaluated under each parameter configuration. Figure S6 presents histograms that distinguish the counts of accurately versus inaccurately reconstructed trees within this sample of 7000. In the upper two panels of Figure S6, the distribution of accurate and inaccurate reconstructions is approximately balanced. In contrast, the lower panel exhibits a pronounced imbalance: the number of inaccurate reconstructions markedly exceeds that of accurate ones. This is due to the fact that the cutoff value *ϵ* in the lower panel is smaller.

Next, Figures S7, S8, S9, and S10 complement Fig. 1 of the main text by showing results for additional distance functions. Specifically, Figures S7 and S8 show the result for the parsimony distance function (*d* = *d*_P_) with the cutoff values *ϵ* = 0.03 and *ϵ* = 0.1, respectively. Figures S9 and S10 show results for the likelihood (*d* = *d*_L_) and triplets (*d* = *d*_tri_) distance functions, with the cutoff values *ϵ* = 0.05 and *ϵ* = 1*/*5, respectively. In all these results, the conclusion of Fig. 1 holds: only cPHS exhibits a good separation between the distribution of accurate and inaccurate trees. As discussed in the main text, these results imply that cPHS can tell whether the parsimony/likelihood of the reconstructed tree is close to the ground-truth one even better than parsimony and likelihood themselves.

Figures S11 and S12 complement Figures 2 and 3 of the main text, by showing results for the triplets and likelihood distance functions, with cutoff values *ϵ* = 0.1 and *ϵ* = 0.03, respectively. Figures S13, S15 and S16 complement Figures 3 and 4 of the main text but show results for additional cutoff values *ϵ* and additional ranges of parameters. Specifically, Figure S13 shows results for the same normalized RF function (*d* = *d*_RF_), but with the cutoff value *ϵ* = 1*/*4. Next, Figure S14 shows the AUPRC (area under the Precision-Recall curve) as a function of the sequence length *k*, generalizing the results presented in the bottom panels of Figure 2. Finally, Figures S15 and S16 show results across ranges of three other model parameters: the number of mutations *m*, the mutation probability at a leaf *ρ*, and the number of observed leaves *n*. Across all these settings, the qualitative conclusions remain consistent with those presented in the main text.

## L Analysis of the KP data set: set up

The KPTracer study deposited character matrices for 85 tumor samples (Yang et al., 2022). We selected the samples for analysis in two stages. First, we applied the quality-control criteria of Yang et al. (2022), which keep a sample if it has at least 5% of cells with a unique indel profile, at least 20% unsaturated target sites, and more than 100 cells.

**Figure S6:**
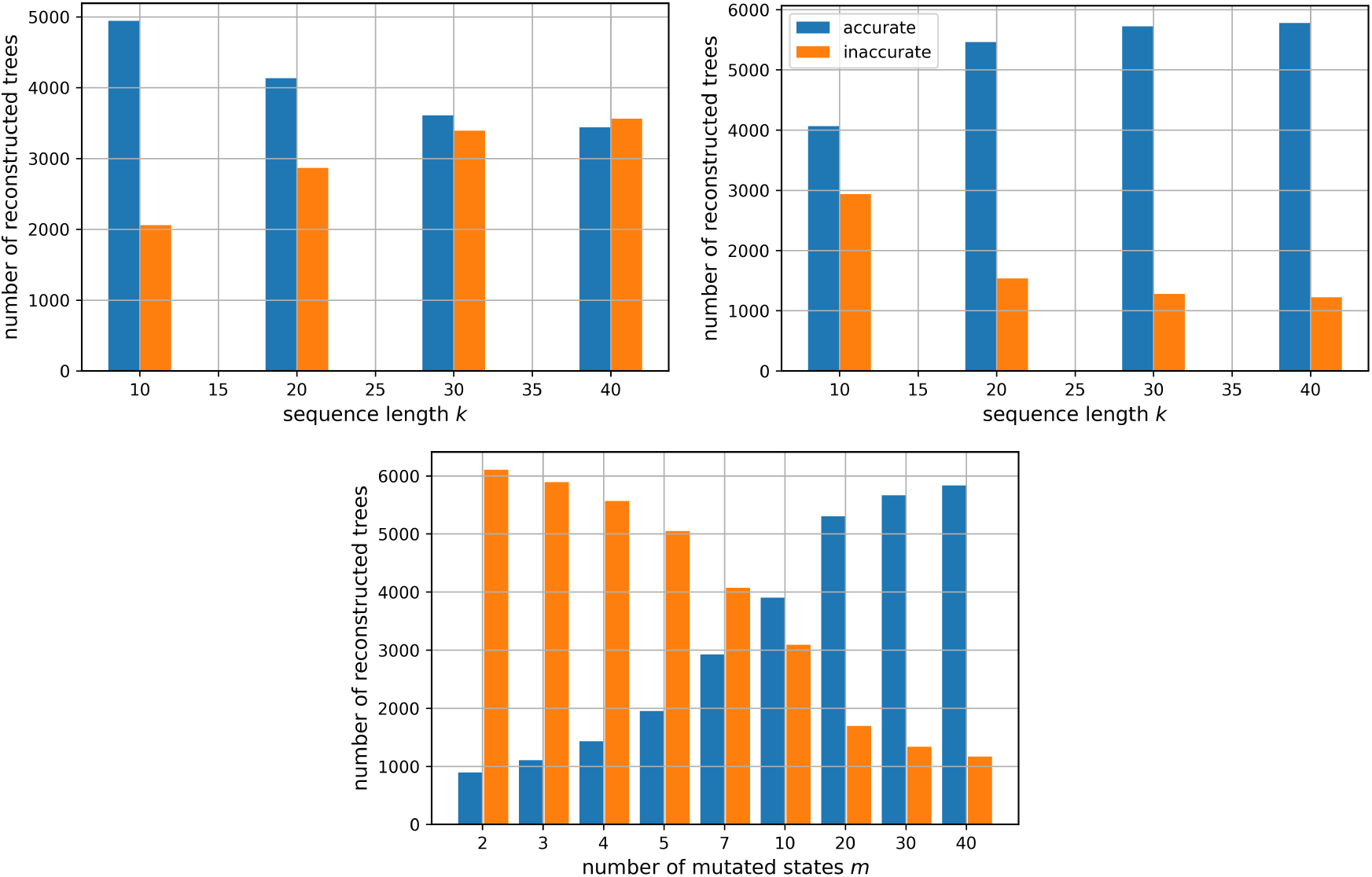
Histograms of accurate and inaccurate reconstructed trees, according to the hypothesis testing (3), as a function of sequence length *k* and number of mutated states *m*. Upper left panel: *d* = *d*_P_ and *ϵ* = 0.01. Upper right panel: *d* = *d*_RF_ and *ϵ* = 1*/*7. Lower panel: *d* = *d*_RF_ and *ϵ* = 1*/*7.

For the composite samples (those combining cells from multiple lesions of an animal, labeled All or Fam), the indel and saturation statistics are not given in the deposit, so we applied only the cell-count criterion (*n* > 100). This stage removed 11 samples, leaving 74. Second, we reconstructed each remaining sample with all six reconstruction algorithms available in Cassiopeia package and removed any sample for which an algorithm failed to produce a tree (most often Neighbor Joining, which errors on some inputs), so that every analyzed sample has a complete set of six reconstructions. This removed a further 11 samples (6 individual tumors and 5 composites), leaving the **63 samples** analyzed in this work. The complete per-sample summary, with all recording parameters, is provided as a supplementary spreadsheet (`KPTracer_summary.xlsx`).

## M Threshold calibration for the KP data

The use of cPHS score with the KP data requires a simulation study dedicated to calibrate the threshold for the parameters of the data. We generated ground-truth trees with the same birth-death process and CRISPR-Cas9 mutation overlay as in Section 4, spanning leaf counts *n* ∈ { 100, 200, 300, 500, 1000, 2000, 3000 } character-site counts *k* ∈ { 7, 10, 15, 20, 25, 30, 50, 70, 100 }, mutation rates *ρ* ∈ { 0.7, 0.9, 0.99}, and collision rates *q* ∈ { 0.02, 0.1, 0.33} with 100 repetitions per configuration. Trees were reconstructed with Cassiopeia Greedy, Maximum Cut, and SMJ. Each tree was labeled accurate or inaccurate by its normalized Robinson–Foulds distance to the ground truth, under cutoffs *ε* ∈ { 0.10, 0.25, 0.50, 0.67 }; for each configuration we found the optimal cPHS threshold (maximizing balanced accuracy) and evaluated the fixed thresholds *t* ∈ {0.05, 0.01, 10^−3^, 10^−4^, 10^−5^, 10^−6^ }. The results of this analysis and the derived cutoffs (*t*), stratified by *n, k, ρ*, and *q*, are presented in Tables S1, S2, S3 and S4. In the range of *n* values that we analyzed we find that generally, for *k* ≤ 15 or *q* > 0.35 the BA values are low, marking these regimes as a-priori difficult to infer accurate trees. Nevertheless, we select a threshold value for these cases (taking the highest scoring one) to enable our analysis of random instances in those regimes. For *k* > 15 the threshold tightens as *ρ* increases, reflecting that higher mutation rates push cPHS values downward.

**Figure S7:**
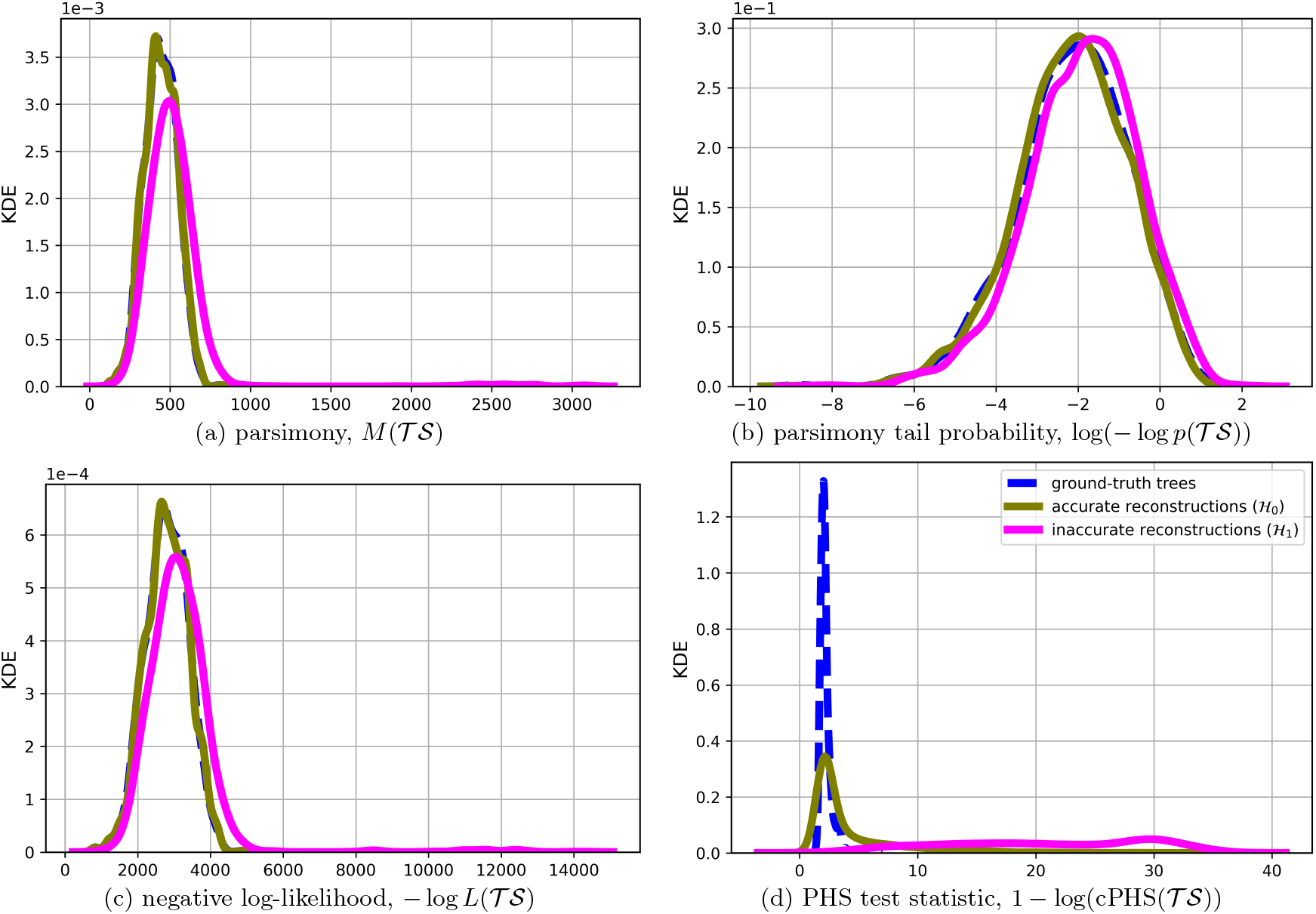
Kernel density estimates for the four accuracy measures, with the same setting as in Fig. 1, but with the parsimony as a distance function (*d* = *d*_P_) and *ϵ* = 0.03.

The calibration results above are summarized into the operating thresholds used throughout the KP analysis (Table S5). The threshold *t* depends on both the number of recording sites *k* and the mutation probability *ρ*. For a given tumor we read off its respective threshold *t* from its measured *k* and *ρ*; since the KPTracer tumors have *ρ* = 0.71–0.999, the analysis uses the two higher-*ρ* columns.

**Figure S8:**
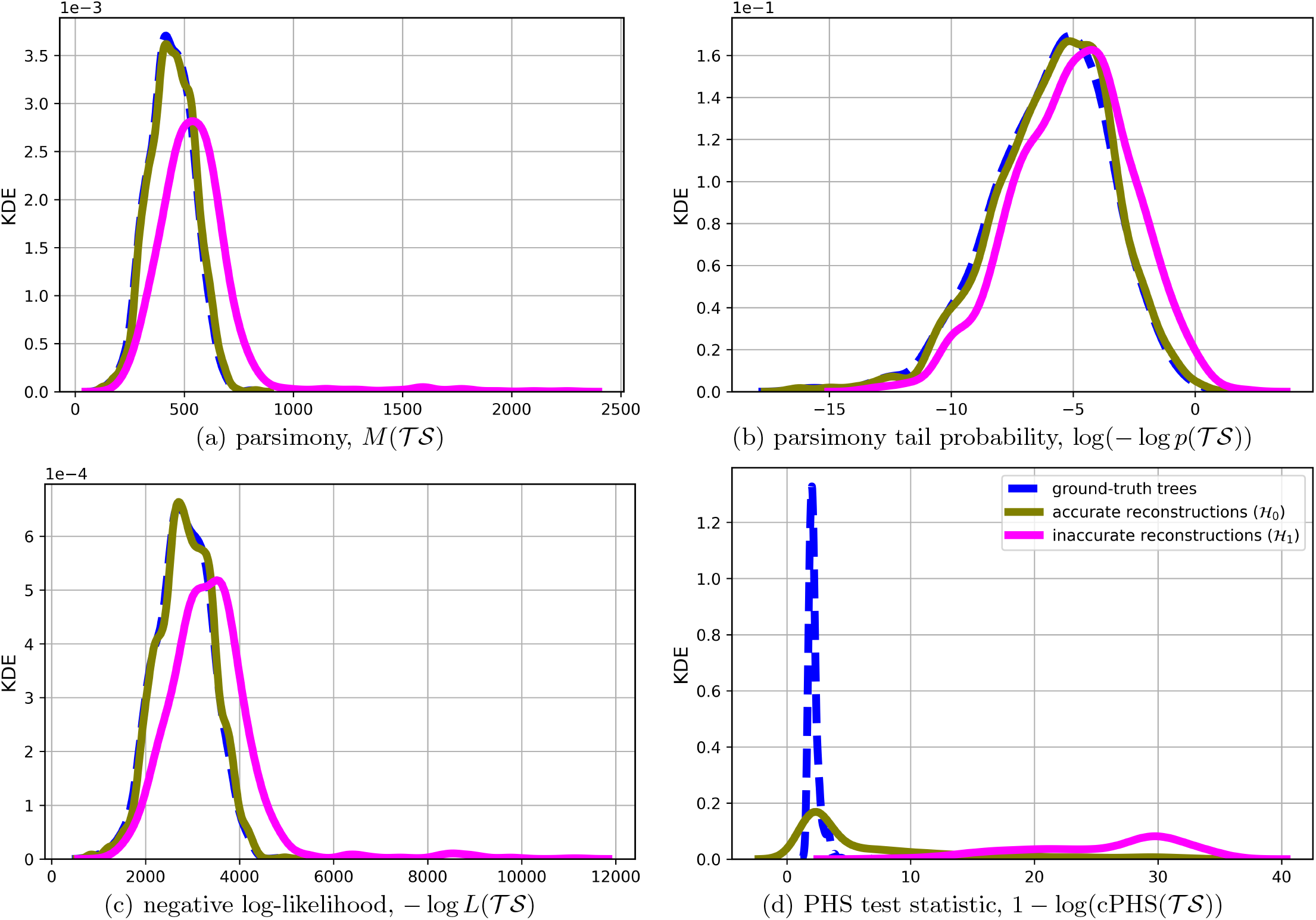
Kernel density estimates for the four accuracy measures, with the same setting as in Fig. 1, but with the parsimony as a distance function (*d* = *d*_P_) and *ϵ* = 0.1.

**Figure S9:**
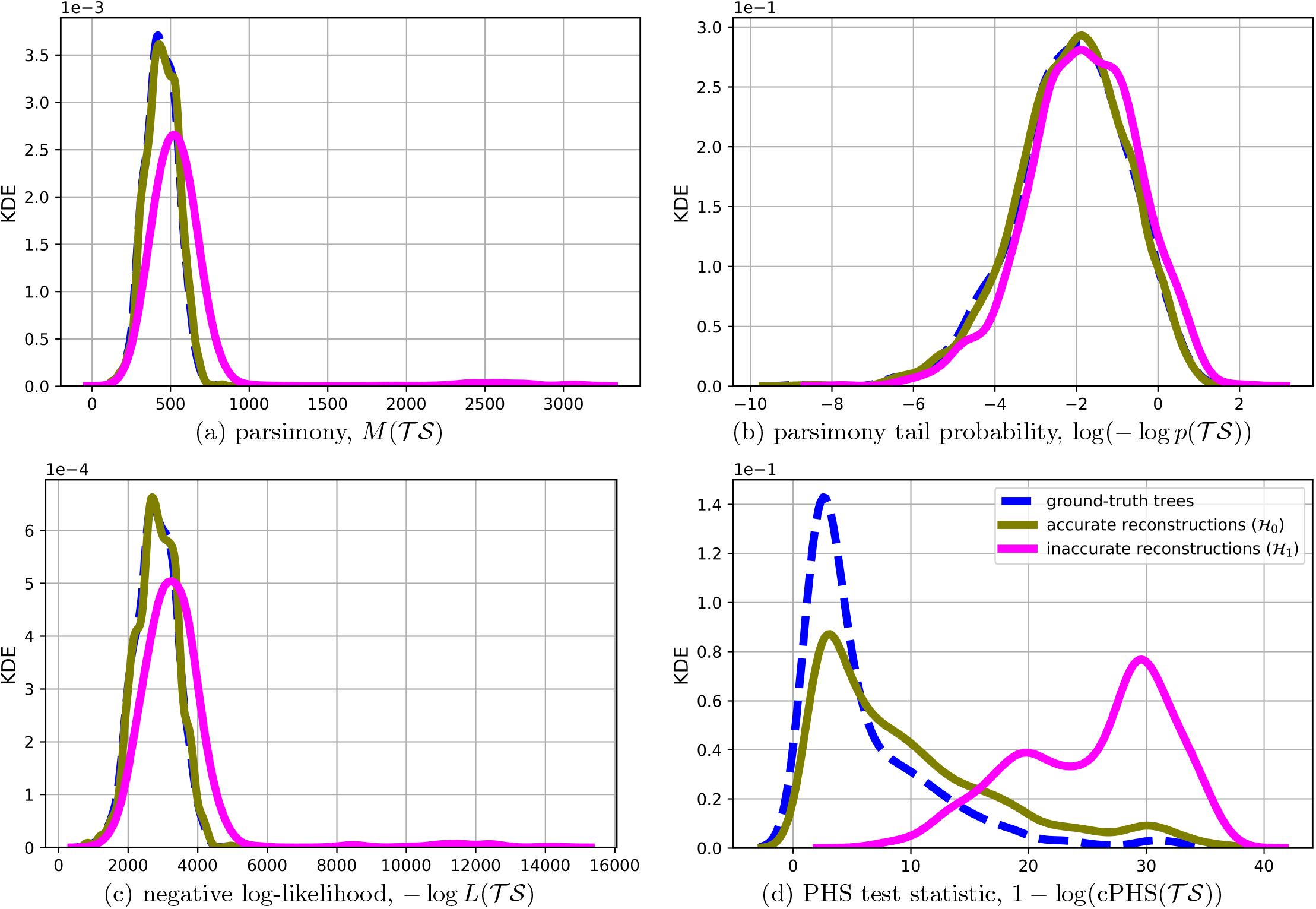
Kernel density estimates for the four accuracy measures, with the same setting as in Fig. 1, but with the likelihood as a distance function (*d* = *d*_L_) and *ϵ* = 0.05.

**Figure S10:**
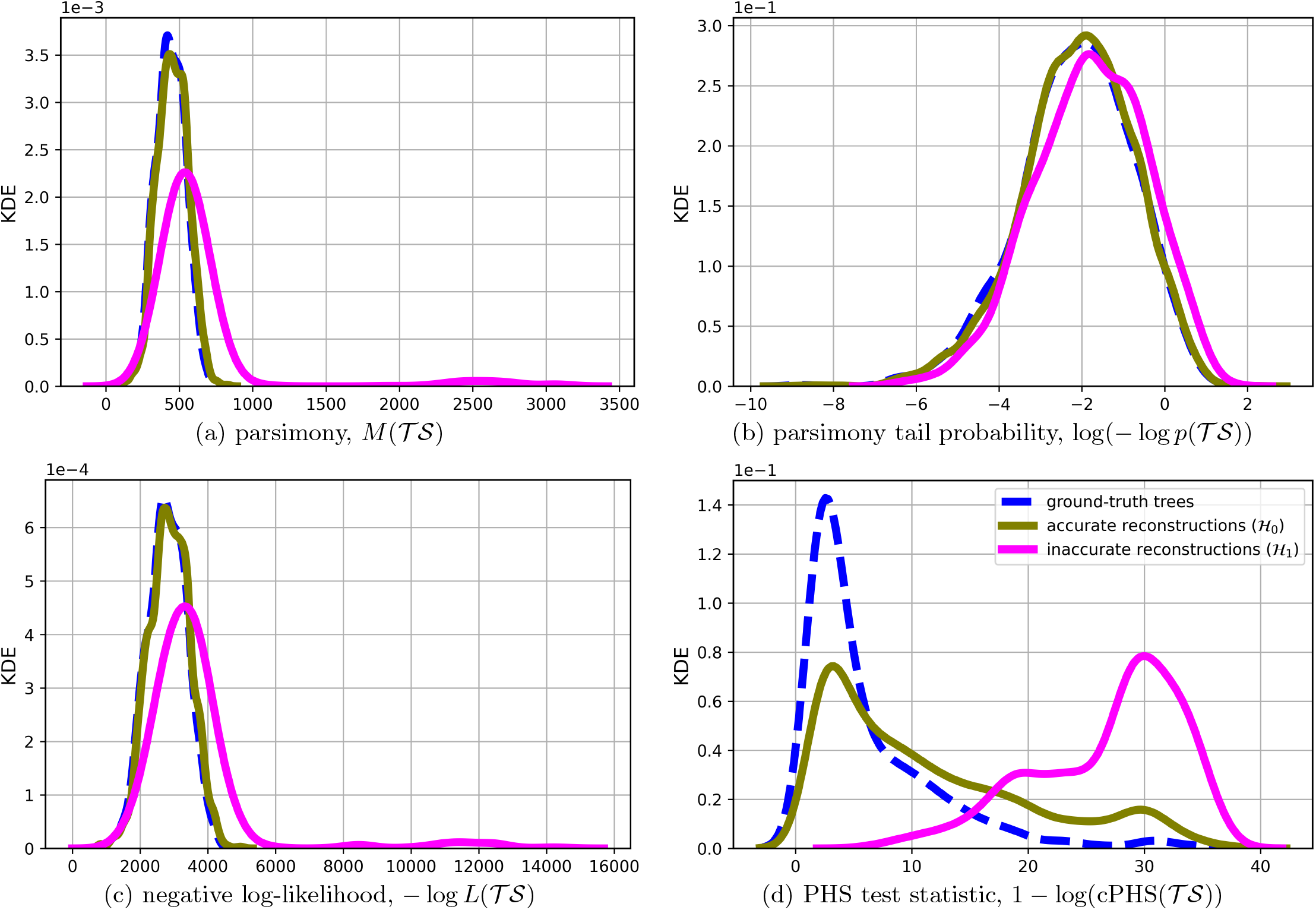
Kernel density estimates for the four accuracy measures, with the same setting as in Fig. 1, but with the triplets as a distance function (*d* = *d*_tri_) and *ϵ* = 1*/*5.

**Figure S11:**
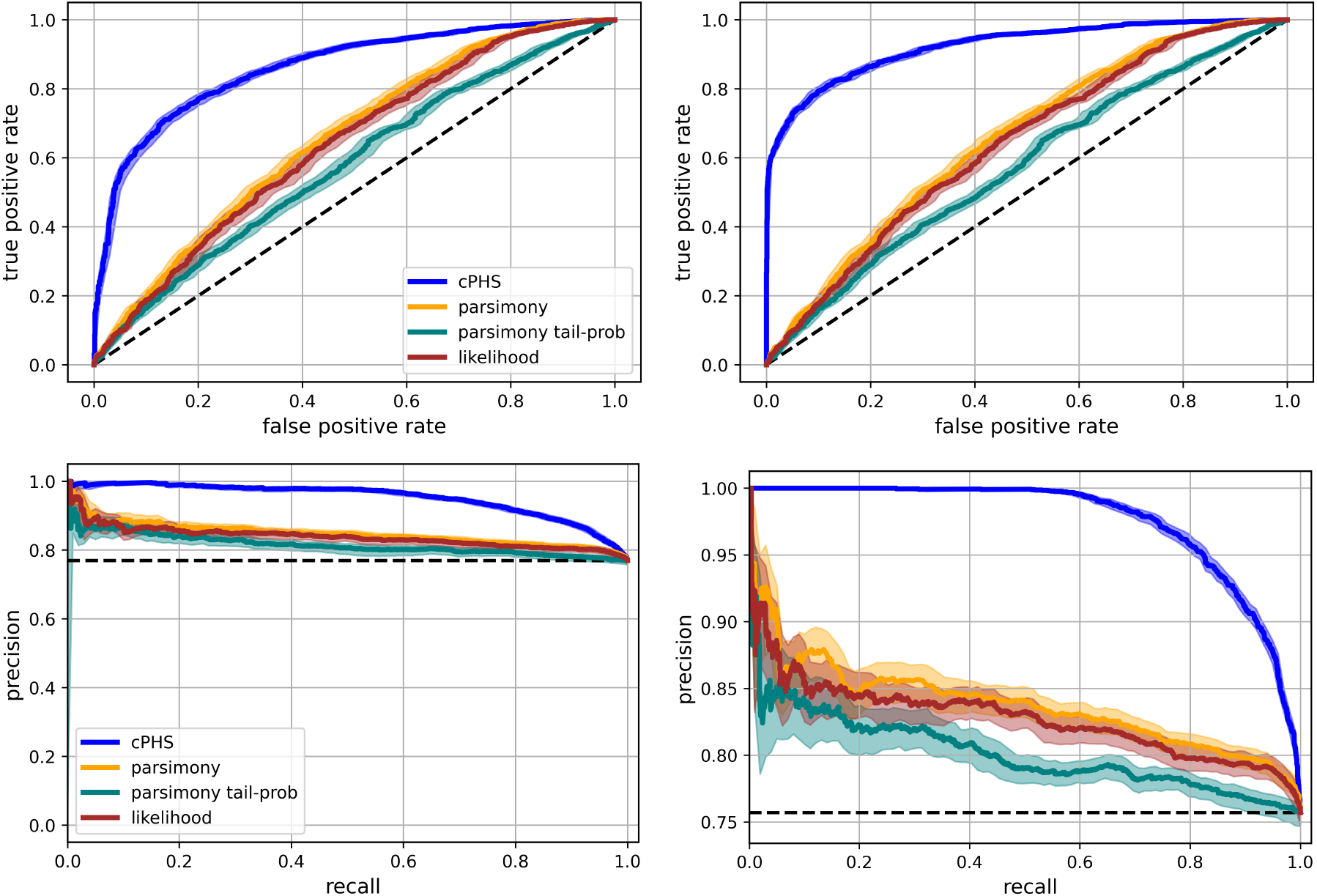
ROC (top) and Precision-Recall (bottom) curves with the same setting as in Fig. 2 except for the distance function. Left: triplets distance (*d* = *d*_tri_) with *ϵ* = 0.1 (left panel). Right: likelihood distance (*d* = *d*_L_) with *ϵ* = 0.03.

**Figure S12:**
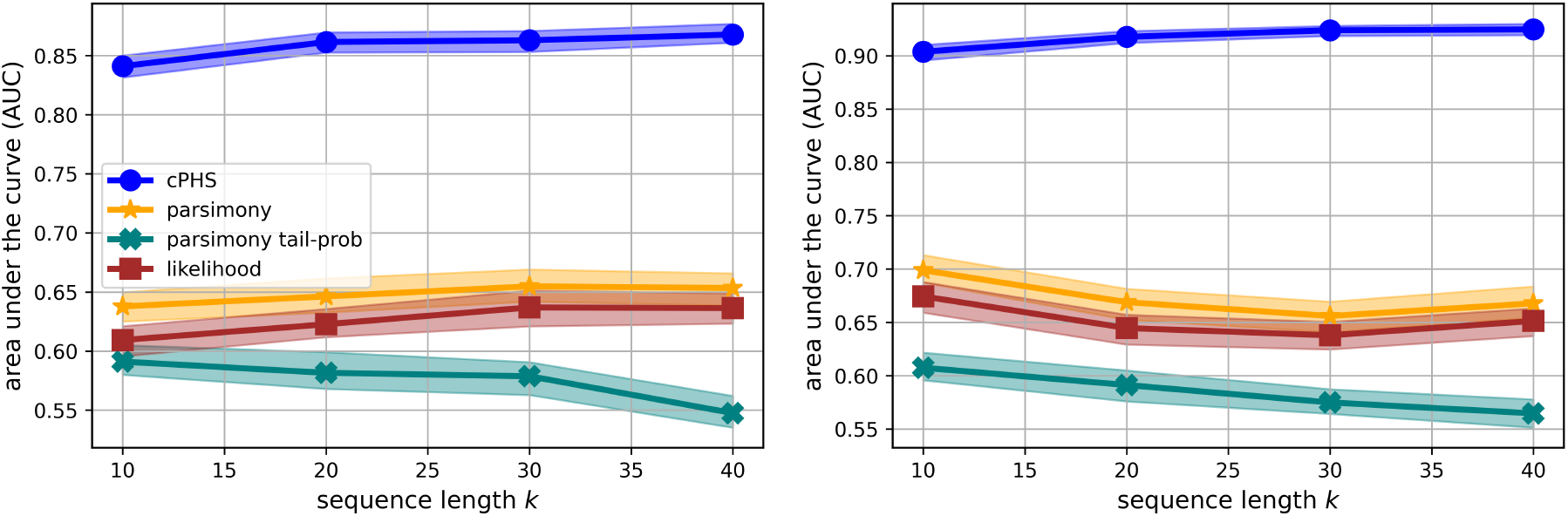
AUC values with the same setting as in Fig. 3(left), except for the distance function. Left: triplets distance (*d* = *d*_tri_) with *ϵ* = 1*/*5. Right: likelihood distance (*d* = *d*_L_) with *ϵ* = 0.01.

**Figure S13:**
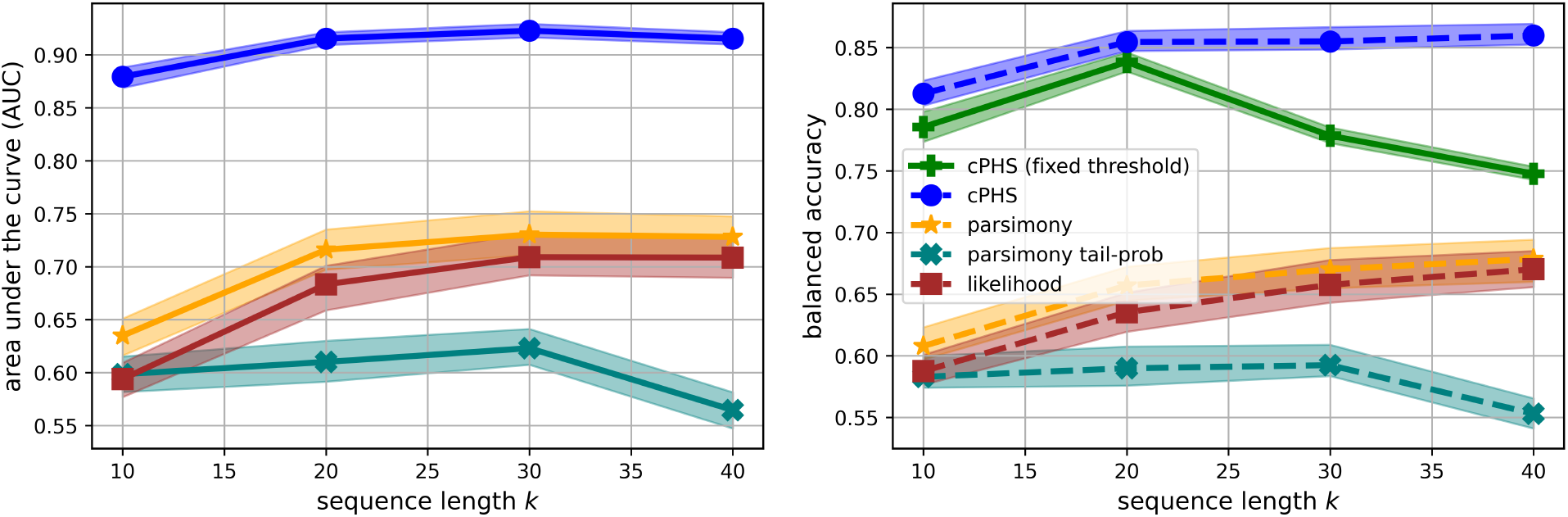
AUC and Balanced Accuracy values with the same setting as in the left panels of Figures. 3 and 4, except for the cutoff value which is *ϵ* = 1*/*4.

**Figure S14:**
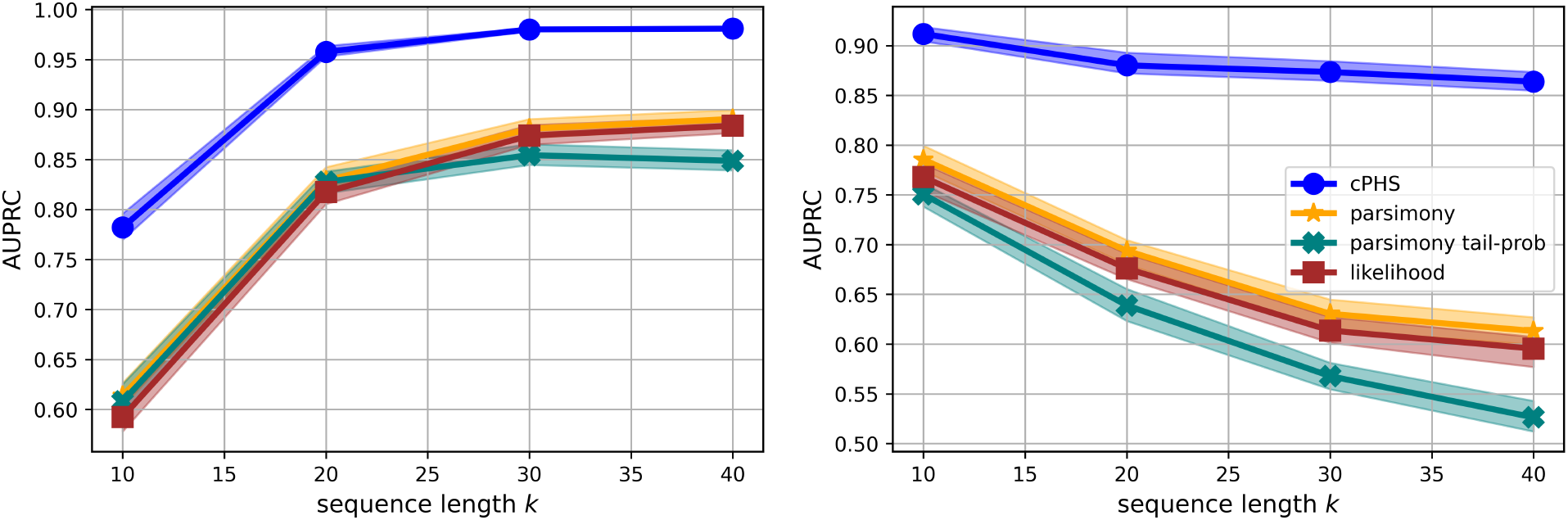
AUPRC (Area Under the Precision-Recall Curve) with the same setting as in the bottom left and right panels of Figure. 2, respectively. That is, left panel shows results for RF distance with *ϵ* = 1*/*7, and right panels shows for parsimony distance with *ϵ* = 0.01.

**Figure S15:**
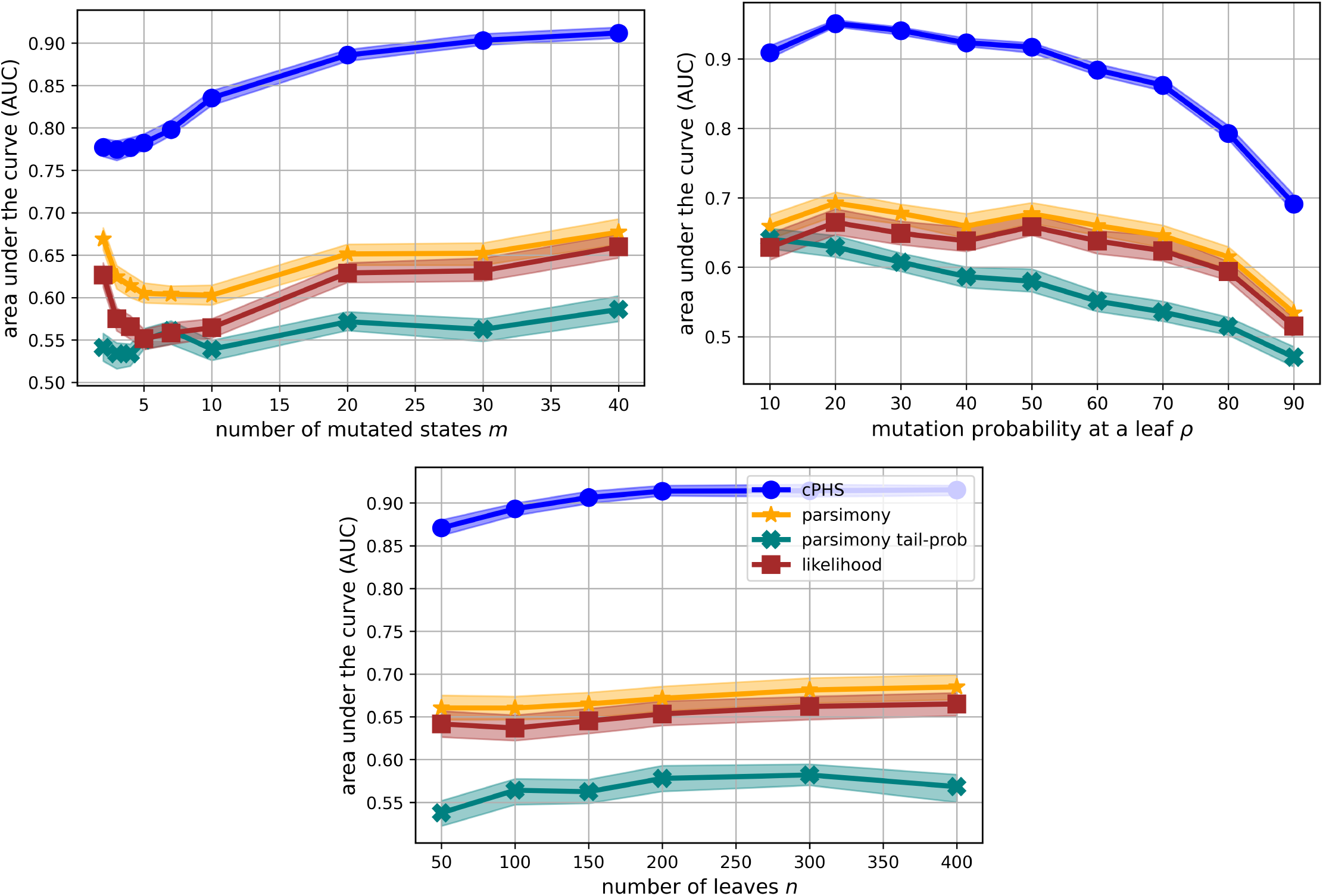
AUC values with the same setting as in Fig. 3, as a function of *m* (top left), *ρ* (top right) and *n* (bottom).

**Figure S16:**
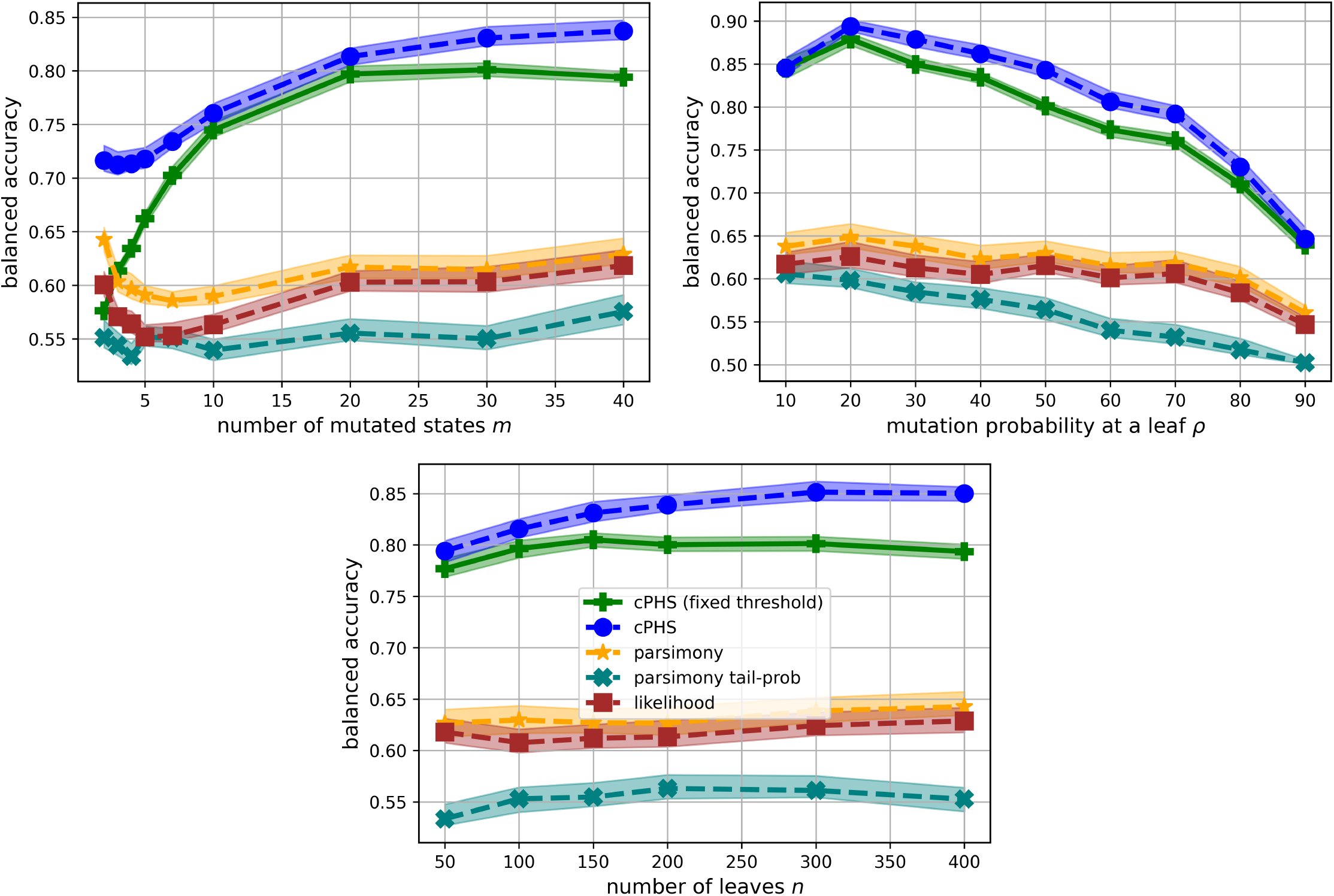
Balanced accuracy of the test statistics with the same setting as in Fig. 4, as a function of *m* (top left), *ρ* (top right) and *n* (bottom).

**Table S1:**
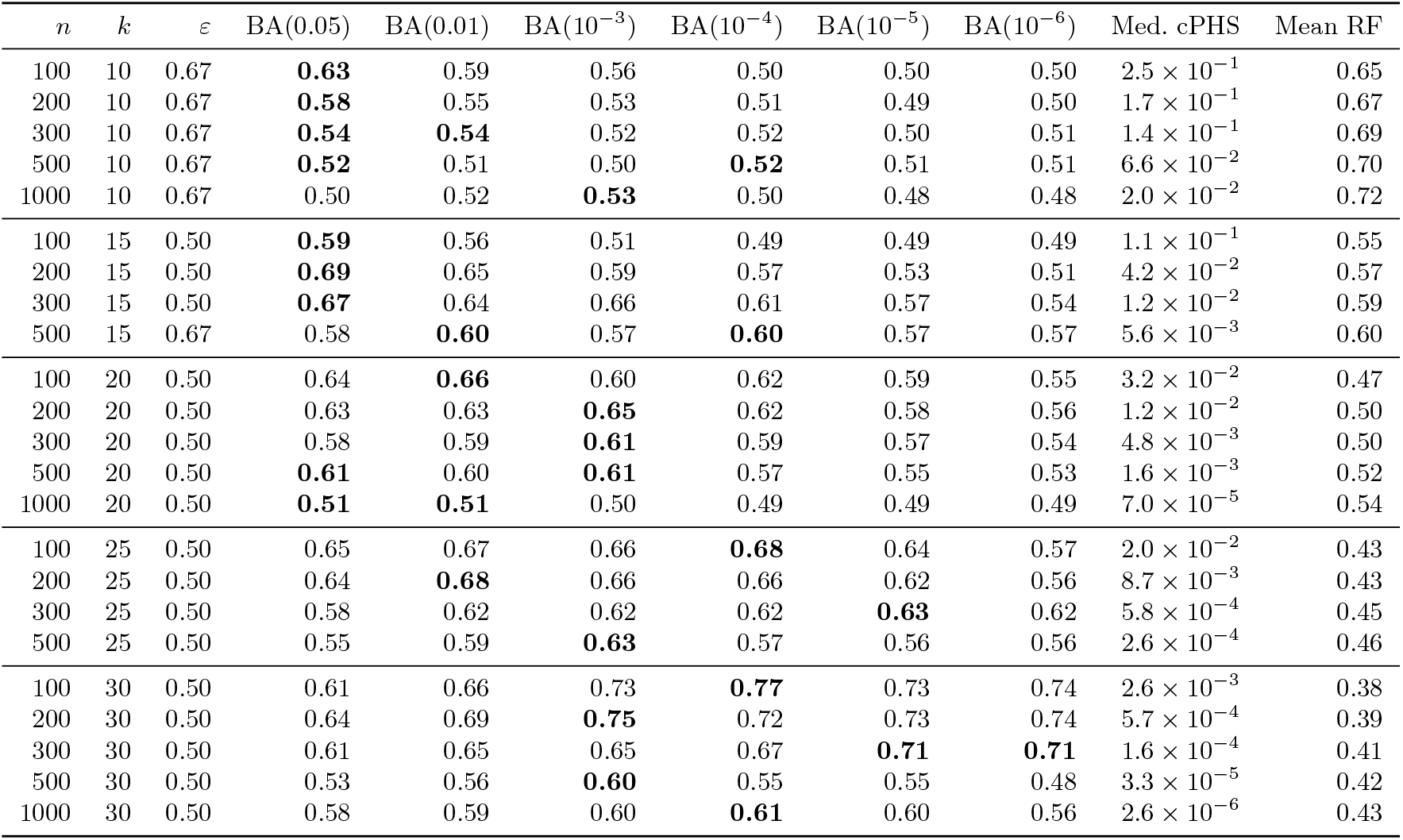
Threshold calibration simulation results at *ρ* = 0.5. For each (*n, k*) configuration, the table shows the balanced accuracy (BA) achieved by six fixed thresholds (best per row in **bold**, where a tree is accepted if cPHS ≥ *t*), and the median cPHS and mean normalized RF distance across all reconstructed trees in that configuration. The *ε* column is the accuracy cutoff used to label a tree accurate (RF ratio ≤ *ε*) or inaccurate; it is fixed by *k*-regime rather than tuned for performance: *ε* = 0.50 (the standard cutoff) for *k* ≥ 15, relaxed to *ε* = 0.67 for *k* = 10, where reconstructions are too poor for *ε* = 0.50 to yield enough accurate trees. The results show that for *k* < 15 no fixed threshold gives reliable balanced accuracy, even under the relaxed cutoff *ε* = 0.67: BA stays near chance (0.5) across all thresholds, confirming that the test lacks power in this regime. Reliable separation emerges only for *k* > 15.

**Table S2:** Effect of mutation probability *ρ* on threshold performance (*n* = 100, *ε* = 0.50). As *ρ* increases, absolute cPHS values drop by orders of magnitude (median cPHS column), shifting the optimal threshold downward. Balanced accuracy (BA) is shown for five fixed thresholds (best per row in **bold**, tree accepted if cPHS ≥ *t*). The adopted thresholds are marked with •: *t* = 10^−3^ for *k* = 16–24 and *t* = 10^−4^ for *k* ≥ 25. The *k* = 15 block is shown for reference only; it is flagged in the analysis, and consistent with Table S1 no fixed threshold separates reliably there.

| $k$ | $\rho$ | BA(0.05) | BA(0.01) | BA($10^{-3}$) | BA($10^{-4}$) | BA($10^{-5}$) | Med. cPHS |
| --- | --- | --- | --- | --- | --- | --- | --- |
| 15 | 0.5 | **0.59** | 0.56 | 0.51 | 0.49 | 0.49 | $1.1 \times 10^{-1}$ |
| 15 | 0.7 | **0.64** | 0.62 | 0.63 | 0.61 | 0.57 | $3.2 \times 10^{-2}$ |
| 15 | 0.9 | 0.67 | 0.69 | **0.73** | 0.72 | 0.70 | $3.7 \times 10^{-3}$ |
| 15 | 0.99 | 0.62 | 0.64 | 0.65 | 0.66 | **0.69** | $4.2 \times 10^{-5}$ |
| 20 | 0.5 | 0.64 | **0.66** | 0.60$\bullet$ | 0.62 | 0.59 | $3.2 \times 10^{-2}$ |
| 20 | 0.7 | 0.69 | 0.71 | 0.73$\bullet$ | **0.77** | 0.69 | $4.6 \times 10^{-3}$ |
| 20 | 0.9 | 0.62 | 0.63 | 0.61$\bullet$ | 0.66 | **0.68** | $3.5 \times 10^{-5}$ |
| 20 | 0.99 | 0.61 | 0.66 | 0.70$\bullet$ | 0.70 | **0.72** | $2.3 \times 10^{-5}$ |
| 25 | 0.5 | 0.65 | 0.67 | 0.66 | **0.68**$\bullet$ | 0.64 | $2.0 \times 10^{-2}$ |
| 25 | 0.7 | 0.66 | 0.71 | 0.77 | 0.82$\bullet$ | **0.86** | $1.4 \times 10^{-3}$ |
| 25 | 0.9 | 0.61 | 0.66 | 0.69 | 0.75$\bullet$ | **0.78** | $1.8 \times 10^{-5}$ |
| 25 | 0.99 | 0.49 | 0.52 | 0.57 | 0.60$\bullet$ | **0.62** | $6.6 \times 10^{-10}$ |
| 30 | 0.5 | 0.61 | 0.66 | 0.73 | **0.77**$\bullet$ | 0.73 | $2.6 \times 10^{-3}$ |
| 30 | 0.7 | 0.61 | 0.65 | 0.72 | 0.78$\bullet$ | **0.83** | $4.7 \times 10^{-4}$ |
| 30 | 0.9 | 0.57 | 0.59 | 0.63 | **0.67**$\bullet$ | 0.56 | $2.3 \times 10^{-8}$ |
| 30 | 0.99 | 0.50 | 0.52 | 0.53 | 0.54$\bullet$ | **0.58** | $4.2 \times 10^{-10}$ |

## N Comparison to Likelihood and Parsimony

The consistency between cPHS scores and that of parsimony and likelihood can be summarized as a best-method concordance. Table S6 reports, for each pair of metrics, the fraction of the 42 reliable tumors in which the two metrics select the same best method. cPHS and likelihood agree most often (36%), cPHS and parsimony slightly less (31%), and parsimony and likelihood least (24%); all three coincide in only 10% of tumors (4 of 42, all on Cassiopeia Greedy). The cPHS–likelihood disagreement is systematic rather than noise: when cPHS prefers a method other than the one likelihood favors, likelihood selects SMJ in two-thirds of those cases (18 of 27), reflecting that SMJ optimizes an objective close to likelihood during construction. The metrics thus capture overlapping but distinct notions of tree quality, consistent with the moderate rank correlations reported above.

## O Factors influencing cPHS performance in KP data

Considering the objective of rejecting random trees - the collision probability *q* is the strongest predictor of cPHS separation (Spearman *ρ* = − 0.54, *p* = 1.4 × 10^−6^), stronger than the effective homoplasy rate *λ* × *q* (*ρ* = − 0.50) or *λ* alone (*ρ* = − 0.31); see Figure S19. Separation is strong for *q* < 0.35 (mean 7.6 orders of magnitude), degrades for 0.35 ≤ *q* < 0.75 (mean 3.2), and collapses for *q* ≥ 0.75. Practically, *q* can be estimated directly from character-state frequencies, so cPHS applicability can be assessed directly.

Focusing on the 42 tumors with *q* < 0.35 and *k* > 15, 14 had no inferred tree passing the cPHS threshold. We examined tumor-level recording parameters and data-quality indicators (number of cells *n*, sites *k*, collision probability *q*, mutation probability *ρ*, percent missing data, unique alleles per site, percent unique cell profiles, percent unsaturated sites, and the derived quantities *k/n, k/* log *n, k* · (1 − *q*), and *k* · log(1*/q*)) and correlated each with the best log_10_(cPHS) per tumor. The correlations are summarized in Table S8.

The dominant correlate is tumor size (*r* = −0.77 between *n* and best log_10_(cPHS)): passing tumors have median 300 cells vs. 1,145 for non-passing, and all 4 tumors with *n* > 2,000 and 8 of 10 with *n* > 1,000 fall in the no-pass category. The rationale is the need for increased number of recording sites for larger number of leaves, which has been shown to be quadratic in nature. The ratio *k/n* is thus the single most discriminative predictor (*r* = +0.70): all 14 non-passing tumors have *k/n* < 0.05, and above this value every tumor has at least one method passing. The positive correlation of *q* with passing (*r* = +0.70) reflects the opposite effect, since higher *q* inflates cPHS toward the threshold by raising chance homoplasy, and is itself a size confound, since larger tumors have lower *q*.

**Table S3:** Effect of mutation probability *ρ* on threshold performance at *n* = 500, *ε* = 0.50 (compare Table S2 for *n* = 100). The same threshold ranking holds, but balanced accuracy is consistently lower, reflecting the increased difficulty of classifying larger trees. Balanced accuracy (BA) is shown for five fixed thresholds (best per row in **bold**, tree accepted if cPHS ≥ *t*). The adopted thresholds are marked with •: *t* = 10^−3^ for *k* = 16–24 and *t* = 10^−4^ for *k* ≥ 25. The *k* = 15 block is shown for reference only; it is flagged in the analysis.

| $k$ | $\rho$ | BA(0.05) | BA(0.01) | BA($10^{-3}$) | BA($10^{-4}$) | BA($10^{-5}$) | Med. cPHS |
| --- | --- | --- | --- | --- | --- | --- | --- |
| 15 | 0.5 | 0.59 | **0.77** | 0.71 | 0.63 | 0.57 | $5.6 \times 10^{-3}$ |
| 15 | 0.7 | **0.60** | **0.60** | 0.57 | 0.55 | 0.55 | $1.2 \times 10^{-3}$ |
| 15 | 0.9 | 0.59 | **0.64** | 0.61 | 0.61 | **0.64** | $2.3 \times 10^{-4}$ |
| 15 | 0.99 | **0.92** | 0.90 | 0.85 | 0.82 | 0.79 | $1.9 \times 10^{-7}$ |
| 20 | 0.5 | **0.61** | 0.60 | **0.61**$\bullet$ | 0.57 | 0.55 | $1.6 \times 10^{-3}$ |
| 20 | 0.7 | 0.60 | 0.63 | 0.62$\bullet$ | **0.64** | 0.61 | $2.5 \times 10^{-4}$ |
| 20 | 0.9 | 0.57 | 0.59 | 0.58$\bullet$ | 0.60 | **0.61** | $1.5 \times 10^{-5}$ |
| 20 | 0.99 | 0.49 | 0.54 | 0.54$\bullet$ | 0.56 | **0.60** | $7.6 \times 10^{-8}$ |
| 25 | 0.5 | 0.55 | 0.59 | **0.63** | 0.57$\bullet$ | 0.56 | $2.6 \times 10^{-4}$ |
| 25 | 0.7 | 0.59 | 0.63 | 0.66 | 0.71$\bullet$ | **0.75** | $1.6 \times 10^{-6}$ |
| 25 | 0.9 | 0.55 | 0.57 | 0.61 | 0.65$\bullet$ | **0.67** | $5.0 \times 10^{-8}$ |
| 25 | 0.99 | 0.51 | 0.53 | 0.56 | 0.59$\bullet$ | **0.60** | $1.2 \times 10^{-8}$ |
| 30 | 0.5 | 0.53 | 0.56 | **0.60** | 0.55$\bullet$ | 0.55 | $3.3 \times 10^{-5}$ |
| 30 | 0.7 | 0.58 | 0.61 | 0.63 | 0.69$\bullet$ | **0.70** | $1.3 \times 10^{-6}$ |
| 30 | 0.9 | 0.55 | 0.57 | 0.60 | 0.62$\bullet$ | **0.65** | $5.6 \times 10^{-10}$ |
| 30 | 0.99 | 0.52 | 0.54 | 0.56 | 0.56$\bullet$ | **0.58** | $3.8 \times 10^{-11}$ |

### Simulation-based validation

Simulating trees across the KP (*n, q*) regime (*n* = 100–2,000, *q* ≈ 0.02–0.33, *k* = 20, 30, *ρ* = 0.9) reproduces the empirical pattern: pass rates fall with *n* and rise with *q*, and the KP tumors that fail cluster in the high-*n*, low-*q* region where simulations also predict low pass rates. This supports the interpretation that cPHS failures in the reliable set are driven by insufficient *k* relative to *n*.

## P Degradation experiments for KP data: NNI perturbation and data degragation

We assessed cPHS sensitivity to topological degradation by using controlled NNI perturbation of the reconstructed trees.

### P.1 NNI perturbation

To evaluate how cPHS responds to controlled degradation of tree topology, we performed Nearest Neighbor Interchange (NNI) perturbation experiments. Starting from the reconstructed tree for each tumor, we applied random NNI moves that swap subtrees across internal edges, progressively distorting the topology while holding the character data fixed. After each perturbation level we measured cPHS, parsimony, likelihood, as well as Robinson–Foulds (RF) distance and triplets-correct distance to the original tree.

#### Edge-normalized perturbation

A fair comparison of perturbation effects between reconstruction methods requires accounting for differences in tree topology. Cassiopeia Greedy produces shallow trees with many polytomies (few internal edges, large fan-out), whereas Shared Mutation Joining (SMJ) produces fully binary trees (many internal edges, all bifurcating). Applying the same number of NNI moves to both tree types produces unequal effective perturbation: each move on a Cassiopeia tree swaps larger subtrees, and so induces greater topological change, than the same move on a binary SMJ tree. To make the two comparable, we normalized the number of NNI moves by the number of internal edges in each tree. Perturbation levels were set at 0, 5, 10, 25, 50, 100, 150, 200% of internal edges. At 100%, both methods reach comparable RF distances to the original tree (Cassiopeia 0.59 *±* 0.04, SMJ 0.57 *±* 0.03), confirming that the normalization produces equivalent effective perturbation.

**Table S4:** Effect of collision probability *q* on threshold performance (*n* = 100, *ε* = 0.50). Higher *q* shifts cPHS values upward (median cPHS column), making strict thresholds less effective. Balanced accuracy (BA) is shown for five fixed thresholds (best per row in **bold**, tree accepted if cPHS ≥ *t*). At *q* = 0.33 (the upper boundary of the low-*q* regime), the adopted threshold *t* = 10^−4^ (•) achieves only moderate balanced accuracy, while *t* = 0.05 becomes the best-performing threshold. This independently validates the partitioning of tumors at *q* = 0.35: the calibrated thresholds are effective for low-*q* tumors but would not be appropriate for high-*q* tumors.

| $k$ | $q$ | $m$ | $\rho$ | BA(0.05) | BA(0.01) | BA($10^{-3}$) | BA($10^{-4}$) | BA($10^{-5}$) | Med. cPHS |
| --- | --- | --- | --- | --- | --- | --- | --- | --- | --- |
| 25 | 0.02 | 50 | 0.5 | 0.65 | 0.67 | 0.66 | **0.68•** | 0.64 | $2.0 \times 10^{-2}$ |
| 25 | 0.02 | 50 | 0.9 | 0.61 | 0.66 | 0.69 | 0.75• | **0.78** | $1.8 \times 10^{-5}$ |
| 25 | 0.10 | 10 | 0.5 | **0.65** | **0.65** | 0.63 | 0.55• | 0.51 | $7.2 \times 10^{-2}$ |
| 25 | 0.10 | 10 | 0.9 | 0.69 | 0.68 | **0.72** | 0.69• | 0.66 | $1.3 \times 10^{-3}$ |
| 25 | 0.33 | 3 | 0.5 | **0.64** | 0.59 | 0.55 | 0.51• | 0.50 | $3.1 \times 10^{-1}$ |
| 25 | 0.33 | 3 | 0.9 | **0.69** | 0.63 | 0.56 | 0.54• | 0.52 | $2.3 \times 10^{-1}$ |
| 30 | 0.02 | 50 | 0.5 | 0.61 | 0.66 | 0.73 | **0.77•** | 0.73 | $2.6 \times 10^{-3}$ |
| 30 | 0.02 | 50 | 0.9 | 0.57 | 0.59 | 0.63 | **0.67•** | 0.56 | $2.3 \times 10^{-8}$ |
| 30 | 0.10 | 10 | 0.5 | **0.74** | **0.74** | 0.70 | 0.65• | 0.59 | $7.2 \times 10^{-2}$ |
| 30 | 0.10 | 10 | 0.9 | 0.67 | 0.67 | **0.71** | 0.68• | 0.67 | $1.3 \times 10^{-3}$ |
| 30 | 0.33 | 3 | 0.5 | **0.63** | 0.63 | 0.57 | 0.54• | 0.52 | $1.9 \times 10^{-1}$ |
| 30 | 0.33 | 3 | 0.9 | **0.66** | 0.63 | 0.61 | 0.56• | 0.53 | $1.4 \times 10^{-1}$ |

**Table S5:** Adopted cPHS thresholds *t* as a function of the number of recording sites *k* and the mutation probability *ρ*. Tumors with *k* ≤ 15 are flagged for insufficient power. Since the KPTracer tumors have *ρ* = 0.71–0.999, the analysis uses the middle and right columns.

| | $\rho = 0.5$–$0.6$ | $\rho = 0.7$–$0.9$ | $\rho = 0.99$ |
| --- | --- | --- | --- |
| $k \leq 15$ | flagged (insufficient power) | | |
| $k = 16$–$24$ | $t = 0.05$ | $t = 10^{-3}$ | $t = 10^{-4}$ |
| $k \geq 25$ | $t = 10^{-3}$ | $t = 10^{-4}$ | $t = 10^{-4}$ |

#### Tumors analyzed

NNI results were computed for 34 of the reliable low-*q* tumors, each perturbed with 20 replicates per level. All 34 contribute to both the cPHS pass-rate and the continuous-metric curves. A tumor is counted as passing at a given perturbation level if a majority of its replicate trees pass cPHS.

#### Results

Figures S20 show that the cPHS pass rate declines with perturbation, pooled across tumors and averaged per tumor respectively, confirming that cPHS detects topological degradation. The two standard tree-comparison metrics move consistently with it: RF distance to the original rises monotonically and triplets-correct falls monotonically as perturbation increases, for both methods, with the Cassiopeia Greedy algorithm showing wider tumor-to-tumor spread than SMJ in both. Figure S21 shows all four continuous metrics (parsimony, RF, triplets, likelihood) degrading monotonically and consistently for both methods, validating the edge-normalized scheme. Per-tumor parsimony, likelihood, RF and triplets trajectories for representative tumors are given in Figure S20, where the tight replicate agreement confirms that the degradation is reproducible regardless of the specific random sequence of moves. Table S10 aggregates the effect over the 48 low-*q* tumors: the cPHS pass rate falls from 41.7% (intact) to 16.7% at 50% NNI, with median cPHS, parsimony, and likelihood all degrading monotonically while random trees remain fully separated (0% pass).

The same perturbation analysis also confirms why the low-*k* tumors are flagged. Table S9 shows cPHS for NNI-perturbed Cassiopeia Greedy trees of the six flagged tumors (*k* ≤ 15): even trees with 25–50% of their topology scrambled routinely retain cPHS values that would clear any reasonable threshold, so cPHS cannot distinguish a good tree from a badly perturbed one at these site counts. This is in direct contrast to the *k* ≥ 25 tumors, where 50%-perturbed trees almost never pass.

**Figure S17:**
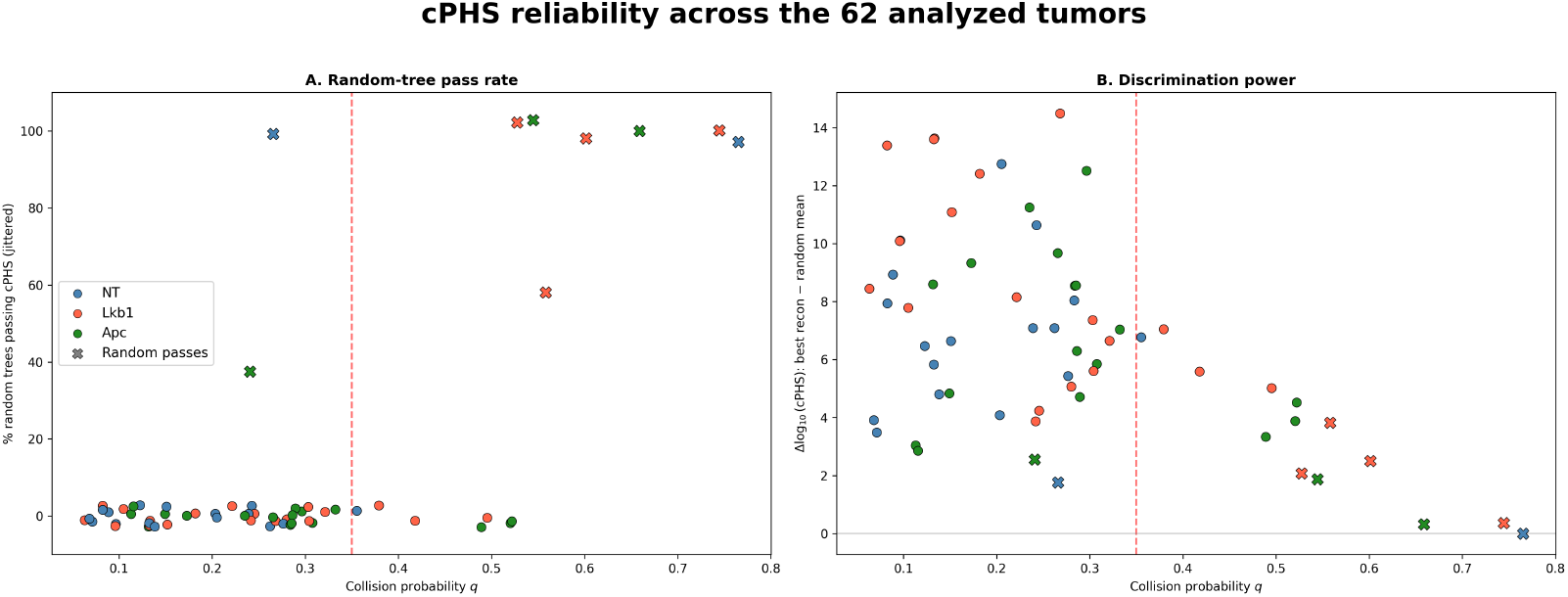
cPHS reliability across the 63 analyzed KPTracer tumors, evaluated at the calibrated thresholds (*t* = 10^−3^ for *k ≤* 24, *t* = 10^−4^ for *k ≥* 25). (A) Fraction of random trees passing cPHS vs. collision probability *q*; markers colored by genotype, X marks tumors with at least one random pass. Random passes occur in 10 tumors, 7 at high *q* (*q* ≥ 0.35, where chance homoplasy lifts cPHS) and 3 at low *q* but few sites (*k* ≤ 15, where the test lacks power). (B) Discrimination power, the log_10_ separation between the best reconstruction’s cPHS and the random-tree mean, vs. *q*; large at low *q*, collapsing as *q* increases. Red dashed line: *q* = 0.35. No random tree passes in the 42 reliable tumors (*k* > 15, *q* < 0.35).

**Table S6:** Best-method concordance across the 42 reliable low-*q* tumors. Each off-diagonal entry is the fraction of tumors in which the row and column metrics select the same best reconstruction method. All three metrics agree on the single best method in only 10% of tumors (4 of 42), reflecting that each rewards a different property of the tree.

|  | cPHS | Parsimony | Likelihood |
| --- | --- | --- | --- |
| cPHS | — | 31% | 36% |
| Parsimony | 31% | — | 24% |
| Likelihood | 36% | 24% | — |

---

**Figure S18:**
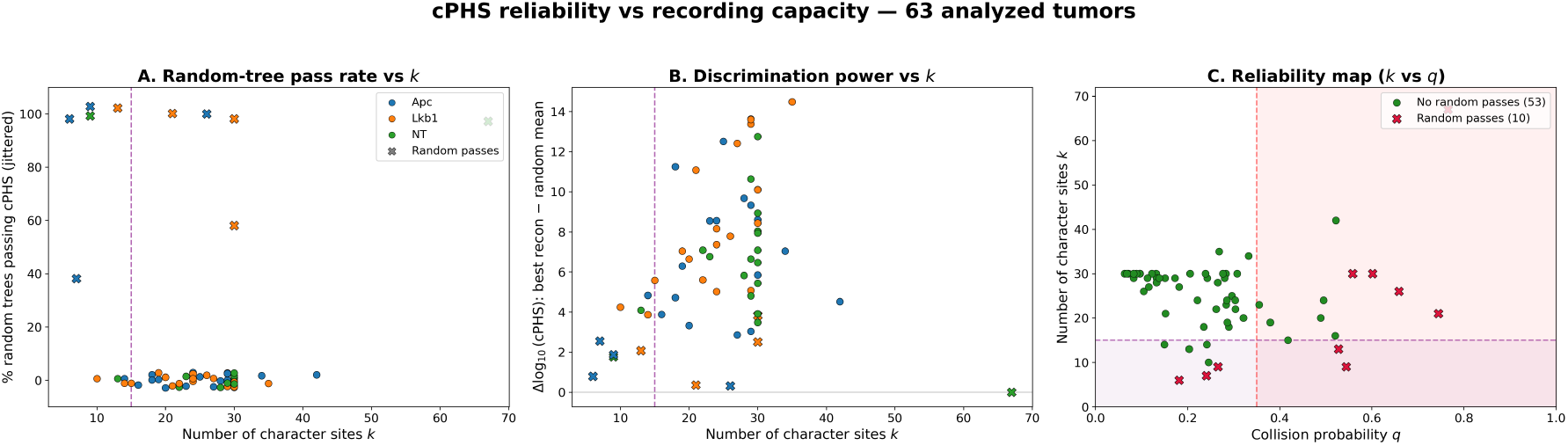
cPHS reliability as a function of recording capacity across the 63 analyzed KPTracer tumors, evaluated at the calibrated thresholds (*t* = 10^−3^ for *k* ≤ 24, *t* = 10^−4^ for *k* ≥ 25). (A) Fraction of random trees passing cPHS vs. number of character sites *k*; markers colored by genotype, X marks tumors with at least one random pass, and the purple dashed line marks *k* = 15. (B) Discrimination power, the log_10_ separation between the best reconstruction’s cPHS and the random-tree mean, vs. *k*. (C) Reliability map of *k* vs. collision probability *q*: tumors with no random passes (green) versus at least one random pass (red X), with the reliable region (*k* > 15, *q* < 0.35) unshaded and the low-*k* (*k* ≤ 15) and high-*q* (*q* ≥ 0.35) zones shaded. Random passes concentrate in the low-*k* and high-*q* zones; the reliable region is free of them.

**Table S7:** Number of tumors (out of the 42 reliable) in which each method achieves the best score under each metric. Cassiopeia Greedy is preferred most often by cPHS and parsimony, whereas SMJ is preferred most often by likelihood, reflecting that the three metrics reward different properties.

| Method | Best cPHS | Best parsimony | Best likelihood |
| --- | --- | --- | --- |
| Cassiopeia Greedy | 18 | 24 | 14 |
| Maximum Cut | 8 | 17 | 3 |
| SMJ | 11 | 1 | 25 |
| Neighbor Joining | 5 | 0 | 0 |
| Spectral Greedy | 0 | 0 | 0 |
| Spectral | 0 | 0 | 0 |

**Figure S19:**
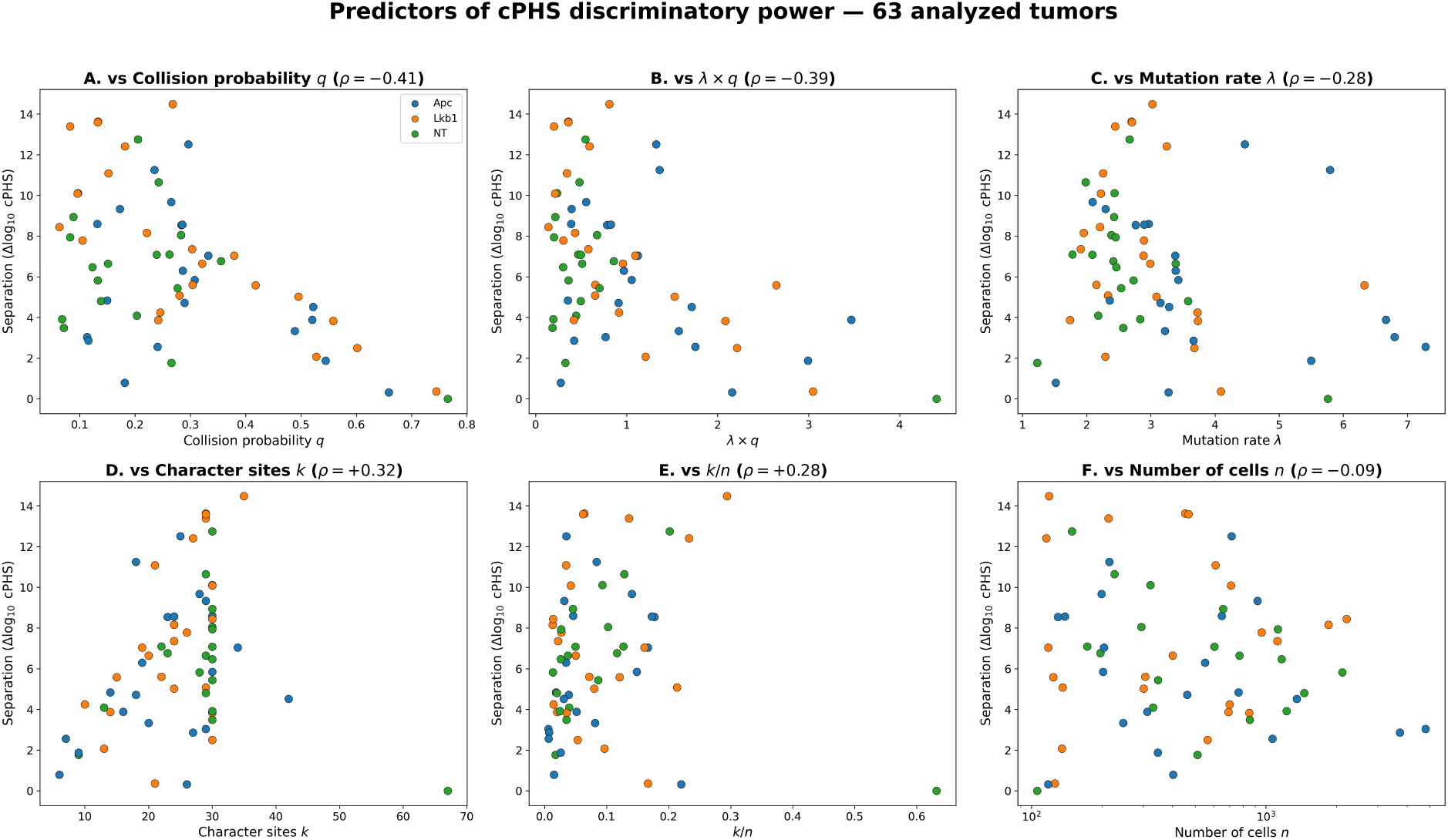
Predictors of cPHS discriminatory power across the 63 analyzed KPTracer tumors. Separation is the log_10_ gap between the best reconstruction’s cPHS and the random-tree mean; points colored by genotype. Collision probability *q* is the strongest single predictor (A, *ρ* = −0.41), followed by *λ × q* (B, *ρ* = −0.39), with weaker dependence on the mutation rate *λ* (C, *ρ* = −0.28), character sites *k* (D, *ρ* = +0.32), the ratio *k/n* (E, *ρ* = +0.28), and number of cells *n* (F, *ρ* = −0.09, n.s.).

**Table S8:** Key quality indicators for the 42 reliable low-*q* tumors, stratified by cPHS outcome. Spearman *r* is the correlation with log_10_(best cPHS) across all 42. ^∗^ *p* < 0.05; ^∗∗^ *p* < 0.01.

| Indicator | Passes (28) | No pass (14) | Spearman $r$ | $p$ |
| --- | --- | --- | --- | --- |
| $n$ (cells) | 300 | 1,145 | -0.77 | $< 10^{-4**}$ |
| $k$ (character sites) | 28.5 | 29.5 | -0.31 | 0.048* |
| $q$ (collision prob.) | 0.267 | 0.114 | +0.70 | $< 10^{-4**}$ |
| $k/n$ | 0.090 | 0.026 | +0.70 | $< 10^{-4**}$ |
| % missing data | 21.3 | 15.9 | +0.34 | 0.027* |
| Unique alleles/site | 11.7 | 22.9 | -0.71 | $< 10^{-4**}$ |
| % unique cells | 61.8 | 53.8 | +0.13 | 0.43 |
| % unsaturated sites | 93.9 | 100.0 | -0.29 | 0.082 |
| $k \cdot (1 - q)$ | 20.8 | 25.9 | -0.63 | $< 10^{-4**}$ |
| $k \cdot \log(1/q)$ | 37.3 | 61.9 | -0.68 | $< 10^{-4**}$ |

**Table S9:**
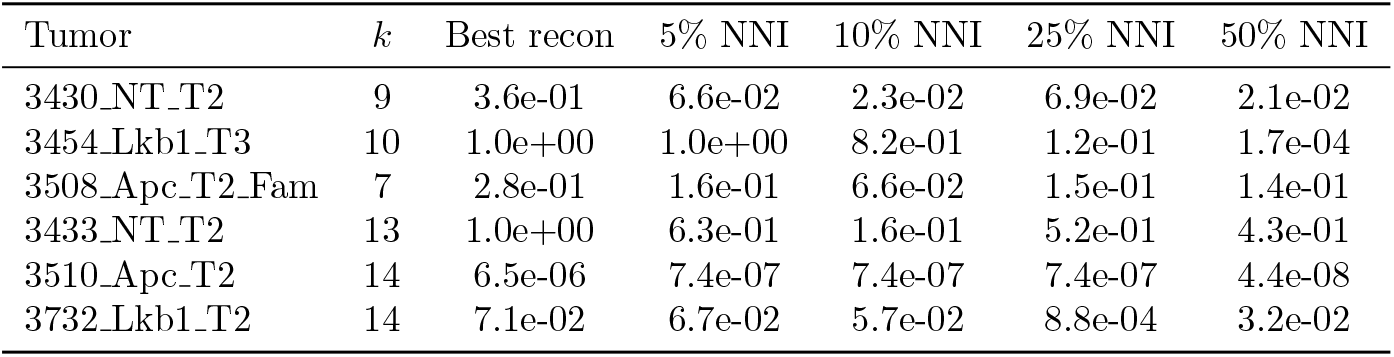
cPHS of perturbed Cassiopeia Greedy trees for the 6 flagged tumors (*k* ≤ 15). Even trees with 25–50% of their topology scrambled by NNI moves routinely achieve cPHS values that would pass any reasonable threshold, confirming that cPHS lacks discriminatory power for these tumors. For comparison, in tumors with *k* ≥ 25 the 50%-perturbed trees almost never pass.

| Tumor | $k$ | Best recon | 5% NNI | 10% NNI | 25% NNI | 50% NNI |
| --- | --- | --- | --- | --- | --- | --- |
| 3430_NT_T2 | 9 | 3.6e-01 | 6.6e-02 | 2.3e-02 | 6.9e-02 | 2.1e-02 |
| 3454_Lkb1_T3 | 10 | 1.0e+00 | 1.0e+00 | 8.2e-01 | 1.2e-01 | 1.7e-04 |
| 3508_Apc_T2_Fam | 7 | 2.8e-01 | 1.6e-01 | 6.6e-02 | 1.5e-01 | 1.4e-01 |
| 3433_NT_T2 | 13 | 1.0e+00 | 6.3e-01 | 1.6e-01 | 5.2e-01 | 4.3e-01 |
| 3510_Apc_T2 | 14 | 6.5e-06 | 7.4e-07 | 7.4e-07 | 7.4e-07 | 4.4e-08 |
| 3732_Lkb1_T2 | 14 | 7.1e-02 | 6.7e-02 | 5.7e-02 | 8.8e-04 | 3.2e-02 |

**Figure S20:**
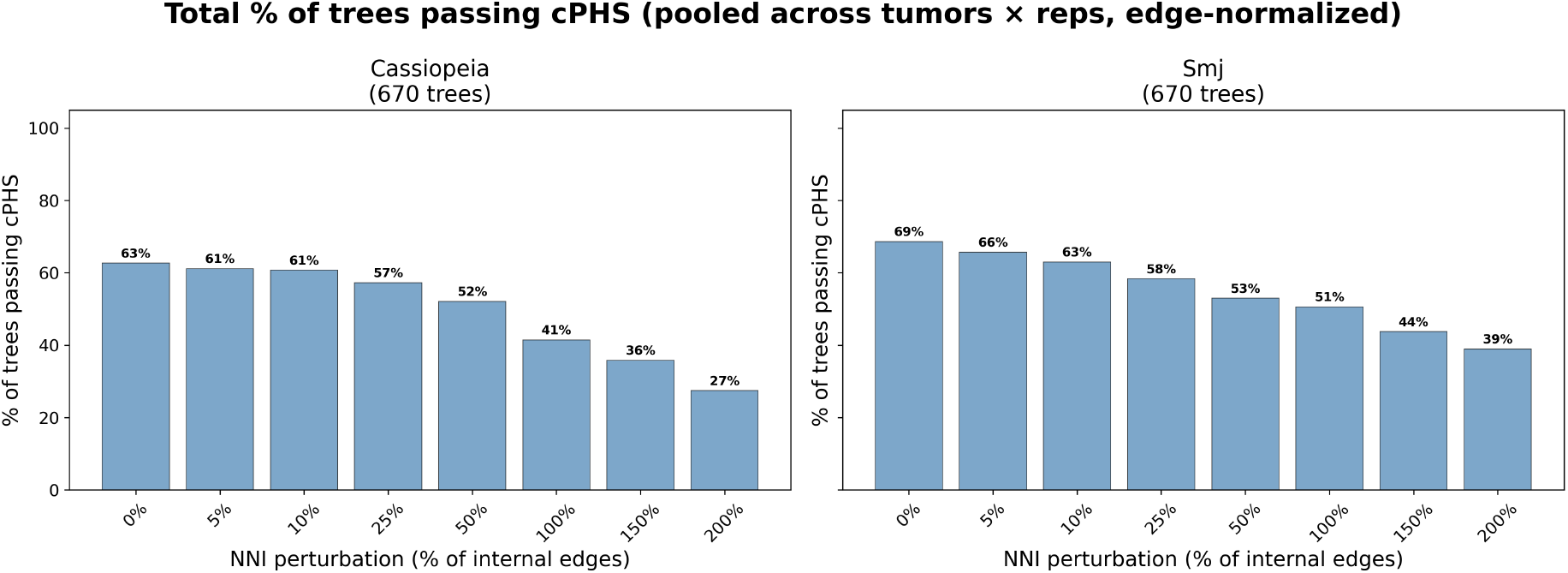
Total percentage of trees passing cPHS at each NNI perturbation level (edge-normalized), pooling all tumors and replicates. Left: Cassiopeia; right: SMJ. Pass rates decline with increasing perturbation.

**Table S10:** Effect of NNI perturbation on reconstruction quality, restricted to the 48 low-*q* tumors (*q* < 0.35) where cPHS is fully reliable. Starting from the Cassiopeia Greedy tree, increasing fractions of NNI moves are applied. All metrics degrade monotonically with perturbation level. cPHS median and mean log_10_ show the typical cPHS value; Pars vs rand and Lik vs rand show the mean percent improvement over random trees.

| Level | $N$ | cPHS pass | cPHS median | Mean $\log_{10}$ | Pars vs rand | Lik vs rand |
| --- | --- | --- | --- | --- | --- | --- |
| Cassiopeia (intact) | 48 | 41.7% | 2.24e-03 | −4.72 | 82.8% | 69% |
| Cass + 5% NNI | 48 | 39.6% | 1.60e-03 | −4.84 | 81.2% | 66% |
| Cass + 10% NNI | 48 | 35.4% | 2.95e-04 | −5.08 | 79.6% | 64% |
| Cass + 25% NNI | 48 | 22.9% | 2.83e-05 | −6.06 | 76.1% | 60% |
| Cass + 50% NNI | 48 | 16.7% | 1.22e-05 | −6.31 | 70.8% | 55% |
| Random | 240 | 0.0% | 5.43e-13 | −10.71 | −0.0% | 0% |

**Figure S21:**
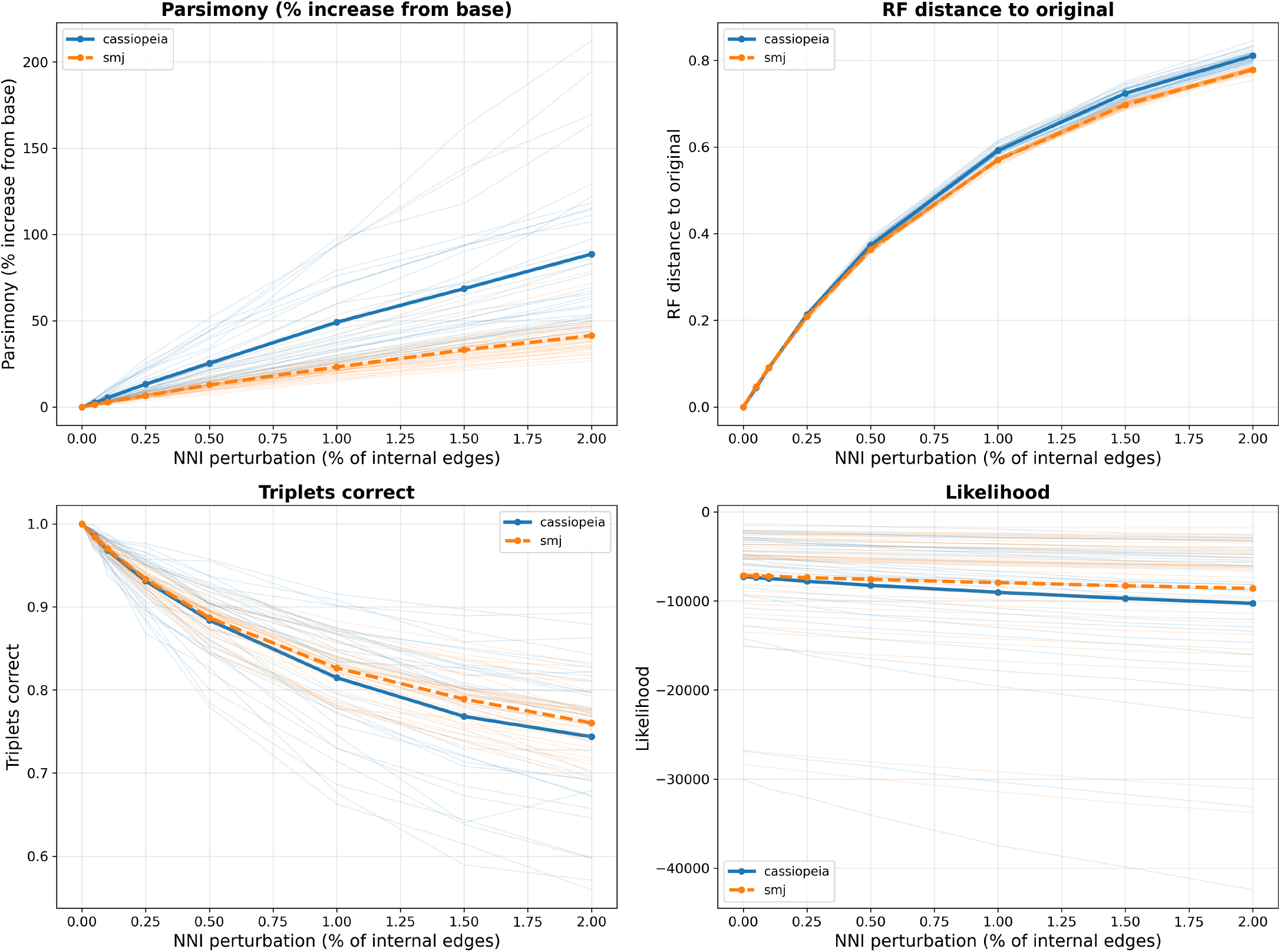
Degradation of quality metrics under edge-normalized NNI perturbation, across all 34 tumors. Top left: parsimony (% increase from base). Top right: RF distance. Bottom left: triplets correct. Bottom right: likelihood. Faint lines: individual tumor means; bold lines: grand mean. Blue: Cassiopeia; orange: SMJ.

## Footnotes

1 A Python implementation of our method will be made publicly available upon publication.

